# Ecology-relevant bacteria drive the evolution of host antimicrobial peptides in Drosophila

**DOI:** 10.1101/2022.12.23.521774

**Authors:** M.A. Hanson, L. Grollmus, B. Lemaitre

## Abstract

Antimicrobial peptides are host-encoded immune effectors that combat pathogens and shape the microbiome in plants and animals. However, little is known about how the host antimicrobial peptide repertoire is adapted to its microbiome. Here we characterize the function and evolution of the *Diptericin* antimicrobial peptide family of Diptera. Using mutations affecting the two *Diptericins* (*Dpt*) of *Drosophila melanogaster*, we reveal the specific role of *DptA* for the pathogen *Providencia rettgeri* and *DptB* for the gut mutualist *Acetobacter*. Strikingly, presence of *DptA-* or *DptB-*like genes across Diptera correlates with the presence of *Providencia* and *Acetobacter* in their environment. Moreover, *DptA-* and *DptB-*like sequence predicts host resistance against infection by these bacteria across the genus *Drosophila*. Our study explains the evolutionary logic behind the bursts of rapid evolution of an antimicrobial peptide family, and reveals how the host immune repertoire adapts to changing microbial environments.

## Introduction

Animals live in the presence of a complex network of microorganisms known as the microbiome. The relationship between host and microbe can vary from mutualist to pathogen, which is often context-dependent (*1*). To ensure presence of beneficial microbes and prevent infection by pathogens, animals produce many innate immune effectors as a front-line defence. Chief among these effectors are antimicrobial peptides (AMPs): small, cationic, host defence peptides that combat invading microbes in plants and animals (*2–5*). While many studies have shown important roles for AMPs in regulating the microbiome (reviewed in Bosch and Zasloff (*6*)), presently, we cannot define why animals have the particular repertoire of AMPs their genome encodes.

Innate immunity has been characterized extensively in *Drosophila* fruit flies (*7, 8*). Antimicrobial peptide responses are particularly well-characterized in this insect (*2, 9, 10*). In *Drosophila*, AMP genes are transcriptionally regulated by the Toll and Imd NF-κB signalling pathways (*8*). Recent work has shown that individual effectors can play prominent roles in defence against specific pathogens (*11–19*). Consistent with this, population genetics studies have highlighted genetic variants in AMPs correlated with susceptibility against specific pathogens. A landmark study in *Drosophila* found that a serine/arginine polymorphism at residue 69 in one of the two fruit fly Diptericins, “S69R” of DptA (Fig. 1A), is associated with increased susceptibility to *Providencia rettgeri* bacterial infection (*20*). Mutant study later showed that flies lacking both *Diptericin* genes (“*Dpt^SK1^*” flies lacking *DptA* and *DptB*) are as susceptible to *P. rettgeri* infection as Imd pathway mutants, while flies collectively lacking five other AMP families nevertheless resist infection as wild-type (*21*). Like these investigations in *Drosophila*, a G49E polymorphism in the AMP Calprotectin of Persian domestic cats is associated with susceptibility to severe “ringworm” fungal skin disease (*22*). Similar AMP variation is common across animals (*23*– *26*). However, while *P. rettgeri* is an opportunistic pathogen of wild flies, and ringworm is common in certain cat breeds, whether these AMPs are evolving to selection imposed by these microbes is unclear. Given recent studies on AMP roles beyond infection (*27–31*), other fitness trade-offs could also explain AMP evolution.

**Figure 1:**
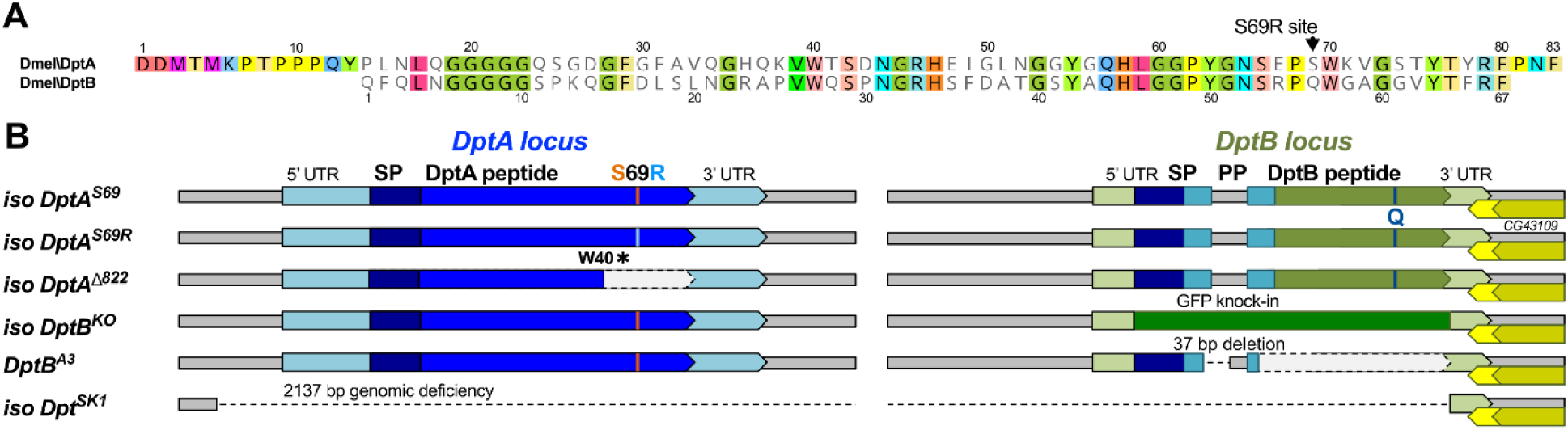
*Diptericins* of *D. melanogaster*. A) Alignment of *D. melanogaster* mature DptA and DptB peptides, which are ∼52% identical. The DptA^S69R^ site is noted (Q in DptB, and see Fig. 1supp1 for protein folding predictions). B) The two *Diptericin* genes are located in tandem on Chromosome 2R;55F with only 1130bp between them. *DptA^Δ822^* encodes a premature stop (W40✱). Strain *DptB^A3^* encodes a 37bp deletion overlapping the *DptB* intron-exon boundary, causing loss of function (Fig. 1supp2). The *Dpt^SK1^* deficiency removes 2137bp deleting the coding region of both genes. *DptB* also encodes a secreted propeptide (PP), similar to *Drosophila* Attacins (Fig. 1supp1, Fig. 1supp3, Fig. 1supp4). “SP” = Signal Peptide.

It is now clear that antimicrobial peptides shape the microbiome (*6*), but defining if or how the host immune repertoire itself is shaped by the microbiome has been challenging. Here we characterize the function and evolution of the *Diptericin* gene family of flies, revealing that these AMPs were selected to control ecologically-relevant microbes.

## Results

### Diptericin B is specifically required for defence against Acetobacter bacteria

*Acetobacter* bacteria are mutualists of *Drosophila* that supplement host nutrition, and are common in wild flies (*32–35*). We previously showed that a strain of A*cetobacter* grows out of control in the gut of *Relish* mutant flies (*Rel^E20^*) lacking Imd pathway activity, or in flies carrying deletions removing 14 AMP genes (*ΔAMP14*) (*36*). Here we identify this *Acetobacter* species as *A. sicerae strain BELCH* (Fig. 2supp1). Gnotobiotic association with *A. sicerae* does not cause mortality, even in *ΔAMP14* flies (Fig. 2supp2). However, pricking flies with a needle contaminated with *A. sicerae* kills *ΔAMP14* flies (*12, 36*), also causing an abdominal bloating phenotype that precedes mortality (shown later). This route of bacterial infection is similar to what flies experience when their cuticle is pierced by natural enemies (e.g. nematodes, wasps, mites (*37–39*)). As *ΔAMP14* flies are killed by *A. sicerae* systemic infection, one or more AMPs are likely required to control opportunistic infections by this microbe. We therefore used flies carrying overlapping sets of AMP mutations (*21*), including a *Diptericin* mutant panel affecting each of the two *Diptericins* (Fig. 1B), to narrow down which AMP(s) protect the fly against *A. sicerae* infection.

Ultimately, deleting just *DptB* fully recapitulates the susceptibility of *ΔAMP14* flies. *Dpt^SK1^*, *DptB^KO^*, and *DptB^A3^* flies (Fig. 2B) suffered 100% mortality after infection, with survival curves mirroring *ΔAMP14* and *Rel^E20^* flies; these *DptB*-deficient flies also present similar levels of abdominal bloating (Fig. 2A-B). Furthermore, ubiquitous RNAi silencing of *DptB* causes both mortality and bloating after *A. sicerae* pricking (Fig. 2supp2B-C). On the other hand, *DptA^S69R^*, *DptA^Δ822^*, and even *ΔAMP8* flies collectively lacking five other AMP gene families (*Drosocin, Attacin, Defensin, Metchnikowin,* and *Drosomycin* (*21*)), resisted infection comparable to wild-type. Finally, *DptB* mutants display increased *A. sicerae* loads pre-empting mortality (Fig. 2C), suggesting a direct role for *DptB* in suppressing *A. sicerae* growth.

**Figure 2:**
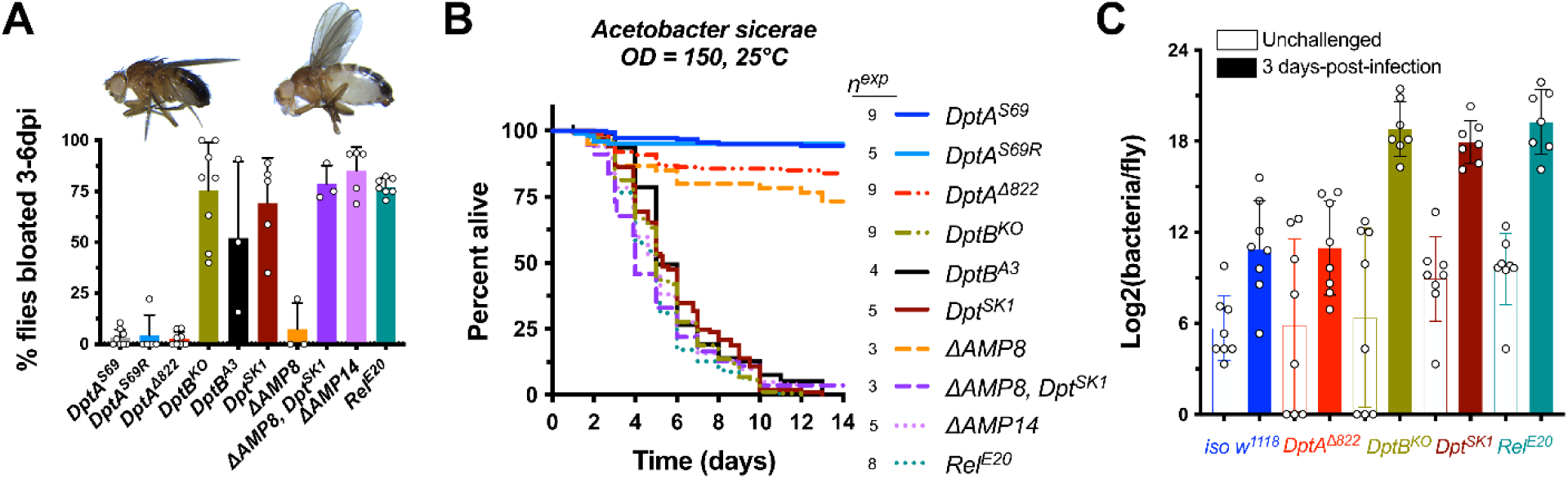
*DptB* is specifically required for defence against *A. sicerae*. A) Flies lacking *DptB* bloat after *A. sicerae* systemic infection. Each data point reflects the average from one replicate experiment (∼20 males). B) Sum survival curves show *DptB* is critical for defence against *A. sicerae*. C) *A. sicerae* bacterial load increases prior to mortality. Each data point reflects the average of 5 pooled flies. *n^exp^* = number of experiments.

After revealing the critical importance of *DptB* in defence against *A. sicerae*, we tested if *DptB* has a broader role in the control of other *Acetobacter* species. To this end, we infected flies with a panel of *Acetobacter* species including *A. aceti*, *A. indonesiensis, A. orientalis, A. tropicalis,* and *A. pomorum*. While these *Acetobacter* species displayed different levels of virulence, *DptB* specifically promotes survival and/or prevents bloating against all virulent *Acetobacter* species (Fig. 2supp3, Fig. 2supp4).

Collectively, these results indicate that *DptB* is an AMP of specific importance in defence against multiple *Acetobacter* species, revealing another example of high specificity between an innate immune effector and a microbe relevant to host ecology. As *Acetobacter* are common in fermenting fruits (*40, 41*), the major ecological niche of *Drosophila*, *DptB* might be especially important for flies to colonize this niche.

### Diptericin A is specifically required to defend against P. rettgeri

The Gram-negative bacterium *P. rettgeri* was isolated from hemolymph of wild-caught flies (*20*), suggesting it is an opportunistic pathogen in *Drosophila*. Previous studies showed that *Diptericins* play a major role in surviving *P. rettgeri* infection (*20, 21*), including a striking correlation between the DptA S69R polymorphism and resistance against this bacterium: flies encoding arginine were more susceptible than flies encoding serine at this site (*20*). However, it is unknown if *DptB* contributes to defence against *P. rettgeri*.

We therefore infected our panel of *Diptericin* mutants by pricking with *P. rettgeri* (Fig. 3A). We confirmed the *DptA^S69R^* allele reduces survival after *P. rettgeri* infection, here with a controlled genetic background (*P* < 2e-16). *DptA^Δ822^* flies also paralleled mortality of *Dpt^SK1^* flies lacking both *Diptericin* genes (*P =* .383). Initially, we found that *DptB^KO^* flies showed higher susceptibility to *P. rettgeri* (*P =* 9.44e-11), correlated with higher bacterial load (Fig. 3supp1A). However, we realized our isogenic *DptB^KO^* flies have only ∼57% induction of the *DptA* gene compared to our isogenic *DptA^S69^* wild-type at 7 hours post-infection (hpi) (Fig. 3supp1B). We therefore confirmed that *DptB^A3^* flies carry the *DptA^S69^* allele, have wild-type *DptA* expression (Fig. 1supp2), and actually survive infection by *P. rettgeri* even better than *DptA^S69^* (Fig. 3A, *P* = 5.03e-4). Moreover, silencing *DptB* by RNAi did not significantly affect survival against *P. rettgeri* (Fig. 3B). We therefore conclude that *DptB* itself does not have a major effect on resistance to *P. rettgeri*, although a cis-genetic background effect found in *DptB^KO^* flies causes lesser induction of *DptA,* and accordingly, higher susceptibility.

**Figure 3:**
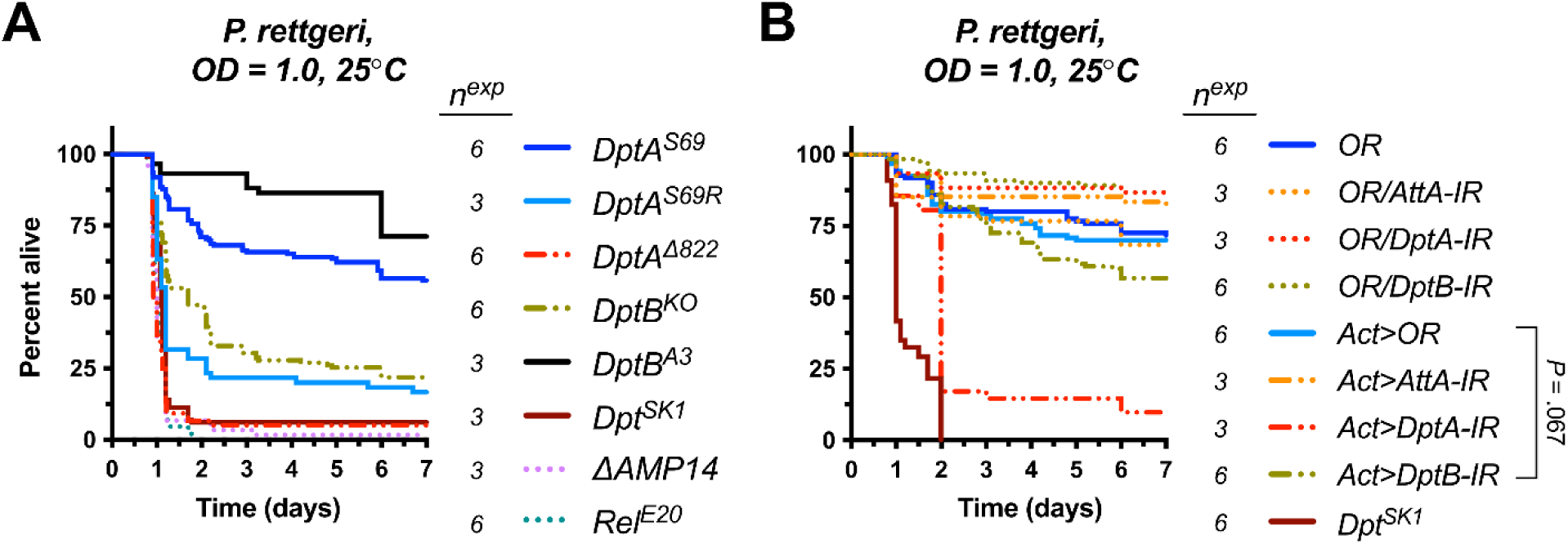
*DptA* is specifically required for defence against *P. rettgeri*. A) Sum survival curves of *Diptericin* mutants after infection with *P. rettgeri*. B) Silencing *DptB* by RNAi *(Act>DptB-IR)* does not significantly affect fly survival compared to *Act>OR* controls.

Taken together, our *Diptericin* mutant panel shows that *DptA* plays a major role in defence against *P. rettgeri,* but not *A. sicerae.* Conversely, *DptB* plays a major role against *A. sicerae*, but not *P. rettgeri*. Thus, these two *Diptericin* genes are highly specific effectors explaining most of the Imd-mediated defence of *D. melanogaster* against systemic infection by either bacterium.

### The Diptericin family shows multiple bursts of rapid evolution across Diptera

Given the high specificity of *D. melanogaster Diptericins* for different ecologically-relevant microbes, we next asked if host ecology might explain *Diptericin* evolution. First, we reviewed the evolutionary history of *Diptericins* across Diptera using newly-available genomic resources (Fig. 4).

**Figure 4:**
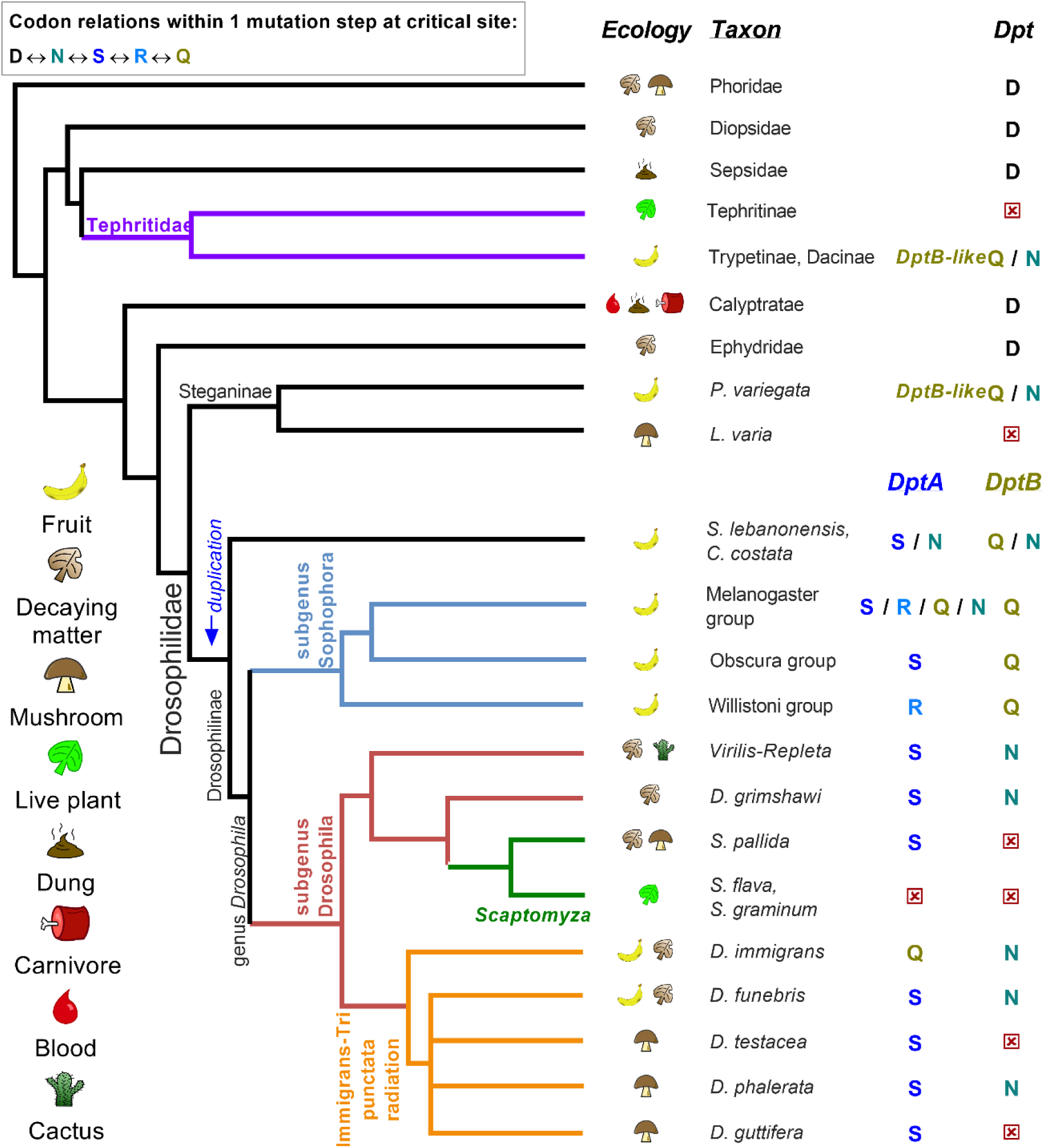
*Diptericin* evolution correlates with host ecology and presence of *Acetobacter* or *Providencia*. *Diptericin* presence was screened in diverse Diptera. The residue aligned to the DptA^S69R^ or DptB^Q56N^ polymorphism is shown. *DptB*-like sequence evolved first in the ancestor of Drosophilidae, and the serine-coding allele in *DptA* evolved at least twice (Fig. 4supp1, Fig. 4supp3). The relatedness of the codons used to encode the S/R/Q/N polymorphism enables their diversification in the subgenus Sophophora (summary in top left). Fruit-feeding tephritids convergently evolved a *DptB*-like gene (Fig. 4supp1, Fig. 4supp2) including a parallel Q/N polymorphism, and *P. variegata* encodes an independent *DptB* duplication, where the two daughter genes encode either version of the Q/N polymorphism. Within Drosophilidae (bottom part), three species with mushroom-feeding ecology have lost their *DptB* genes: *L. varia, D. testacea,* and *D. guttifera*. In both Drosophilinae (*Scaptomyza*) and Tephritidae (Tephritinae), divergence to plant-feeding is also correlated with loss of *Diptericin* genes (Fig. 4supp5). Systematic review of microbiome studies (Fig. 4supp4) suggests absence of *Providencia* and *Acetobacter* in the host ecology is correlated with *DptA* and *DptB* loss respectively. Red [x] = gene loss confirmed. Copy number variation noted in Fig. 4supp3. Phylogenetic cladogram drawn from consensus of (42–45).

*Diptericins* are found across brachyceran fly species, indicating an ancient origin of this antibacterial peptide (>150ma)(*46, 47*). The extant *Drosophila DptB*-like gene was originally derived in the Drosophilidae ancestor through rapid evolution (Fig. 4supp1, Fig. 4supp2, first shown in (*47, 48*)). Later, a duplication of *DptB* gave rise to the *DptA* locus in the Drosophilinae ancestor about ∼50ma (date per (*43*)), which began as a *DptB*-like gene, but then evolved rapidly after the duplication (shown in (*48*) and see Fig. 4supp1, Fig. 4supp2, Fig. 4supp3). Given these repeated bursts of evolution, and only ∼52% similarity between *DptA* and *DptB* (Fig. 1A), distinct antibacterial activities are not necessarily surprising. In reviewing *Diptericin* evolution, we further realized the DptA^S69^ residue of *D. melanogaster* is also present in the subgenus Drosophila via convergent evolution: different codons are used by the subgenus Sophophora (e.g. AGC) and subgenus Drosophila (e.g. TCA) to produce DptA^S69^ residues (Fig. 4supp3), providing further evidence that adaptive evolution selects for serine at this site (complementing (*20, 48*)). Moreover, across species, there is a high level of variation at this site: in addition to the S69R polymorphism, this site can also encode either glutamine (Q) or asparagine (N) in DptA of other *Drosophila* species. Interestingly, Q/N is also seen at the aligned residue of DptB across *Drosophila* species (Q56N in *DptB*). These four residues (S, R, Q, N) are derived compared to the ancestral aspartic acid residue (D) found in most other dipterans (Fig. 4supp3).

Collectively, this analysis suggests that the extant *DptB*-like gene first evolved in the drosophilid ancestor, while *DptA* emerged from a duplication of a *DptB*-like gene, followed by rapid diversification. The DptA^S69^ residue was also derived at least twice, and this site is highly polymorphic across genes and species. These repeated bursts of evolution suggest fly *Diptericins* evolved responding to selection in the drosophilid ancestor.

### Diptericin evolution correlates with microbe presence in host ecology

The diversity of *Drosophila* ecologies, alongside many wild-caught fly microbiome studies, places us in a unique position to pair each host’s microbial ecology with patterns in the evolution of their *Diptericins*, which have microbe-specific importance.

We carried out a systematic review of Diptera microbiome literature (Fig. 4supp4). *Acetobacter* bacteria are regularly found across species feeding on rotting fruits in microbiome studies (*32, 34, 49, 50*). However, *Acetobacter* appear to be absent from rotting mushrooms (*51*), and are largely absent in wild-caught mushroom-feeding flies themselves (*51, 52*). Meanwhile, *Providencia* bacteria related to *P. rettgeri* are common in species feeding on both rotting fruits and mushrooms ((*34*) and Fig. 4supp4). Strikingly, we observed that three drosophilid species with mushroom feeding ecology, *D. testacea*, *D. guttifera*, and *Leucophenga varia*, have independently lost their *DptB* genes (Fig. 4 and (*47*)). Thus, three independent *DptB* loss events have occurred in flies with a mushroom-feeding ecology specifically lacking in *Acetobacter*.

There is another *Drosophila* sublineage whose ecology lacks *Acetobacter*: *Scaptomyza* (Fig. 4 green branch). *Scaptomyza pallida* feeds on decaying leaf matter and mushrooms, while *Scaptomyza flava* and *Scaptomyza graminum* feed on living plant tissue as leaf-mining parasites (*53*). The *S. flava* microbiome shows little prevalence of either *Acetobacter* or *Providencia* (*54*). We checked if these *Scaptomyza* species had pseudogenized either of their copies of *DptA* (two genes, *DptA1* and *DptA2*) or *DptB* (one gene). We found independent premature stop codons in *DptA1* in the leaf-mining species *S. flava* (Q43✱) and *S. graminum* (G85✱), but not the mushroom-feeding *S. pallida* (Fig. 4supp5). We also analysed the promoter regions of these *DptA* genes for presence of Relish NF-κB transcription factor binding sites (“Rel-κB” sites from (*55*), Fig. 4supp5A), confirming only the *S. pallida DptA1* promoter retains Rel-κB sites and likely immune-induction. Thus, *Scaptomyza DptA1* genes show pseudogenization specifically in the leaf-mining species that lack *Providencia* in their present-day ecology. However, *DptA1* appears functional in *S. pallida*, a mushroom-feeding species likely exposed to *Providencia* through its ecology. *Scaptomyza DptA2* genes show variable presence of Rel-κB sites, but no obvious loss-of-function mutations in coding sequence, and *DptA2* remains expressed in *S. flava* (Fig. 4supp5B). Screening the *DptB* genes of *Scaptomyza*, we found no obvious loss-of-function mutations in coding sequences. However, all three *Scaptomyza* species lack Rel-κB sites in their *DptB* promoter regions (Fig. 4 supp5A). Whether due to plant-feeding or mushroom-feeding, none of these *Scaptomyza* have an ecology associated with *Acetobacter*. Using RNA-seq data from the *S. flava* midgut (*56*), we confirmed a lack of expression of both the pseudogene *DptA1* and *DptB* compared to the abundant expression of *DptA2* (Fig. 4supp5B). Taken together, *Scaptomyza* species have independently pseudogenized *DptA* and *DptB* genes correlated with presence or absence of *Providencia* or *Acetobacter* in their ecology.

Finally, convergent evolution towards *DptB*-like sequence has occurred in another lineage of “fruit flies”: Tephritidae (*47, 48*); (see Fig. 4supp1, Fig. 4supp2 for protein alignment and paraphyly of tephritid *Diptericins* clustering with drosophilid DptB). This family of Diptera is distantly related to Drosophilidae (last common ancestor ∼111ma). Many tephritid lineages feed on fruits like *Drosophila* (e.g. Trypetinae, Dacinae), but one lineage parasitizes live plants like *Scaptomyza* (Tephritinae: Fig. 4 purple branches). In light of the present study, the tephritid species that feed on *Acetobacter*-associated fruit (*40, 57, 58*) have convergently evolved a *DptB*-like gene, including a parallel Q/N trans-species polymorphism at the critical Diptericin residue (Fig. 4supp3). Like *Scaptomyza*, plant-parasitizing tephritids lack both *Acetobacter* and *Providencia* in their microbiomes (*47*), and have lost their *Diptericin* genes (Fig. 4 and (*47*)). Thus, *DptB*-like genes evolved in both Tephritidae and Drosophilidae species associated with fruit-feeding ecology where *Acetobacter* is a dominant member of the microbiome. The fact that *DptB*-like genes are not found in species unless their ancestor had a fruit-feeding ecology suggests two things: **1)** the *Acetobacter*-rich fruit-feeding niche was colonized prior to the derivation of *DptB*-like sequence, and **2)** selection imposed by *Acetobacter* resulted in the ancestors of both Tephritidae and Drosophilidae evolving *DptB*-like genes to help control this microbe.

Collectively, our phylogenetic and ecological survey reveals multiple parallels between the host immune effector repertoire, ecology, and the associated microbiome. This suggests that these dipteran species have derived *DptA*- or *DptB*-like genes as their evolutionary solution to control important bacteria found in their microbiome. In contrast, specific *Diptericin* genes become superfluous when their hosts shift to ecologies lacking *Diptericin*-relevant microbes, leading to gene loss.

### Variation in DptA or DptB predicts host resistance across species separated by 50 million years of evolution

Our study indicates that among the suite of immune genes involved in *Drosophila* host defence, the AMPs *DptA* and *DptB* are critically important against two environmentally relevant bacteria: the opportunistic pathogen *P. rettgeri* and the gut mutualist *Acetobacter*. Moreover, our phylogeny-microbiome analysis reveals striking correlations in terms of gene emergence, retention, and loss. If *DptA* and *DptB* really evolved to control *P. rettgeri* and *Acetobacter*, the outcomes of *P. rettgeri* and *A. sicerae* infection across species should be readily predicted using just variation in these two *Diptericins*. We therefore chose 12 *Drosophila* species with variation in the polymorphic site in *DptA* and presence/absence of *DptB*, and infected them with *P. rettgeri* or *A. sicerae*. Of note, experiments in *D. melanogaster* suggest that *DptA^S69R^* affects defence against *P. rettgeri*, but how *DptA^S69Q^* or *DptA^S69N^* affects defence against this bacterium has never been tested. Similarly, the effect of *DptB^Q56N^* on defence is also untested, and so we have no *a priori* expectations for how these polymorphisms affect peptide activity. To analyze these experiments, we used a linear mixed-model approach (see materials and methods), including *D. melanogaster* flies from our *Diptericin* mutant panel as experimental controls. This helped to calibrate our model for the expected effect size for variants of *DptA* or *DptB* within a single species or across species. We also conducted these experiments at 21°C to avoid heat stress to some species, which reduced *D. melanogaster* mortality compared to 25°C (Fig. 5supp1).

Summaries of fly species mortality are shown in Figure 5. As found in *D. melanogaster*, resistance to *P. rettgeri* is associated with a *DptA^S69^* allele across species. Indeed, *DptA^S69R^* found in either *D. melanogaster* or *D. willistoni* correlates with increased susceptibility to *P. rettgeri* (t = −9.59, *P* < 2e-16). *Drosophila yakuba* with *DptA^S69N^* was also more susceptible than its close relatives, suggesting asparagine (N) is an immune-poor allele against *P. rettgeri* (t = −7.26, *P =* 4e-13). Meanwhile, *DptA^S69Q^* flies (*D. suzukii, D. immigrans*) had similar survival after *P. rettgeri* infection compared to *DptA^S69^* flies (t = +0.07, *P =* .35), suggesting glutamine (Q) is a competent defence allele against *P. rettgeri* when coded by *DptA* (Fig. 5A). Overall, ∼74% of variation in susceptibility can be attributed to variation in *DptA* alone as a fixed effect (marginal R^2^ = .743).

**Figure 5:**
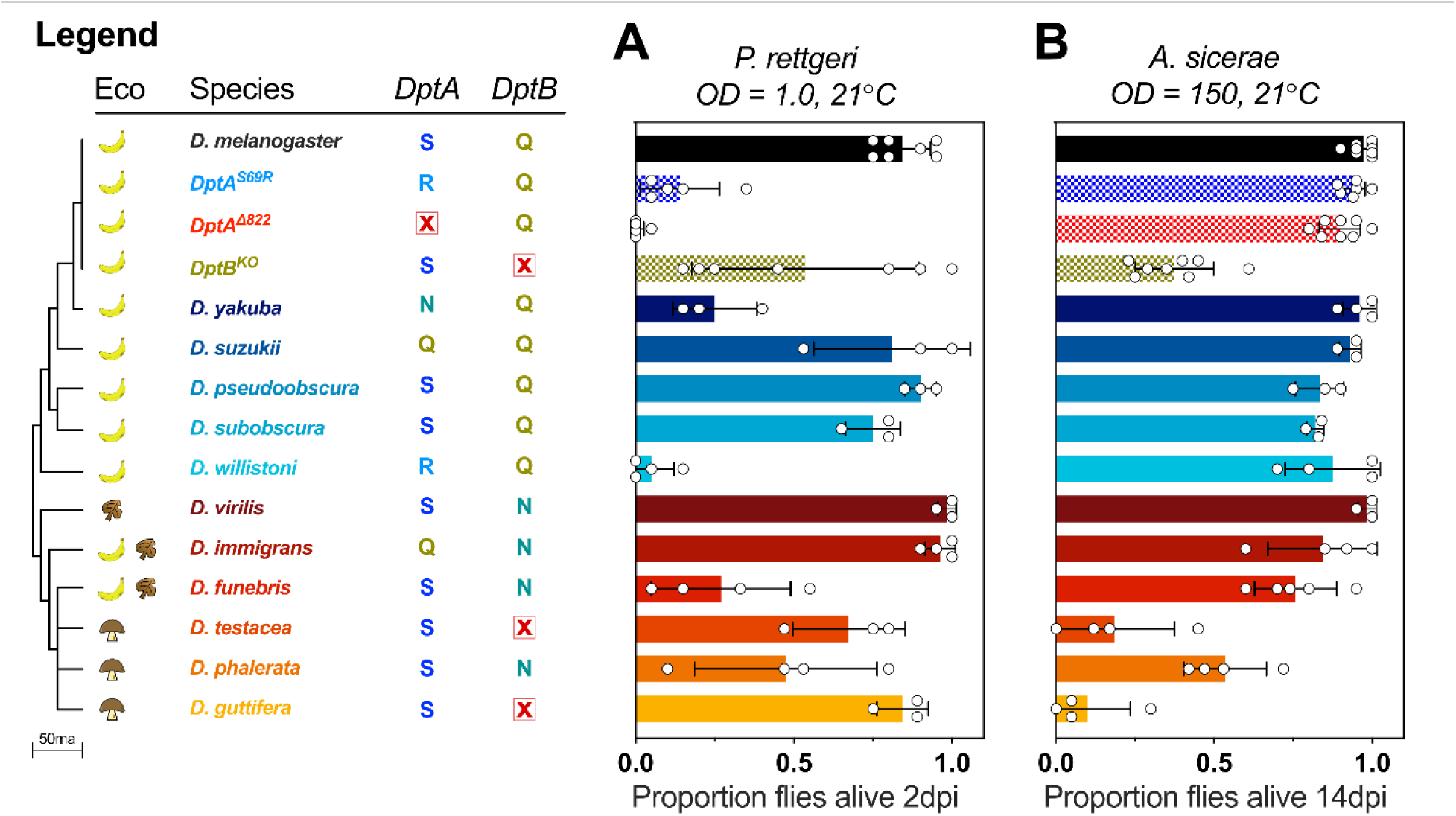
Diptericins predict pathogen-specific survival across *Drosophila*. Host phylogeny, ecology, and *Diptericin* complement shown. Clean injury in Fig. 5supp2. A) Susceptibility to *P. rettgeri* infection varies across species, with survival largely explained by *DptA* allele, particularly within the subgenus Sophophora (blue-shaded species). B) Susceptibility to infection by *A. sicerae* is predicted by presence/absence of *DptB*, though mushroom-feeding flies also had a higher susceptibility to *A. sicerae* infection independent of *DptB* loss. Each data point represents one replicate experiment using 20 male flies.

For infections with *A. sicerae*, absence of *DptB* in the mushroom-feeding species *D. testacea* and *D. guttifera* correlates with increased susceptibility compared to their close relatives (t = −10.83, *P* < 2e-16). Of note, mushroom-feeding flies displayed increased susceptibility to *A. sicerae* infection independent of *DptB* status (t = −3.77, *P =* 2e-4). However, even within this susceptible lineage, *DptB* loss still increases mortality to a similar extent as *DptB* deletion in *D. melanogaster*, indicating the contribution of *DptB* to defence against *A. sicerae* is independent of host genetic background (Fig. 5B). Overall, ∼87% of variation in susceptibility to *A. sicerae* can be explained by just *DptB* absence and host ecology as fixed effects (marginal R^2^ = .868).

These survival data establish that the specific resistance conferred by *Diptericins* observed in *D. melanogaster* applies across *Drosophila* species separated by ∼50 million years of evolution. In conclusion, the host immune repertoire adapts to the presence of ecologically-relevant microbes through evolution of specialized AMPs as weapons to combat specific microbes.

## Discussion

Susceptibility to infection often correlates with host phylogeny (*59, 60*), though host ecology greatly influences microbiome community structure (*34, 61*). Early studies of immune evolution suggested that AMPs were mostly generalist peptides with redundant function, suggesting AMP variation was not due to adaptive evolution (*2, 3*). Instead, studies on immune adaptation have found whole-pathway level effects, or identified factors specific to a given species (e.g. host-symbiont coevolution (*62–64*)). As a result, despite a rich literature on immunity-microbiome interactions, the evolutionary logic explaining why the host genome encodes its particular immune effector repertoire has been difficult to approach experimentally.

Here we identify how ecological microbes promote the rapid evolution of effectors of the immune repertoire, tailoring them to be highly microbe-specific. The two *D. melanogaster Diptericin* genes also provide a textbook example of how gene duplication can promote immune novelty, equipping the host with extra copies of immune tools that can be adapted to specific pathogen pressures. The *Drosophila* Diptericin mechanism of action has been elusive due to technical difficulties in peptide purification (*2, 10*). However studies using *Phormia terranovae* highlight many directions for future research ((*46, 65, 66*) and see supplemental discussion). Future studies combining both fly- and microbe-genetics should be fruitful in learning how host and microbe factors determine specificity. One goal of infection biology is to try to identify risk factors for susceptibility present in individuals and populations. Our study suggests that characterizing the function of single effectors, interpreted through an evolution-microbe-ecology framework, can help explain how and why variation after infection occurs within and between species.

The fly *Diptericin* repertoire reflects the presence of relevant microbes in that species’ ecology. Conversely, loss or pseudogenization of *Diptericins* is observed when the microbes they target are no longer present in their environment. In a sense, this means that some AMPs seen in the genomes of these animals are vestigial: immune genes evolved to fight microbes the extant host rarely encounters (e.g. *DptB* in *D. phalerata*). Indeed, flies that lack *DptB* genes are likely disadvantaged on *Acetobacter*-rich food resources, where the possibility of *Acetobacter* systemic infection poses a constant threat. Thus, loss of this AMP makes recolonization of *Acetobacter*-rich rotting fruits a risky proposition, entrenching the host in its derived ecological niche.

While other mechanisms of defence surely contribute to resistance, *Diptericins* have evolved recurrently as the fly genome’s solution to control specific bacteria. Given our findings, we propose a model of AMP-microbiome evolution that includes gene duplication, sequence convergence, and gene loss, informed by the host ecology and the associated microbiome (Fig. 6). In doing so, we explain one part of why various species have the particular repertoire of AMPs that they do. This ecology-focused model of AMP-microbiome evolution provides a framework to understand how host immune systems rapidly adapt to the suite of microbes associated with a new ecological niche. These findings are likely of broad relevance to immune evolution in other animals.

**Figure 6:**
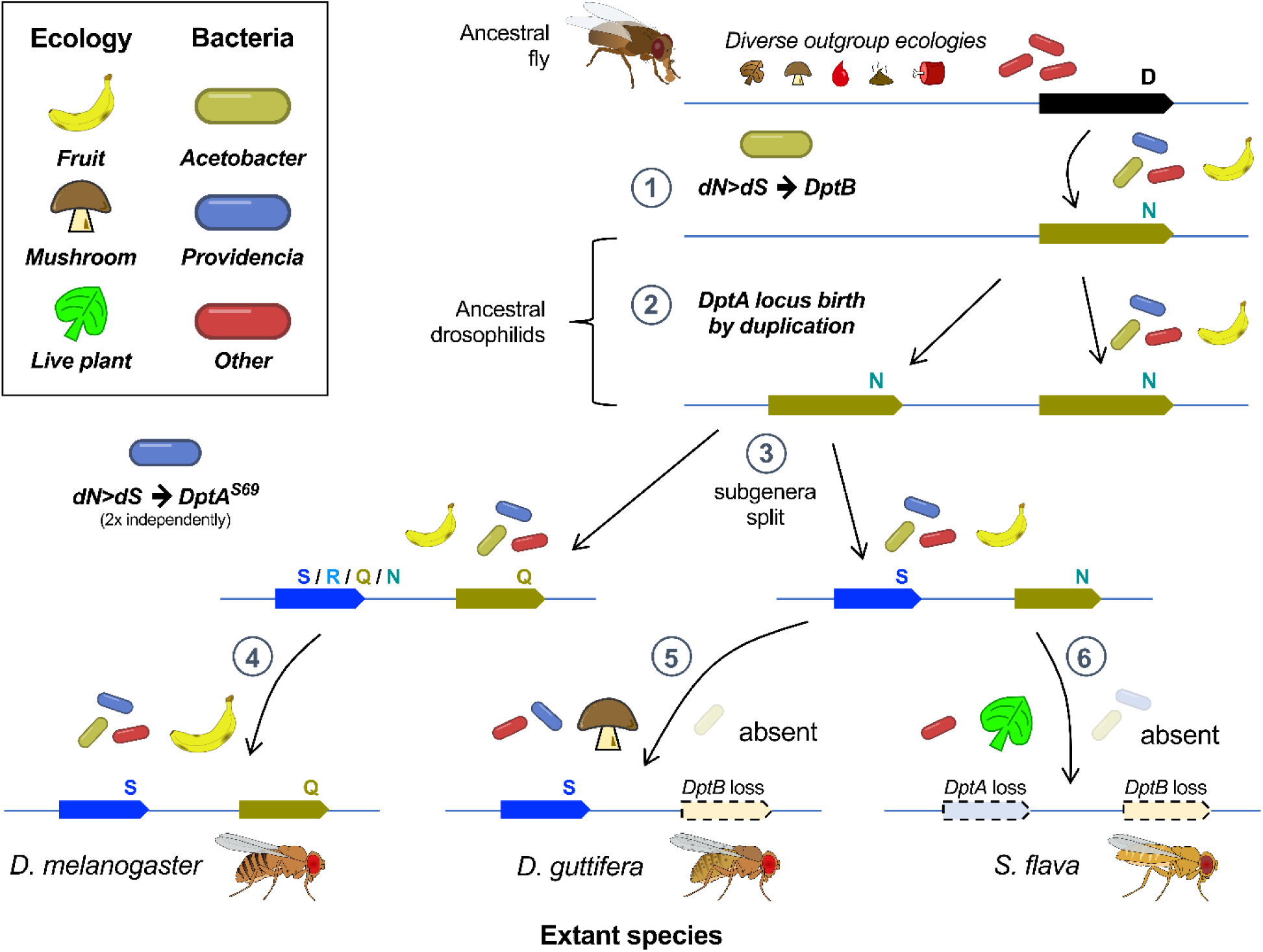
AMP evolution explained using *Diptericins* and a microbial ecology framework. Outgroup ecologies per Fig. 4. **1)** The drosophilid ancestor fed on fruit and was consequently exposed to *Acetobacter* bacteria. *DptB*-like sequence evolved rapidly (dN>dS) to control this novel microbe. **2)** A duplication of *DptB* gave rise to the *DptA* locus. **3)** The addition of a second *Diptericin* gene permitted evolutionary tinkering to control another relevant microbe: *P. rettgeri*, including convergent evolution of the critical S69 residue. In the sublineage including *D. melanogaster*, codon *versatility* enables any of S, R, Q, or N residues. The sublineage including *D. guttifera* and *S. flava* evolved its S residue using a different codon, evolutionarily fixing this residue (Fig. 4supp3). **4)** In *D. melanogaster*, host ecology remains associated with both *Acetobacter* and *Providencia*, which continually select for maintenance of both genes. **5)** In mushroom-feeding *D. guttifera*, *Providencia* remains a threat, but mushroom ecology lacks *Acetobacter*. Consequently, selection is relaxed on *DptB*, leading to pseudogenization. **6)** In leaf-mining species like *S. flava, Acetobacter* and *Providencia* are absent from the microbiome. Consequently, selection is relaxed on both *Diptericin* genes. This AMP-evolution-ecology framework makes sense of why AMPs have microbe-specificity, and helps understand how shifts in microbial ecology can promote rapid evolution for *AMP-microbe* specificity, or loss of “vestigial AMPs” that are relevant *primarily* against microbes the host no longer encounters.

## Acknowledgements

We would like to thank Noah Whiteman’s lab, both Vincent Martinson and Ben Longdon for consultation, Stuart Macdonald for the *DptB^A3^* stock, and JP Boquete for transgenic fly injections. We would also like to thank colleagues for their comments on previous versions of this manuscript. This work was funded by SNSF Postdoc.Mobility grant number P500PB_211082 awarded to Mark Hanson, and Sinergia grant CRSII5_186397 and Novartis Foundation 532114 awarded to Bruno Lemaitre.

## Materials and Methods

Full discussion in supplementary text. **In brief:** *D. melanogaster* fly stocks include both natural mutations, and a transgenic insertion disrupting *DptB*, which were isogenized into the DrosDel isogenic background as indicated in Fig. 1 with the prefix “*iso*”. Non-isogenic *DSPR A3* flies (*DptB^A3^*) are from (*67*). Survival experiments were performed and analyzed as described previously (*21*), with the temperature and OD600 density of the bacteria (“OD”) indicated within figures. 20 male flies were used per experiment unless otherwise indicated, and at least 3 replicate experiments were performed for all data shown in main figures, with raw data available in the supplement. In fly bloating, bacterial load, and gene expression graphs, error bars show standard deviation. The Fig. 4 cladogram and annotations were generated by literature review (Fig. 4supp4), with gene search and annotation methods per (*47*).

The script for Figure 5 is available in the supplement. Briefly, we used a linear mixed-model (“lme4” and “performance” packages in R) with species relatedness and experiment block included as random factors, and variation in *DptA* or *DptB* loci including copy number or alleles at key residues (*D. melanogaster* DptA N52 or S69 alleles) as fixed factors; when loss-of-function was present allele was called as “deleted”. We explored our model both by AIC model selection, and by iterative linear mixed model testing where non-significant fixed factors (e.g. *DptB* allele in explaining survival after *P. rettgeri* infection), and their interactions, were relegated to being random factors in the final model: these two approaches provided similar results, and we use values from linear mixed models in the main text.

## Supplemental discussion

### Roles of AMPs in regulating the microbiome

Many studies have demonstrated the importance of AMPs in shaping the microbiome. In plants, environmental microbes stimulate innate immunity pathways that regulate AMPs, that in turn regulate environmental microbe colonization (*5*). In *Hydra*, expression of specific AMPs correlates with microbiome settlement and composition (*68, 69*). In bobtail squid, AMPs are upregulated upon colonization by *Vibrio* symbionts, helping to develop the squid microbiome (*70, 71*). In oysters, specific AMPs have activity in vitro against certain microbes in specific or synergistic fashions (*72*). In mice, circadian rhythm-dependent expression of AMPs is driven by the microbiome, generating diurnal patterns in resistance to *Salmonella* gut infection (*73*). In insects, disruption of AMPs can lead to otherwise mutualistic bacteria growing out of control (*36, 62*). Despite these many studies, until recently it was presumed AMPs worked as a synergistic cocktail, where each peptide contributed to collectively control microbes. It was therefore surprising to discover that multiple Drosophila AMPs have an extremely disproportionate importance in contributing to defence against certain microbes (*2, 3, 9*). The present study finally provides evidence to inform why AMPs evolved with such high specificity against certain microbes: those microbes are highly important in the ecological niche colonised by the hosts, generating a need to evolve immune effectors suited to dealing with those microbes.

### Mechanisms of immune adaptation have typically taken either a very broad, or very narrow perspective

Previous studies have identified elements of immune evolution associated with ecology or symbiosis. For instance, loss of entire immune pathways has been described for aphids living in ‘clean’ or protected ecologies associated with sap-feeding (*63*). Meanwhile, host coevolution with microbial symbionts can take different forms: in weevils, the AMP coleoptericin-A constrains the growth of their nutritional symbionts (*62*), while in aphids and fruit flies, symbionts themselves can contribute defence against parasites, reducing selection on host immunity (*64*). In *Drosophila,* ecological adaptations for certain food resources may also reduce the risk of parasitism, which again reduces selection on host immunity (*74*). However, in most cases these studies show loss of entire processes (e.g. Imd signalling, genes essential for melanization activity). In the present study, we identify precise bacteria that impose selection on specific host immune effectors. Our study differs as we identify how the immune system of fruit flies has adapted to microbes found in the environment. We found that *DptA* is specifically important to control an opportunistic pathogen of the *Providencia* genus (*P. rettgeri*) found in both rotting fruits and mushrooms, but absent in live-plant ecologies. *Diptericin*s can also be important specifically, or synergistically with other AMPs, against other *Providencia* species (e.g. *P. stuartii, P. burhodogranariea* (*21*)); a specific importance of the Buletin peptide encoded by the *Drosocin* gene was also shown in defence against *P. burhodogranariea* (*12*). We further show the importance of *DptB* in controlling *Acetobacter* bacteria common in fruit ecologies, but absent in mushroom or live-plant ecologies. Using a panel of *Acetobacter*, we show this peptide is protective against infection by many, but not all, of the tested *Acetobacter* strains: of specific interest, *Acetobacter aceti* did not kill wild-type or mutant flies in a marked way, but caused the bloating phenotype in both wild-type and Imd-mutant flies at similar rates. As *A. aceti* is a close relative of the virulent *A. sicerae* (Fig. 2supp1), a focused screen of these bacterial species may reveal bacterial factors that determine *Acetobacter* virulence and the mechanism of bloating that is kept in check by controlling *Acetobacter*, which in most cases, is accomplished by presence of *DptB*.

In the studies discussed above, coevolutionary dynamics are evident: in weevils, the host AMP coleoptericin-A has evolved to control its *Sodalis* bacterial symbionts, which in turn show genome reduction and reliance on their hosts for survival (*75*). A similar process has occurred in aphids, where the Imd immune pathway is lost, alongside the gain of multiple bacterial symbionts that supplement host nutrition, which may have been negatively affected by aberrant immune responses. Aphids also gained other symbionts that surveil the hemolymph and confer defence in place of the generic host immune response (*64*). Our study differs from these examples, as the relationship of *Drosophila* AMPs and ecological microbes is more likely a one-sided evolutionary dynamic, where we show a specific host pattern adapted to environmental microbes, rather than a coevolution of host and microbe.

### Diptericin protein domain discussion

Diptericins are membrane-disrupting peptides, though *D. melanogaster* Diptericins have been technically difficult to purify, preventing their in vitro characterization (*2, 10*). Previous studies with purified Diptericin from *Phormia terranovae* (*46, 65, 66*), have shown that O-glycosylation of threonine in *P. terranovae* Diptericin is critical for antibacterial activity (*65*). Of note, the *P. terranovae* threonine at residue 72 (T72) is universally conserved in *Drosophila* DptB (Dmel\DptB residue T39), but not DptA, which encodes asparagine (Dmel\DptA residue N52) in most subgenus Sophophora species, and is variable in the subgenus Drosophila (Fig. 1A, Fig. 4supp1); asparagine can also be glycosylated (*76, 77*). This difference could help explain the specific importance of *DptB* against *Acetobacter*. This site in *DptA* is highly variable across species: the *D. virilis DptA* uniquely encodes isoleucine at this site (I), *D. willistoni DptA* uniquely encodes tyrosine (Y), and *D. funebris DptA* uniquely encodes glycine (G), while other subgenus Sophophora flies encode asparagine (N) and subgenus Drosophila flies encode alanine (A) (Fig. 4supp1). It’s unclear how any of these variants might contribute to resistance or susceptibility. While the residue aligned to S69R is already sufficient to explain the vast majority of variation in resistance to *P. rettgeri*, the interaction of these two residues could be of interest to future study assessing the mechanism of DptA*-P. rettgeri* specificity, or differences between *DptA* and *DptB* activity. Notably, convergent evolution of *DptB*-like genes in both Tephritidae and Drosophilidae for the Q56N polymorphism suggests the Q56N site plays a key role in *DptB*-*Acetobacter* specificity, and so while the exact differences between *DptA* and *DptB* that mediate microbe-specificity await further investigation, the residue aligned to DptA S69R is nonetheless of clear importance in AMP-microbe specificity in both genes. Ultimately, determining the exact differences between *DptA* and *DptB* activity that mediate microbe-specificity await further investigation.

Ultimately, our results parallel recent descriptions of highly-specific AMP-microbe interactions (*12, 20, 22*), and emphasize their importance by providing an evolutionary perspective.

### Diptericin evolution and naming convention

For simplicity of presentation, the *DptA* clade of the subgenus Sophophora and subgenus Drosophila were not distinguished in the main text. Hanson et al. (*48*) note that *DptA* of the subgenus Drosophila has very low similarity to *DptA* of subgenus Sophophora: ∼40% protein similarity in their analysis comparing consensus sequences (Fig. 4supp1, Fig. 4supp2). The two subgenera, by independent means, nevertheless both encode DptA^S69^ (Fig. 4supp3). However, comparing *DptB* by the same method yields ∼65% similarity between the consensus sequences of the two subgenera. This is reflected in the long branch lengths visible in the Fig. 4supp2 protein tree between DptA of subgenus Sophophora (blue) and DptA of the subgenus Drosophila (red), but relatively short branch lengths between the two subgenera within the DptB clade (olive). Thus it may be expected that differences in DptA among the two subgenera may explain some of the variation seen in survival against *P. rettgeri* infection in Fig. 5. Indeed, there was a high degree of variation in susceptibility to *P. rettgeri* and *A. sicerae* among the subgenus Drosophila species that we sampled. For instance, *D. funebris* with DptA^S69^ has comparable survival to *D. yakuba* with DptA^S69N^. Interestingly, *D. funebris* is the only species across our species panel that encodes glycine (G) at the Dmel\DptA^N52^ site, and glycine is never seen in any Diptericin of any other drosophilid species in broader surveys of Diptericins (*47*). This is not the only unique residue to be found at this site (as mentioned, *D. virilis* uniquely encodes isoleucine (I) in its two *DptA* genes), and so this observation is strictly post-hoc and is not robust on its own. A future study testing this residue’s importance in defence against *P. rettgeri*, and possible role in determining Diptericin activity against *A. sicerae*, could reveal principles of how AMP-microbe specificity can evolve (e.g. presence of glycosylated residues).

## Materials and Methods

### Materials availability statement

All publicly-available reagents used in this study are indicated (e.g. Bloomington Stock #). Primers are listed in supplementary tables below. Fly stocks generated as part of this study are available upon request (dispatched by Bruno Lemaitre).

Supplementary data tables including all experimental raw data, *A. sicerae* BELCH sequencing statistics, and alignments and phylogenies, are given in nested folder format sorted per relevant figure and table at: doi:10.5061/dryad.dz08kps2p.

### Fly and bacteria stocks

The wild-type flies used in this study were *DptA^S69^* (iso *w^1118^* DrosDel) and Oregon-R (*OR-R*). The isogenic *Dpt^SK1^*, *ΔAMP8* (previously *ΔAMPs^+Dpt^*)*, ΔAMP8,Dpt^SK1^*(previously *ΔAMP10*)*, ΔAMP14*, and *Rel^E20^* genotypes were as used previously (*21, 78*). The *DptB^KO^*mutation was generated by Barajas-azpeleta et al. (*31*), and then isogenized into the DrosDel background over seven generations of backcrossing as described in (*79*). *DSPR A3* flies containing the *DptB^A3^* allele were generously provided by Stuart MacDonald. The *DptB* loss-of-function mutation in this strain was first noted by Smith et al. (*80*), who we thank for this useful observation. The two mutations in *DptA* new to this study were isolated from DGRP genotypes, and backcrossed to the DrosDel isogenic background over seven generations. *DptA^S69R^* was isolated from DGRP line 38, and *DptA^Δ822^* was isolated from DGRP line 822. The following lines were used for RNAi experiments: Actin5C-Gal4/CyO-GFP (as used in (*47*)), DptA RNAi (BDSC 53923), DptB RNAi (BDSC 28975), AttA RNAi (BDSC 56904), crossed to *OR-R* for *+/RNAi* controls.

We Sanger sequenced the *Diptericin* loci of our various species to ensure that cryptic variation did not affect our interpretation of the correlations between *Diptericin* variation and susceptibility to infection in Figure 5. Of note, we did recover a Q residue at the DptA polymorphic site in our strain of *D. immigrans* (wild-caught, received from Ben Longdon; Fig. 4supp1), which disagrees with the S residue of the Kari17 reference genome available from Kim et al. (*81*). This codon disagreement is possible via a 1bp deletion and 1bp insertion downstream of two consecutive adenine nucleotides, which replaces S with Q while restoring the correct reading frame compared to the Kari17 reference sequence. This observation provides an example that helps explain how transitions to Q residues at this site are achieved in certain species compared to the SNP-based transition route outlined in Fig. 4 top left, which is only possible in subgenus Sophophora flies (Fig. 4supp3).

Bacteria stocks were as follows: *Acetobacter sicerae strain BELCH* was isolated in Erkosar et al. (*82*) from the shared fly facility of the University of Lausanne (UNIL) and Swiss Federal Institute of Technology Lausanne (EPFL) by Berra Erkosar. We name this strain BELCH for “Berra Erkosar (BE), Lausanne (L), Switzerland (CH)”. See below for strain information. *Acetobacter pomorum strain WJL* was a gift from Won-Jae Lee. The other *Acetobacter* strains were ordered from the Leibniz Institute DSMZ collections: *Acetobacter indonesiensis DSM15552*, *Acetobacter aceti DSM3508*, *Acetobacter tropicalis DSM15551*, *Acetobacter orientalis DSM15550*. *Providencia rettgeri strain Dmel* was a gift from Brian Lazzaro.

### Bacteria culturing and infection experiments

*Acetobacter* species were cultured at 29°C overnight in MRS broth with 25g/L mannitol added, and then pelleted and diluted to a final concentration of OD_600_ = 150 in MRS + mannitol. As our study is among the first to do *Acetobacter* septic injury infections, we will leave the following notes for future research: we performed our *Acetobacter* infection experiments with a consistent OD throughout, but pellets from some strains were quite viscous at OD_600_ = 150. We found OD_600_ = 100 gives similar results in pilot experiments using *A. sicerae BELCH*, and OD_600_ = 100 is far easier to pellet/pipette. Lower doses may also be more appropriate depending on the fly genotype and the temperature flies are kept at: e.g. at OD = 29°C, OD_600_ = 10 *A. sicerae* sufficed to kill 100% of *DptB* mutants during the period of observation in our hands, though at 29°C, more regular flipping is recommended to avoid vial stickiness effects. *Providencia rettgeri* was cultured at 37°C in LB overnight and then pelleted and diluted to a final concentration of OD_600_ = 1.0 in LB.

Septic infections were performed by dipping a 0.2μm needle into the pellet of bacteria, and then piercing the fly cuticle at the junction of the thoracic pteropleura and mesopleura (*83*). Afterwards, flies used in *D. melanogaster*-only experiments were kept on lab standard *D. melanogaster* food. For infections across species, flies were instead kept on Nutri-Fly instant fly medium (Cat# 66-117) resuspended with 0.5% propionic acid in water. Vials were flipped three times per week, and mortality and bloating was recorded daily for *Acetobacter* experiments. Due to inter-experiment variation in onset of bloating and mortality, bloating statistics across experiments are collated to be the last time point prior to onset of mortality to avoid mortality affecting bloating counts. In one experiment (01-03-2023), due to scheduling difficulties, bloating was not recorded. Mortality was recorded twice daily for the first 3 days of *P. rettgeri* experiments, and then daily out to day seven.

### Genome sequencing and phylogenetic placement of *A. sicerae strain BELCH*

*Acetobacter sicerae BELCH* corresponds to previous isolate record *Acetobacter sp. ML04.1*, described as *Acetobacter aceti/Acetobacter nitrogenifigens* by *16s* barcoding previously (*36*). The *A. sicerae BELCH* genome was sequenced by the Microbial Genome Sequence Centre (MiGS) on an Illumina NextSeq 2000 with 2×151bp reads. DNA was extracted from pelleted bacteria using the Illumina DNA Prep kit and IDT 10bp UDI indices. Demultiplexing, quality control and adapter trimming was performed with bcl-convert (v3.9.3). Sequencing statistics are available in supplementary file S1, and the genome SRA is available via GenBank as run SRR21197989.

To construct the *Acetobacter* phylogenetic tree to identify *A. sicerae BELCH*, we collected available sequence data from GenBank and amplified the following genes from DSMZ strains used in this study: *16s, rpoB, GroEL,* and *DnaK*. Primers are listed in supplementary table 1 below. We then concatenated all sequences we could collect/amplify and used PhyML (maximum likelihood, 100 bootstraps) to construct the tree found in Fig. 2supp1 in Geneious R10. This alignment and concatenated tree file are available as file S2, provided in both .nexus and .geneious format.

**Supplementary table 1:**
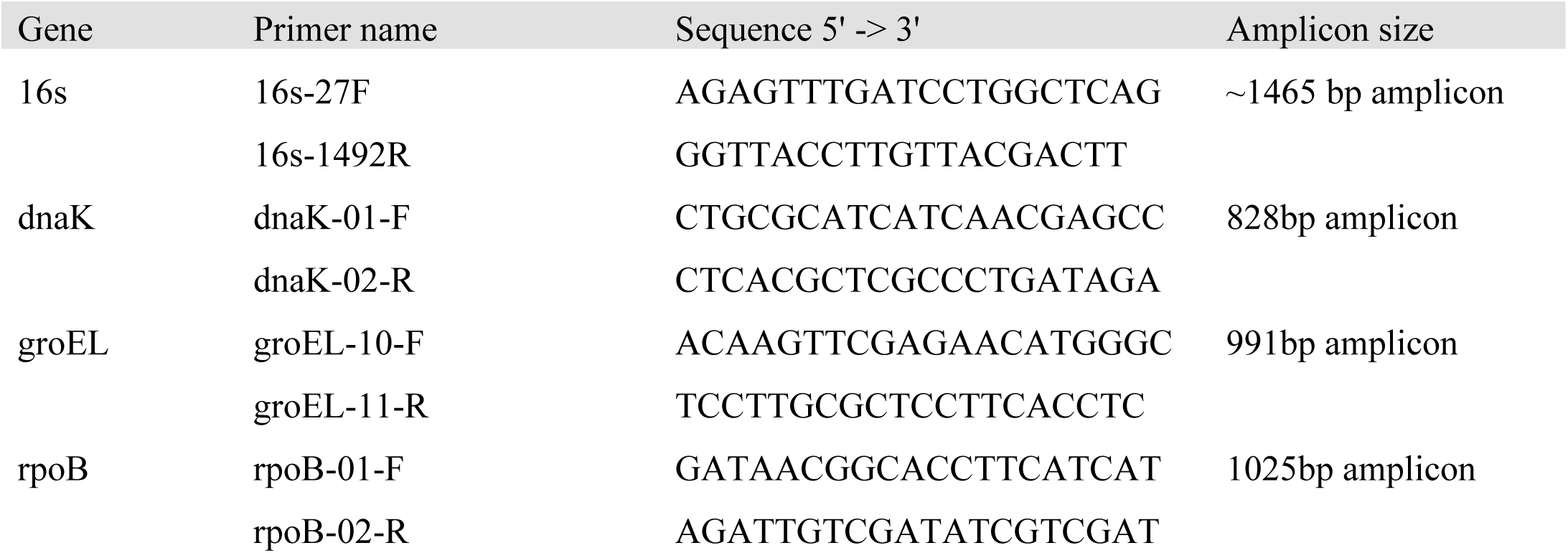
PCR primers used to amplify genes from *Acetobacter* strains for phylogenetic analysis.

### Gene expression analyses

RNA extraction, cDNA synthesis, qPCR experiments and analyses were performed as described in Hanson et al. (*13*). Primers used for qPCR are given in supplementary table 2 below. Of note, the *DSPR A3* strain required a separate set of *DptB* and *Drc* primers to match the *DptB* indel and a *Drc* SNP specific to this strain. As a result, qPCR data for *DSPR A3* (*DptB^A3^*) used a separate set of *DptB* and *Drc* primers from other strains.

**Supplementary table 2:**
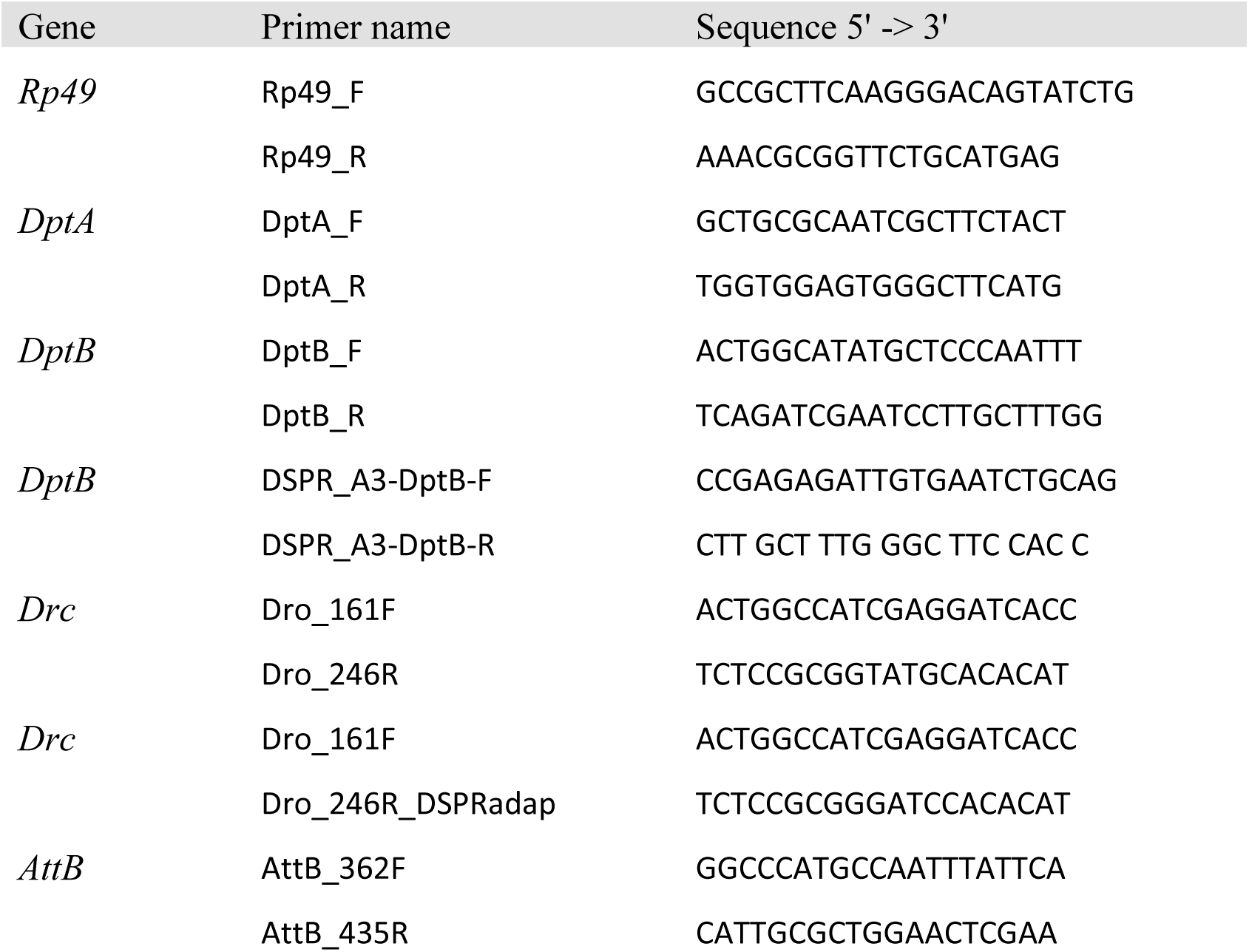
qPCR primers used to measure gene expression.

### Diptericin bioinformatic detection, annotation, and microbiome survey

Antimicrobial peptides are difficult to identify across large phylogenetic distance due to their small size. For this reason, *Diptericin* genes were detected across species using a recursive tBLASTn approach with an artificially high E-value threshold to compensate for the very short query length (E < 100), as used previously (*47*). When no *Diptericin* locus was detected, we instead used tBLASTn to detect the orthologues of longer nearby genes, such as *jheh3* (∼10kb away from *DptA* in *D. melanogaster*), to find the putative *Diptericin* scaffold, then manually scanned the region for conserved *Diptericin* motifs with 1 degree of degeneracy using Geneious R10 (example motifs: “FRF*”, “GGPYG”, “RXRR”), followed by manual curation. If we still failed to detect *Diptericins*, or *Diptericin* pseudogenes that had become degenerate, we made a subset alignment of Diptericin proteins only from the closest relatives and repeated our searches of potential *Diptericin*-containing contigs with 1 degree of degeneracy using whatever motifs appeared common to that lineage (e.g. YRF* in some lineages as the last four protein residues). If we still failed to find recognizable *Diptericin* sequence, the gene was called absent. We could also confirm the origin of the *DptA* locus compared to the independent *Diptericin* duplications of *P. variegata* and *C. costata* (Fig. 4supp1) both by genomic synteny and by virtue of their transcript structure: the *DptA* gene is coded as one exon, while *DptB* and outgroup *Diptericins* are encoded through two exons. The two *DptB*-like genes of *P. variegata* are both encoded as two exons, suggesting an independent *DptB* duplication event. Meanwhile, the *S. lebanonensis* and *C. costata* one-exon *DptA* genes phylogenetically cluster within the *DptB* clade (Fig. 4supp1, 4supp2), confirming that at the inception of this locus, the gene was effectively a second copy of *DptB* that went on to evolve rapidly in both the subgenus Sophophora and subgenus Drosophila.

In all cases within the genus *Drosophila*, we could detect their *Diptericin* pseudogenes directly through premature stop codons or loss of NF-κB binding sites. For *Leucophenga varia* (Drosophilidae) and flies of the Tephritinae subfamily (Tephritidae), no recognizable *Diptericins* were recovered, and we called these genes as absent. In the case of *L. varia*, the *jheh3*-containing chromosome was detected, and no recognizable *Diptericins* were found within 100kb upstream or downstream of *jheh3*. For Tephritinae species, we screened the genomes of *Eutreta diana*, *Tephritis californica*, and *Trupanea jonesi* (*42*), which all lacked *Diptericins* by tBLASTn even using recursive BLAST with all available *Diptericin* sequences collected from Tephritidae and close outgroups.

We surveyed the literature for patterns in microbiome community composition across Drosophilidae and outgroup species. Literature cited is provided in Fig. 4supp4. Chen et al. (*34*) provide a useful summary of multiple microbiome studies in mushroom-feeding, cactophilic, and fruit-feeding drosophilids. Chen et al. also generate microbiome sequencing data for the live plant-feeding *Colocasiomyia* species that lack sequenced genomes, precluding *Colocasiomyia* from our *Diptericin* survey analyses, however *Colocasiomyia* species and their plant hosts both lacked Acetobacteraceae OTUs (*34*). We also surveyed studies of tephritid fruit fly microbiomes, which spanned across both fruit-feeding and live plant-feeding tephritids, finding similar results as found in Drosophilidae for fruit-feeders and *Scaptomyza* plant-feeding species (*57*). Of note, we have presented a simplified view of each species’ ecology for clarity of presentation. For instance, mushroom-feeding likely reflects only part of the ecology of any given mushroom specialist, as these species persist year-round in temperate woodlands even when mushrooms are ephemeral and primarily occur in the fall. Feeding on fungal mycelia or other microbes in decaying vegetation on the woodland floor may act as an alternate niche, which is still poor in sugars and unlikely to act as a hospitable environment for *Acetobacter* sugar/alcohol fermentation to acetic acid. We further contacted Noah Whiteman (University of California Berkeley) for consultation regarding microbiome data from O’Connor et al. (*54*), to confirm low abundance of *Acetobacter* and *Providencia* OTUs in this dataset; *Scaptomyza flava* microbiomes are overwhelmingly dominated by *Pseudomonadales* and *Ricketsialles* OTUs (personal communication). We further thank Andy Gloss (University of Arizona) and Andrew Nelson (University of Arizona) for clarification of the read mapping in *S. flava* midgut RNA-seq data used in Fig. 4supp5B). We also contacted Vincent Martinson (University of New Mexico) for consultation regarding microbiome data of rotting mushroom sites and mushroom-feeding flies from Martinson et al. (*51*) and Bost et al. (*52*), regarding microbiome communities in this ecological niche and in these flies, where *Acetobacter* is largely absent (or when present, only stochastically in certain samplings), while *Providencia* is regularly detected in reasonable abundance (personal communication).

### Statistical analyses

Survival analyses using *D. melanogaster* mutants were analyzed using the survival package and Cox proportional hazard model in R 3.6.3. Experiment blocks and experimenter (two individuals performed infections over the course of the study) were included as covariates, and removed from the final model if not significant. Bacterial load data were analyzed by one-way ANOVA with Dunnet’s multiple test correction in Prism 9.4.1.

Survival analysis across species pertaining to Figure 5 was performed using a linear mixed-model approach using the lmer(), stargazer(), and performance() packages in R.3.6.3. Specifically, an initial model including *DptA* copy number (0, 1, or 2), DptA residue at the S69R polymorphic site (S, R, Q, N, del), DptA residue at site N52 (N, Y, I, A, G, del), *DptB* copy number (0, 1, 2), DptB residue (Q, N, del) at the polymorphic site, and host ecology (fermenting fruit, sap, mushroom) were entered as fixed factors in initial analyses. Host phylogeny was included in the model by nested random effect using an entry of (1|species:species_group:subgenus), and experiment block was also included as a random effect. Fixed effect variables that were non-significant in the initial analysis were relegated to random effect variables in the final analysis, or in the case that these variables were rank-deficient, they were excluded from the final analysis. The response variable was entered as proportion surviving at the relevant time point after most mortality had occurred: 2dpi for *P. rettgeri* and 14dpi for *A. sicerae*.

The script for the statistical model testing correlations in Figure 5 is available in the supplemental data files. We used a linear mixed-model (lmer() function in R) with species relatedness included as random factor using a nested interaction of (1|species:species_group:subgenus). The model included experimental block as a random factor, and included variation in *DptA* or *DptB* loci including copy number or alleles at key residues as fixed factors, with loss-of-function residues included as “allele = deleted”. We explored our model both by Akaike’s Information Criterion (AIC) model selection, and by iterative linear mixed model testing where non-significant fixed factors (e.g. *DptB* allele in explaining survival after *P. rettgeri* infection), and their interactions, were relegated to being random factors in the final model: these two approaches provided similar results. Statistical values reported in the main text come from linear mixed models using only significant fixed factor variables, with other variables being relegated as random factors or dropped due to insufficient sampling (e.g. only *D. willistoni* encoded two *DptB* genes, and so *DptB* copy number was rank-deficient and dropped from final models). Survival of *DptB^KO^* flies was not included in the final model for *P. rettgeri* as the cis-genetic background effect of this mutation causing lesser *DptA* induction (Fig. 3supp2) created spurious comparisons to other Melanogaster-group flies. *DptB^KO^* flies were retained in the model for *A. sicerae*, with *DptA* copy number not entered.

In some cases, variables were insufficiently sampled to draw robust conclusions. For instance, we observed a significant interaction of DptB residue and ecology in survival against *A. sicerae*, but it is difficult to disentangle the nested phylogenetic effect of mushroom-feeding flies (which, when DptB is present, encode DptB^Q56N^) from a genuine impact of the Q56N polymorphism. The inclusion of *D. virilis* in the model also creates spurious comparisons between *D. virilis* and the rest of the species of the subgenus Drosophila, as it was the only species for which a sap-feeding ecology was entered, is one of only two species with two copies of *DptA*, and is a member of a long branch with no close relatives among our species panel. For this reason, we do not report any statistical significance signals where *D. virilis* alone was a major driver of significance (e.g. sap-feeding was significantly different from feeding on fermenting fruits).

We used only a small number of species (twelve), and only tested variation within species using *D. melanogaster*. A future study manipulating *Diptericins* genetically or in vitro, or using a larger sampling of species, will be useful to better resolve the Diptericin mechanism of action and learn how specific residues contribute to defence against *P. rettgeri* or *Acetobacter*.

Raw data and analysis scripts and outputs are available in the supplementary data files included with this manuscript.

## Supplementary figures

**Figure 1supp1:**
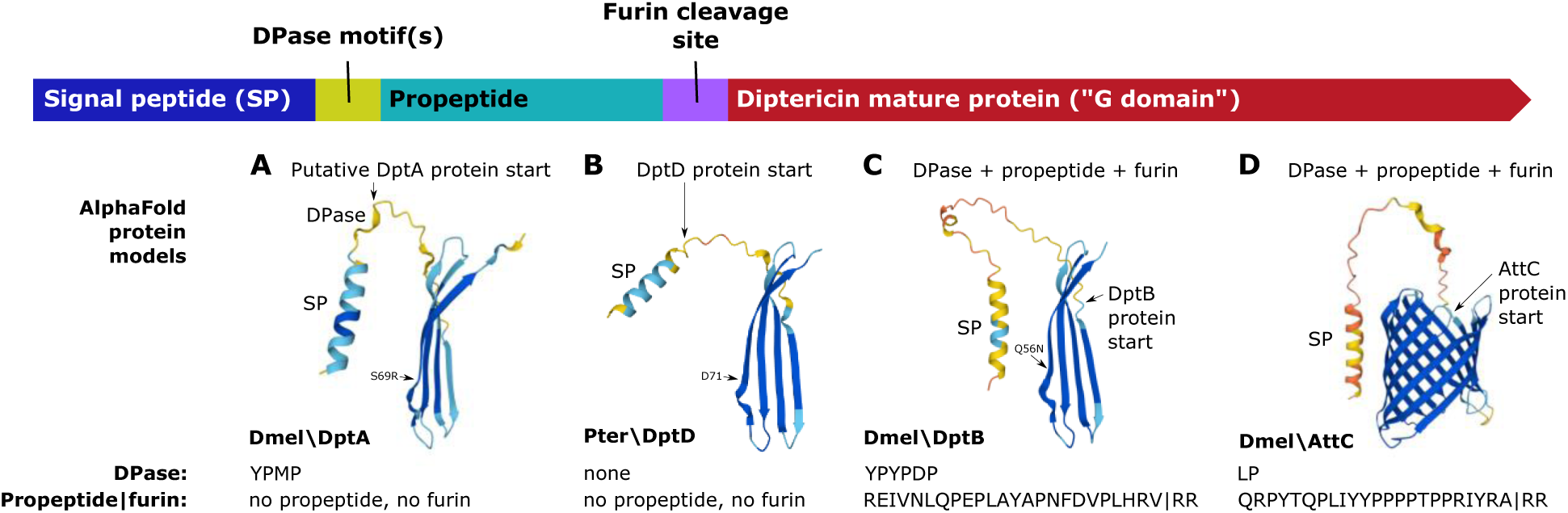
AlphaFold predicted structures for the two *D. melanogaster* Diptericin proteins and comparison to other AMPs. SP = Signal Peptide, DPase = dipeptidyl peptidase motif (“XA” or “XP” motifs highlighted in (*13*)). The residue aligned to the DptA S69R polymorphism is noted for each Diptericin protein, occurring just after a kink in the ß-sheet following a turn. A) DptA has a DPase motif nibbled off the N-terminus of the precursor protein, but no second propeptide. The putative start of the mature DptA protein is indicated with an arrow, corresponding to D^1^ in Fig. 1A. This putative starting residue is supported by isolation of Diptericin beginning at a similar “D” residue as mature protein from *Protophormia terraenovae* (*46, 65*) (B), although this processing has not been confirmed in *D. melanogaster*. C) DptB has tandem repeats of DPase motifs at the N-terminus of its propeptide region, which is a second peptide whose product(s) is secreted into the hemolymph, ending in RV ((*84*) and Fig. 1supp3); “RV” comes from the first two residues of a furin-like cleavage motif (RV|RR, cleavage point indicated as “|”). The putative 67-residue C-terminal Diptericin B protein follows (see Fig. 1A). The arrow pointing to the start of the DptB protein indicates Q^1^ of Fig. 1A, directly following the DptB furin cleavage motif. The ubiquity of furin cleavage in AMP processing (*2*) provides a greater confidence in this being the true mature protein starting residue, particularly as similar N-terminal Q residues are common in other AMPs processed by furin cleavage, also being amidated as part of their maturation (*13*). D) The distantly-related AMP Attacin C (AttC) encodes an antibacterial propeptide called MPAC (*85*), which is preceded by a DPase motif and separated from the mature Attacin pore-forming protein by a furin-like cleavage motif (RA|RR), followed by the protein start indicated by the arrow. The 193-residue AttC protein is much larger than the putative 83-residue DptA protein or the putative 67-residue DptB protein. Importantly, the small size of the Diptericin mature domain suggests that single Diptericin proteins are unlikely to form a pore like the related and larger pore-forming proteins of Attacins. However the predicted concave folding of the Diptericin ß-sheets (A-C), and the Diptericin family’s homology to Attacin family proteins (*86*), suggests they may polymerize, assembling to form toroidal or barrel-stave pores in microbial membranes. We note these AlphaFold models await experimental validation. While Diptericins are membrane-disrupting (*31, 46, 65, 66*), the exact mechanism of action of Diptericins is unknown. We may even speculate the possibility that Diptericins could form hetero-polymers with other AMPs like Attacins to allow or create pores of a different size than any single protein or homo-polymer prediction would suggest, emphasizing the need for experimental validation.

**Figure 1supp2:**
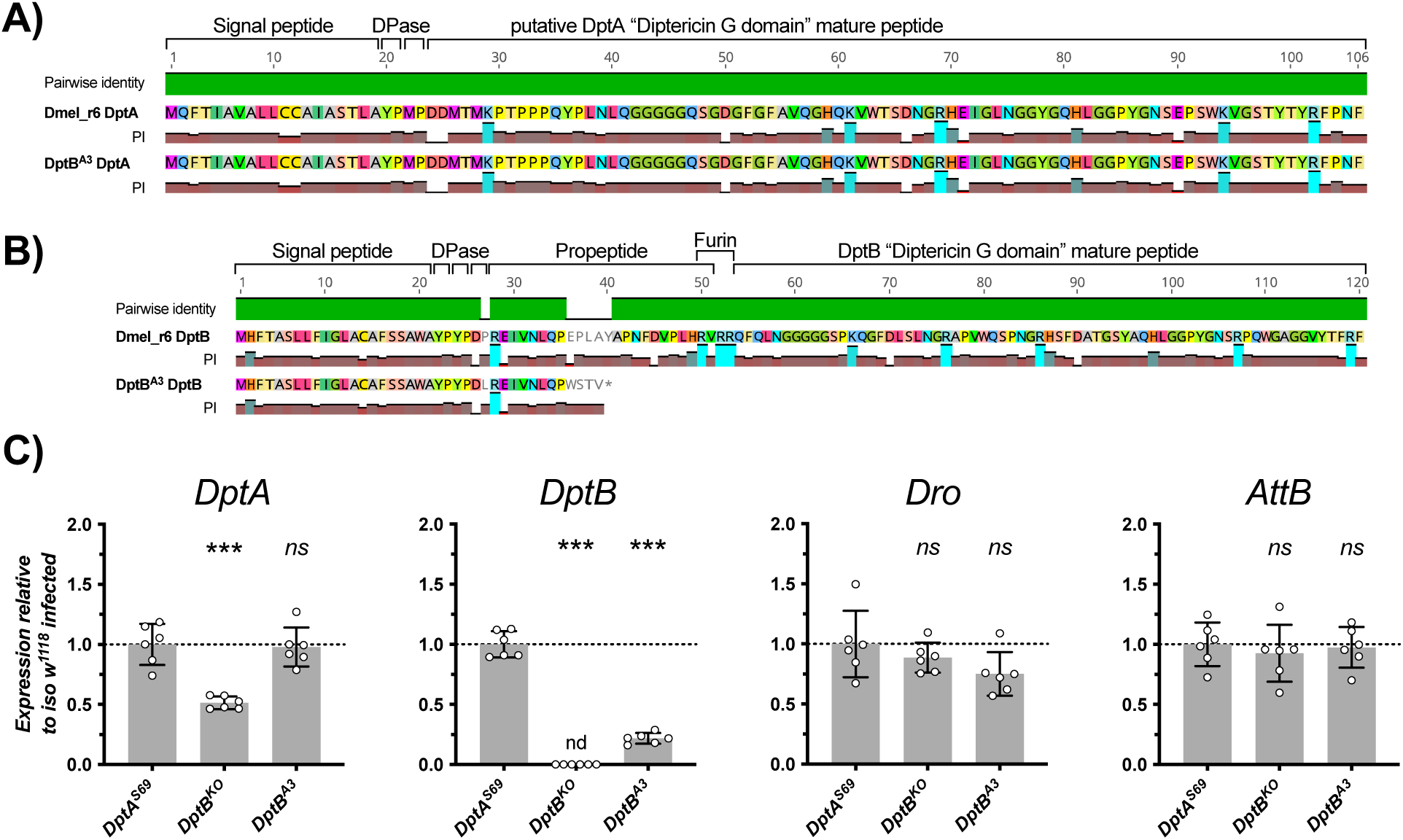
The Drosophila Synthetic Population Resource (DSPR (*67*)) strain *DSPR A3* encodes a natural loss-of-function mutation in *Diptericin B*. A-B) Alignment of strain *DptB^A3^* DptA (A) and DptB (B) proteins to the Dmel_r6 reference genome. Protein domains are annotated (DPase = dipeptidyl peptidase, Furin = furin cleavage site). *DptB^A3^* produces a *DptB* mRNA (C) despite the deletion of 37bp overlapping the intron-exon boundary (Fig. 1B). We confirmed the *Diptericin* sequences of *DptB^A3^* by Sanger sequencing, including the cDNA of the unique *DptB* transcript of *DptB^A3^*, which skips splicing of the intron entirely and matches genomic sequence. As a consequence, there is a frameshift and premature stop present that should cause complete loss of protein function. “PI” = isoelectric point, showing the charged regions of DptA and DptB. Protein domains are annotated, including dipeptidyl peptidase sites (XA/XP, “DPase”) and the furin cleavage motif “RVRR” of DptB that precedes the mature domain beginning in a Q, which is a common structure also found in Attacin and Baramicin where the N-terminal Q is amidated (*10, 13*). C) qPCR of flies 7hpi with *P. rettgeri* confirms a loss of expression of *DptB* in *DSPR A3*, and levels of *DptA, Dro, and AttB* comparable to *DptA^S69^* wild-type levels. *DptB* is not detected (nd) in *DptB^KO^* flies, as expected.

**Figure 1supp3:**
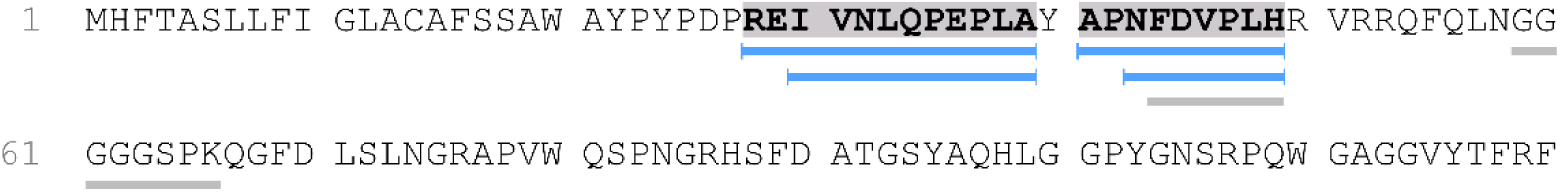
LC-MS data showing detected peptides mapping to the DptB precusor. Hemolymph was collected from *DptA^S69^* wild-type flies 24 hours post-infection by a 1:1 cocktail of OD200 *E. coli* and *M. luteus*, and subjected to trypsin digestion prior to LC-MS performed by the EPFL proteomics core facility. Blue bars show peptides aligned to the DptB propeptide region that were detected in LC-MS with high confidence, matching detection of the same peptides found previously (*84*), although we detected a peptide beginning at “REI”, which matches our prediction of maturated peptide processing given dipeptidyl peptidase sites (XA/XP sites) within the DptB precursor following the signal peptide (Fig. 1supp2B). In contrast, Verleyen et al. (*84*) detected a peptide beginning at “EI”. Grey bars show low-confidence peptides mapping to the DptB precursor. The apparent breakage at the tyrosine (“Y”) between PLA and APN amino acids is not a predicted cleavage of trypsin digestion, suggesting this propeptide region may be maturated into two peptides. Mapping and analysis by “Peaks” software with default settings, performed by the EPFL Proteomics Core Facility.

**Figure 1supp4:**
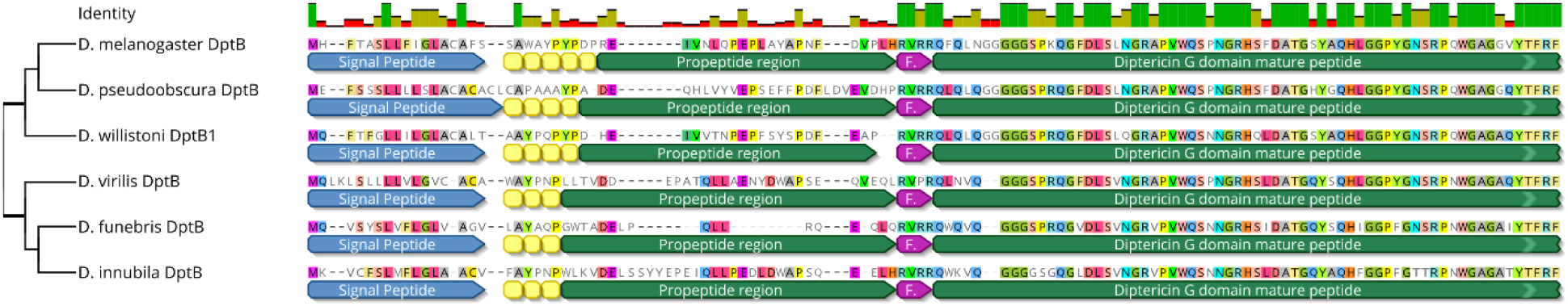
alignment of Diptericin B protein coding sequences. A selection of DptB proteins across diverse *Drosophila* species shows the DptB propeptide region is not conserved across species, while the DptB mature Diptericin domain is highly conserved. Annotations: yellow boxes are “XA” / “XP” dipeptidylpeptidase sites. Purple “F” arrows are RVRR furin-like cleavage motifs.

**Figure 2supp1:**
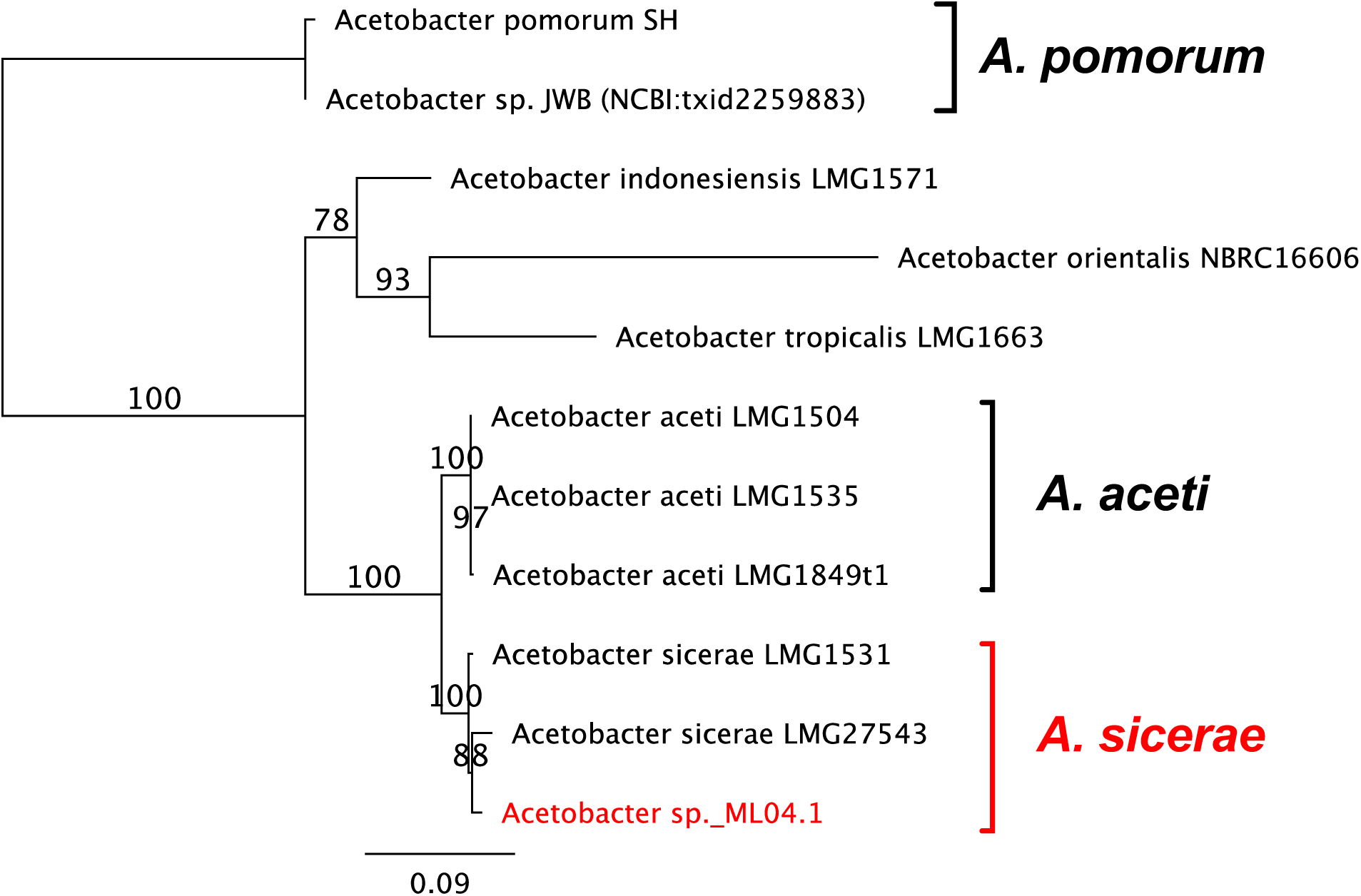
*Acetobacter sp. ML04.1* is a strain of *Acetobacter sicerae*. Maximum likelihood phylogeny (100 bootstraps) made using concatenated sequences of *16S, DnaK, GroEL,* and *RpoB* genes from sequenced strains and Sanger sequences collected for this study. Alignment file available as File S2. We sequenced the genome of *Acetobacter sp. ML04.1*, and designate it *Acetobacter sicerae* strain “BELCH”, for Berra Erkosar (BE) who originally isolated this bacterial species from the food medium of flies reared in the city of Lausanne (L) Switzerland (CH). Sequencing methods and statistics are available in the supplementary materials and methods, and raw sequence reads are available online (GenBank accession: PRJNA873675).

**Figure 2supp2:**
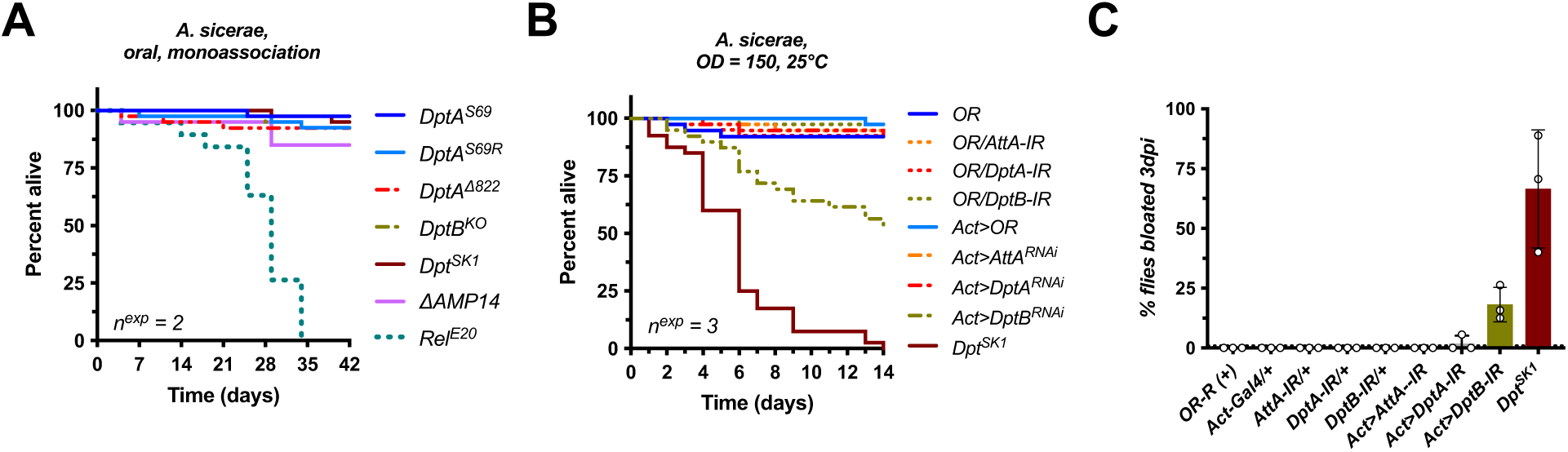
*Acetobacter sicerae* is avirulent upon oral infection, while *DptB* RNAi confirms *DptB* mutant susceptibility to systemic infection. A) *Acetobacter sicerae* is not virulent when administered orally to *DptA^S69^* wild-type*, DptA^S69R^, DptA^Δ822,^ DptB^KO^, ΔAMP14* and *Rel^E20^* flies previously cleared of the microbiota by bleaching, and reared on antibiotic-medium until emergence, whereupon they were given *A. sicerae*-inoculated food. The reduced lifespan of *Rel^E20^* flies is also observed in absence of infection (*87*). B-C) Silencing *DptB* ubiquitously by RNAi (genotype: *Act>DptB-IR*) but not *AttA* or *DptA* induces both susceptibility (B) and the bloating phenotype (C) of *DptB*-deficient flies after systemic infection with *A. sicerae*.

**Figure 2supp3:**
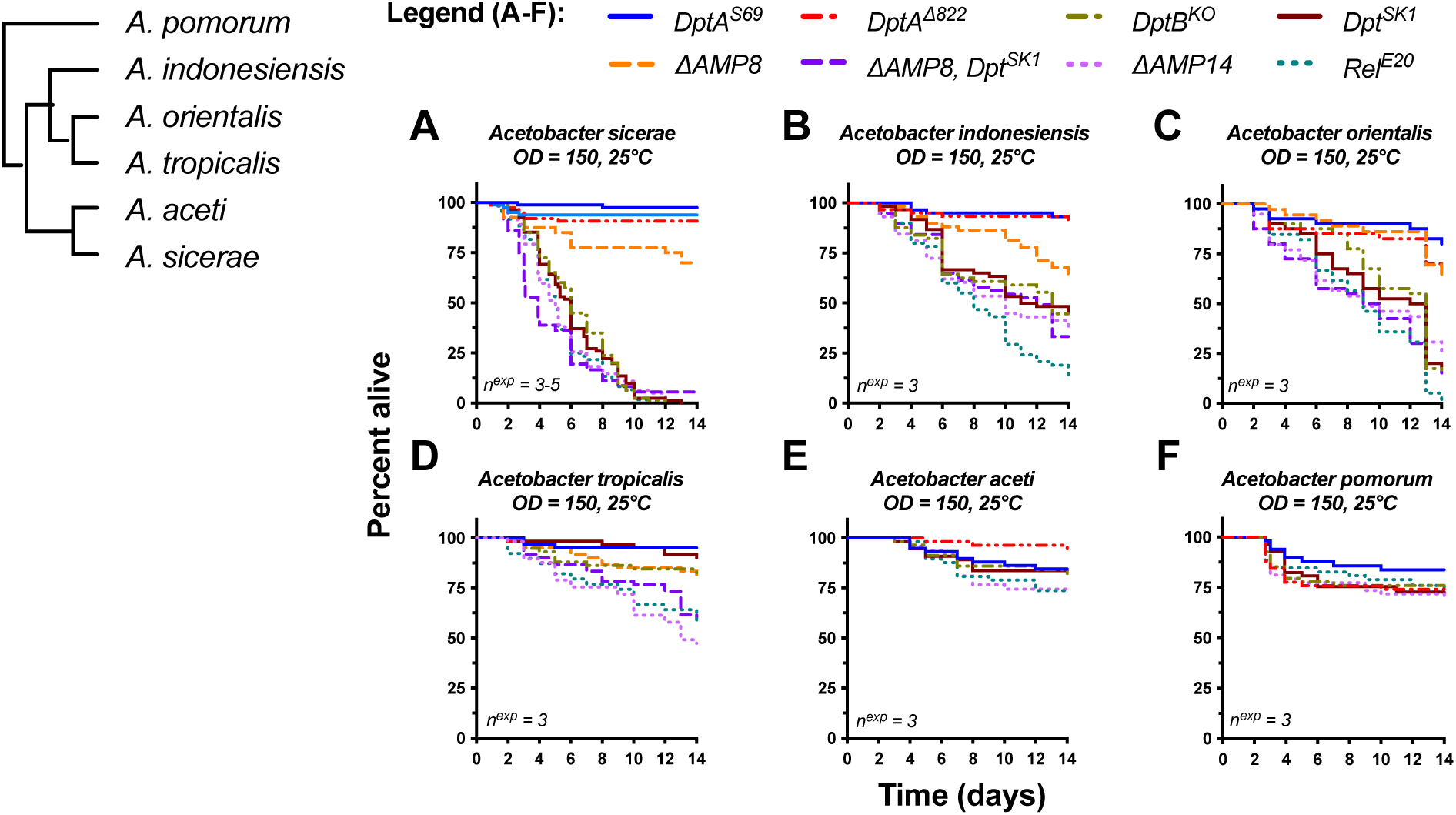
*DptB*-deficient flies display a marked susceptibility to multiple *Acetobacter* species. A-F) Survival of various fly genotypes at 25°C was monitored upon septic injury with multiple *Acetobacter* species. Flies deficient for *DptB* by either *DptB^KO^* or *Dpt^SK1^* mutation (*Dpt^SK1^* present in *ΔAMP14*) were specifically susceptible to infection by *A. sicerae* (A, same experiments as Fig. 2A,B)*, A. indonesiensis* (B), and *A. orientalis* (C). For *A. tropicalis*, both *Diptericins* and other AMPs contribute to resistance, evidenced by survival of *DptA/DptB* mutants, but increased mortality comparing *ΔAMP8* (lacking five AMP families) to *ΔAMP8, Dpt^SK1^* flies also lacking *Diptericins*. Of note, *DptB^KO^* flies, but not *ΔAMP8,* show the *Acetobacter*-induced bloating after *A. tropicalis* infection (Fig. 2supp4D). Infection with *A. aceti* (G) and *A. pomorum* (H) at 25°C did not result in marked mortality, even in *Rel^E20^* flies lacking the Imd pathway. Interestingly, *A. aceti* infection causes bloating in all genotypes tested, with levels comparable even between wild-type *DptA^S69^* and *Rel^E20^* flies. This suggests the bloating phenotype can be induced independent of *DptB-Acetobacter* specificity.

**Figure 2supp4:**
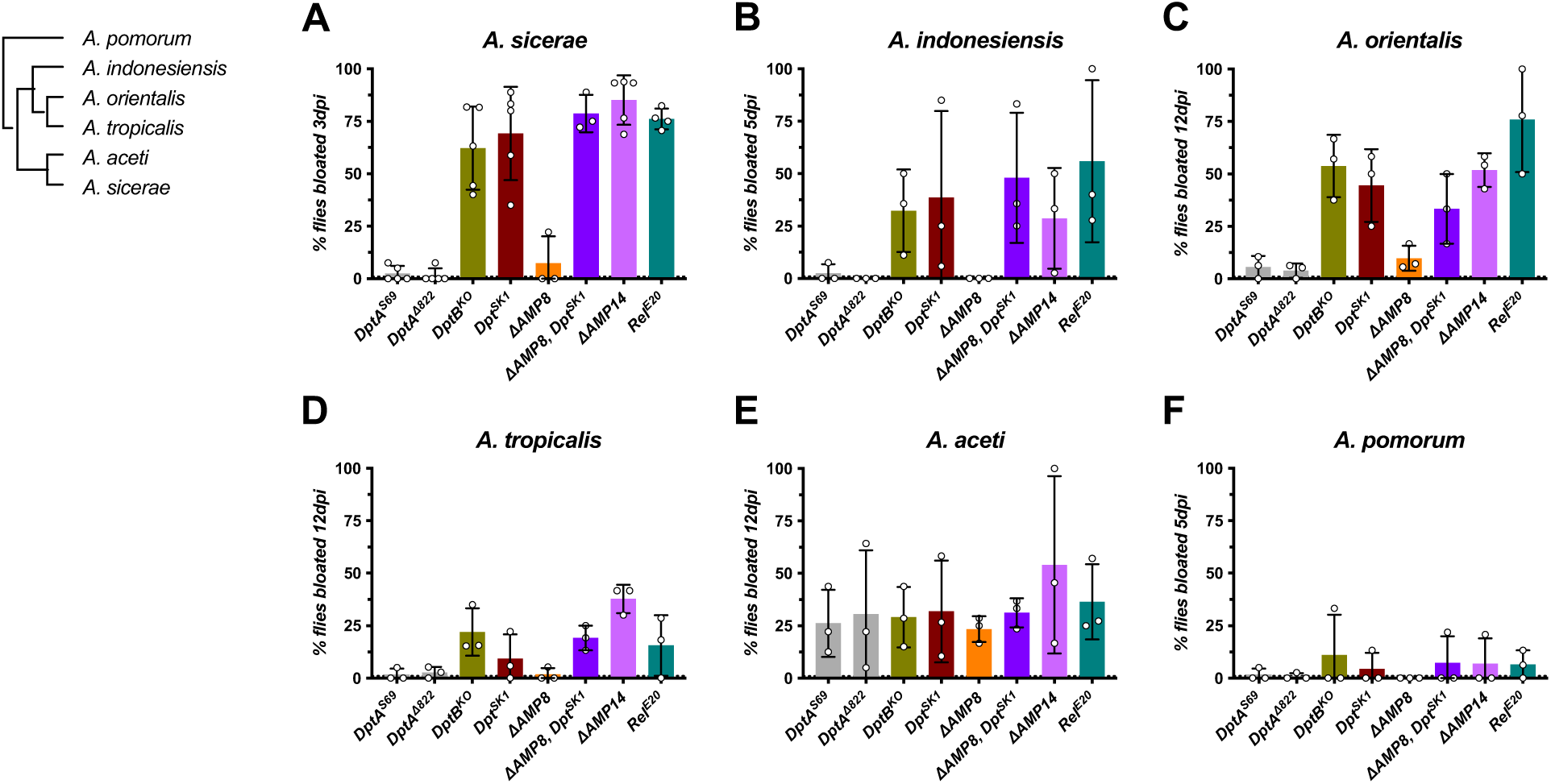
Prevalence of the bloating phenotype in *AMP* deficient flies upon systemic infection with various *Acetobacter* species. The prevalence of the bloating phenotype of various fly genotypes at 25°C was monitored upon septic injury with multiple *Acetobacter* species. Flies lacking *DptB* display a marked increase in the bloating phenotype upon systemic infection with *A. sicerae* (A, same experiments as Fig. 2A,B), *A. indonesiensis* (B), *A. orientalis (C) and A. tropicalis (D).* Meanwhile *A. aceti* infection (E) induces bloating in ∼25% of flies regardless of genotype, although it is relatively avirulent in terms of fly mortality (Fig. 2supp3E). In one of three replicate experiments, bloating was induced by *A. pomorum* in *DptB*-deficient flies while in the two other replicates, bloating was minimal, even in *ΔAMP14* and *Rel^E20^* flies. Bloating was monitored in flies daily, however we only show bloating at the time point prior to onset of major mortality for each *Acetobacter* species, which differ in virulence. We will note here that *DptB*-deficient flies that were not bloated at the time point shown often ultimately became bloated prior to mortality at a later time point for infections with virulent *Acetobacter* species.

**Figure 3supp1:**
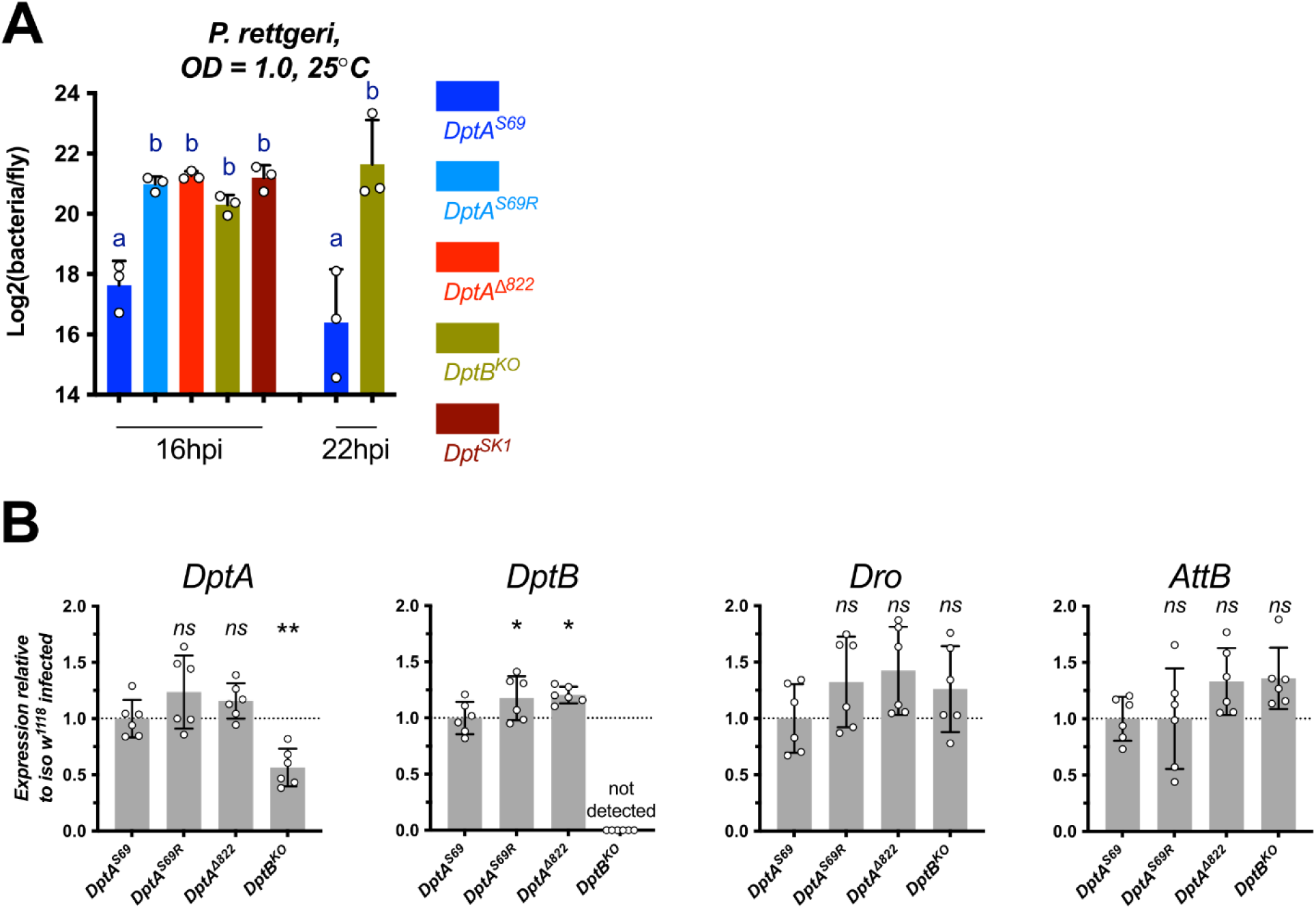
*DptA* expression levels are lower in *DptB^KO^* flies compared to wild-type. The expression of *DptA* is specifically reduced in *DptB^KO^* flies, which is likely due to a cis-genetic effect of the *DptB deficiency* on *DptA* inducibility as another mutation in DptB, *DptB^A3^*, does not cause any susceptibility. The Imd response, evidenced by induction of *Drosocin (Dro)* and *Attacin B* (*AttB)*, is fully-induced in *DptB^KO^* flies. Significance levels: ns = not significant, * = p < .05, ** = p < .01.

**Figure 3supp2:**
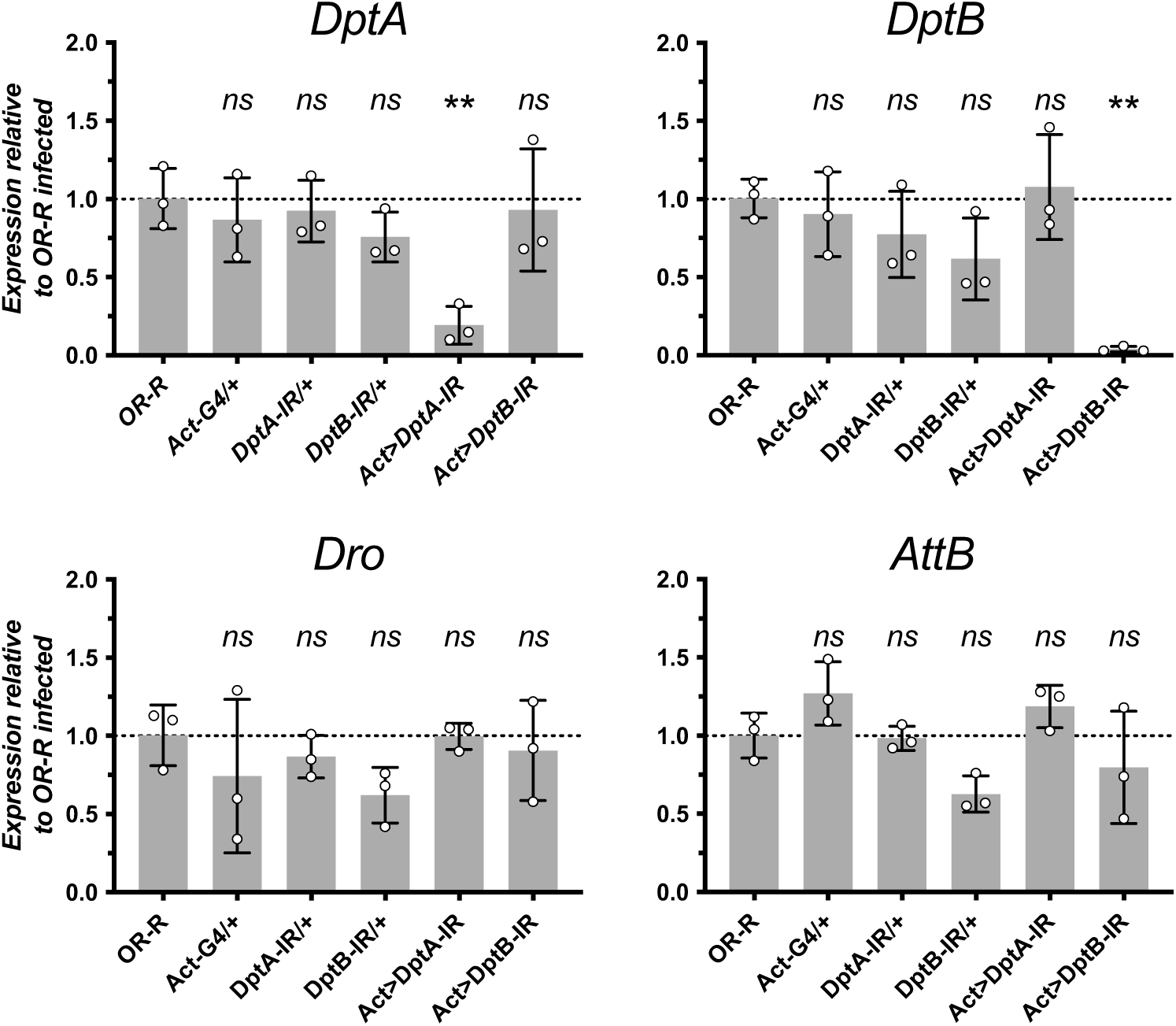
Validation of Imd gene expression in *Diptericin* RNAi (-IR) flies. RT-qPCR at 7hpi in flies infected by *P. rettgeri*, monitoring the expression of *DptA*, *DptB*, *Drososin* (*Dro*) and *Attacin B (AttB)* in flies over-expressing *DptA-IR* and *DptB-IR* confirms gene-specific knockdown. Silencing *DptA* and *DptB* has no effect on the expression of *Dro* and *AttB* indicating that *Diptericin* silencing does not impact the Imd response. Significance levels: ns = not significant, * = p < .05, ** = p < .01.

**Figure 4supp1:**
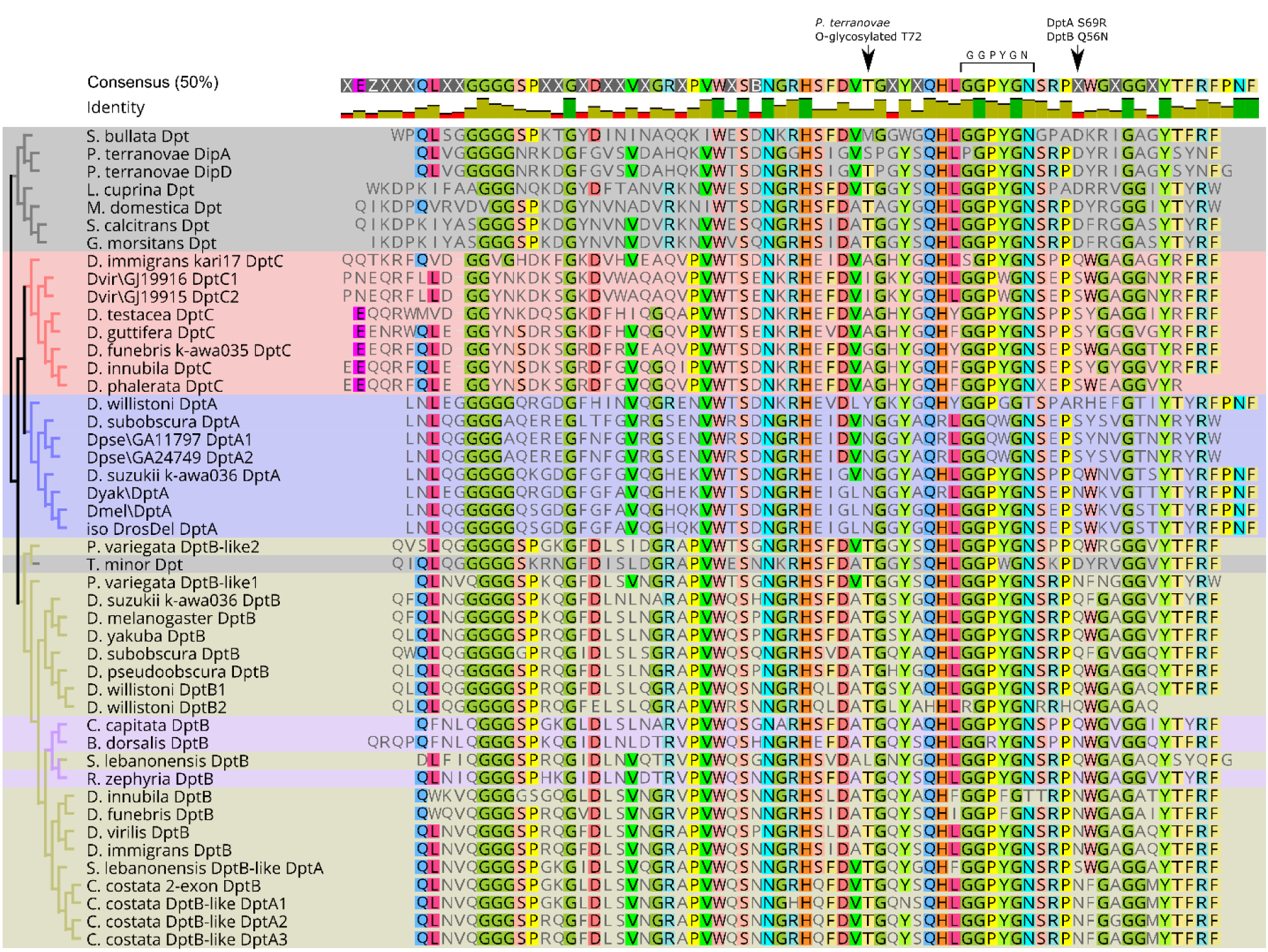
Alignment of the active Diptericin domain (G domain) across species, highlighting key residues in the mature peptide. Blue background = DptA of subgenus Sophophora, Red background = DptA of subgenus Drosophila (“DptC” clade in (*48*)), olive background = *DptB-like* genes of Drosophilidae, purple background = *DptB-like* genes of Tephritidae. Cladogram on left is the result of a neighbor-joining phylogenetic tree (1000 bootstraps, and see Fig. 4supp2). Tephritid Dpt genes cluster paraphyletically within the drosophilid *DptB* clade, indicating convergent evolution towards a DptB-like gene structure more broadly, alongside parallel evolution of the Q/N polymorphism. Of note: *Scaptomyza* species that lack *Relish* binding sites in their *Dpt* gene promoters (Fig. 4supp5) also show unique mutations in highly-conserved motifs: for instance, the universal motif GGPYG<u>N</u> just upstream of the S69R polymorphic site is found across the phylogeny, but has a non-synonymous change to encode GGPYG<u>D</u> in *S. flava* and *S. graminum DptC*, possibly suggesting relaxed selection permitting changes to an otherwise essential Diptericin domain motif. The DptA S69R/Q/N site is noted, as is the site aligned to the O-glycosylated threonine of *P. terranovae* (T72), which is important for its activity (*65*). Of note, this residue differs between DptA and DptB, possibly contributing to their microbe-specific activities. However, this residue is highly variable within DptA/DptC lineages, and so does not readily explain the *DptA* specificity against *P. rettgeri*. Interestingly, *D. funebris* uniquely encodes a glycine (G) at this site, not found in any other *Diptericin* of Diptera, which we confirmed in our stock by Sanger sequencing. In Fig. 5 data, *D. funebris* was more susceptible to *P. rettgeri* infection than any other species encoding DptA^S69^. The two *P. variegata DptB-like* genes are both encoded by 2-exon structures, and are found on different genomic scaffolds. This second *P. variegata DptB-like* gene reflects a separate duplication from the duplication producing the 1-exon DptA locus, which began as a *DptB-like* gene, evidenced by *DptB-like* sequence in the one *S. lebanonensis* and three *C. costata* one-exon *DptA* genes.

**Figure 4supp2:**
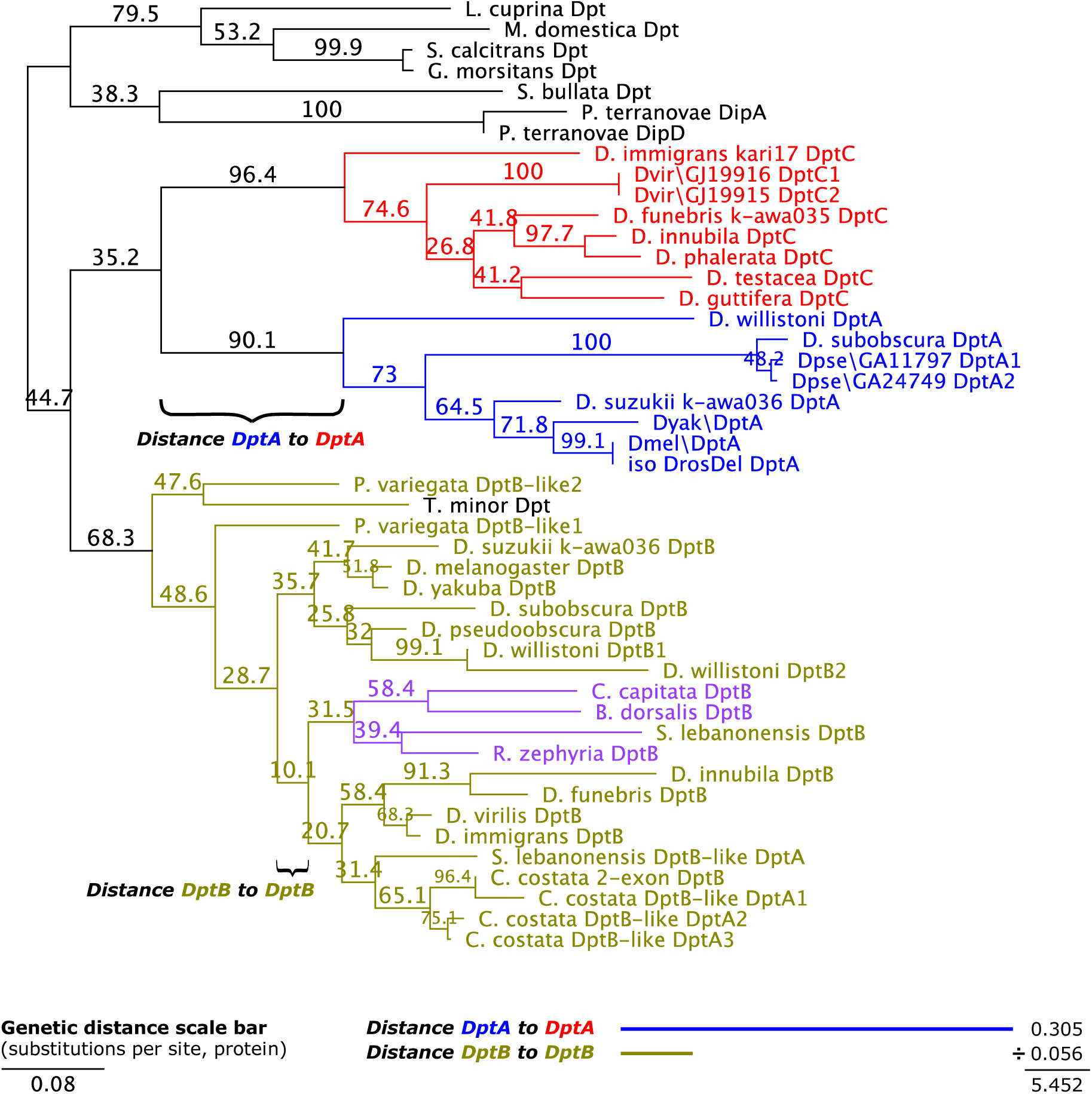
Phylogeny showing rapid evolution of *DptA* locus in the *Drosophila* ancestor. The phylogenetic tree shown here (protein sequence, neighbor-joining, 1000 bootstraps) has branch lengths proportional to genetic distance (substitutions per site), with scale bar indicated in bottom left. In the bottom right, we have drawn the total branch lengths connecting the nodes of the *DptA* clade of subgenus Sophophora (blue) and *DptA* clade of subgenus Drosophila (*“DptC”* genes, red), or the comparable branch within the DptB clade topology. The sum distance of the two branches totals ∼0.305 substitutions per site (protein sequence). The same comparison using the DptB clade gives just 0.056. Thus the genetic distance between the DptA clades of the two subgenera implies this locus has evolved with a rate of substitution of nearly 1 in 3 amino acids over the same timeframe as the DptB clade experienced a substitution rate of just 1 in ∼20 amino acids. The rate of evolution in the DptA clade is ∼5.5 times faster than the same comparison across subgenera using just DptB. This illustration reinforces findings from Hanson et al. (*48*) and Unckless et al. (*20*), which previously found evidence of positive selection (elevated rate of non-synonymous changes) within the *Drosophila Diptericin* lineage.

**Figure 4supp3:**
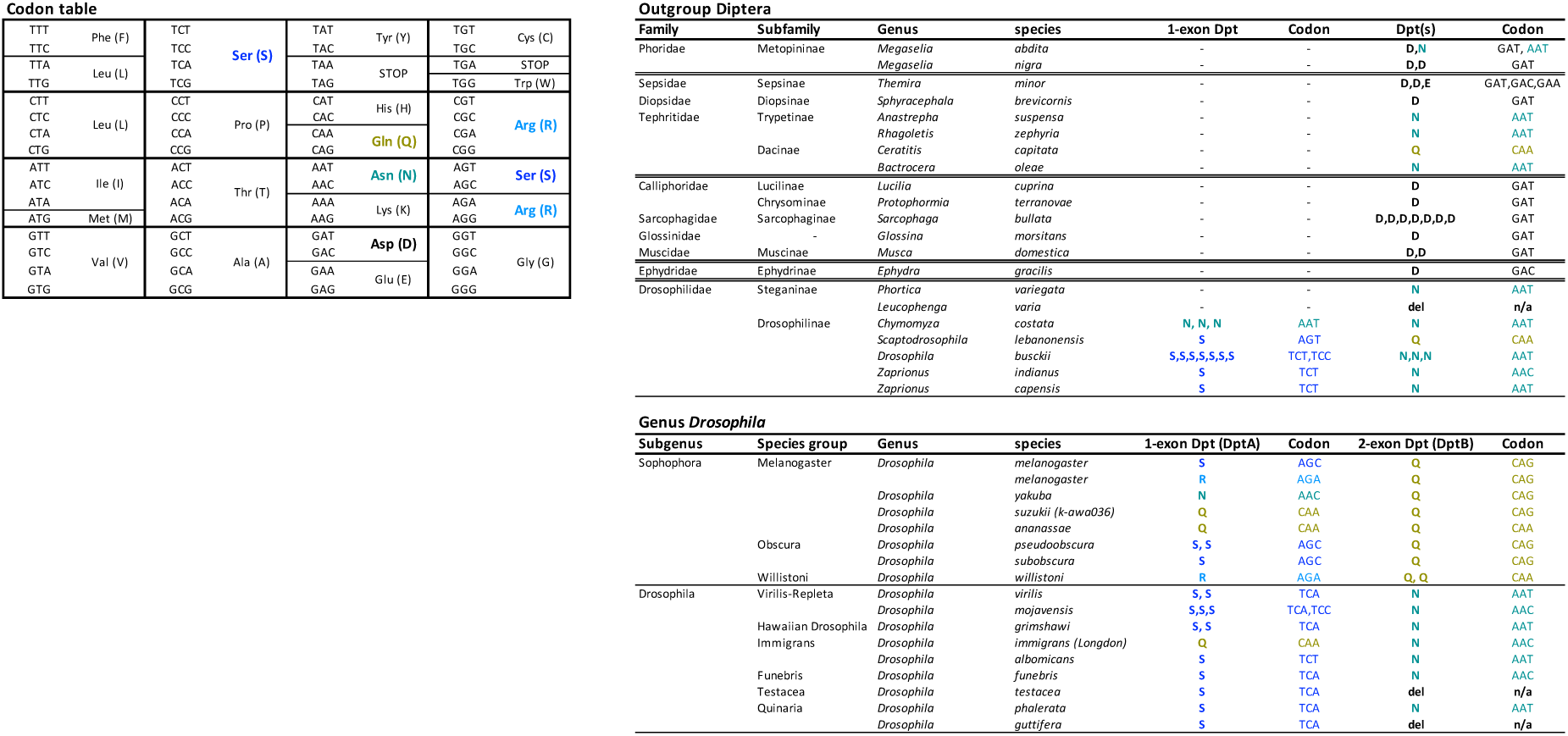
Tracing the evolution of the Diptericin S69R/Q/N/D polymorphic site across Diptera. The ancestral D-encoding codon of Diptericin was GAT, which enables a one-step mutation to AAT encoding N. This AAT codon is likely the ancestral codon in both *DptB*-like genes of Tephritidae and Drosophilidae. The subgenus Sophohora serine is encoded by AGC, suggesting a series of mutations such as AAT>AGT>AGC may have occurred in the ancestor of Sophophora to produce the serine codon. However, the subgenus Drosophila serine is encoded by an ancestral TCA codon, indicating it emerged by convergent evolution through a separate genetic event. Moreover, the Sophophora AGC codon is one mutational step away from either asparagine or arginine, whereas the subgenus Drosophila TCA codon requires a minimum of 2 mutations to reach any other common polymorphism residue. Consequently, little variation is seen in the subgenus Drosophila 1-exon *DptA* locus gene, whereas a high degree of variation is seen in the subgenus Sophophora 1-exon *DptA* between serine and arginine (e.g. *D. melanogaster, D. willistoni*), asparagine (e.g. *D. yakuba*), and glutamine (*D. suzukii, D. ananassae*). Multiple residues/codons for a given species’ gene are given by showing residues separated by commas. These indicate copy number variation and give the residue for individual gene copies.

**Figure 4supp4:**
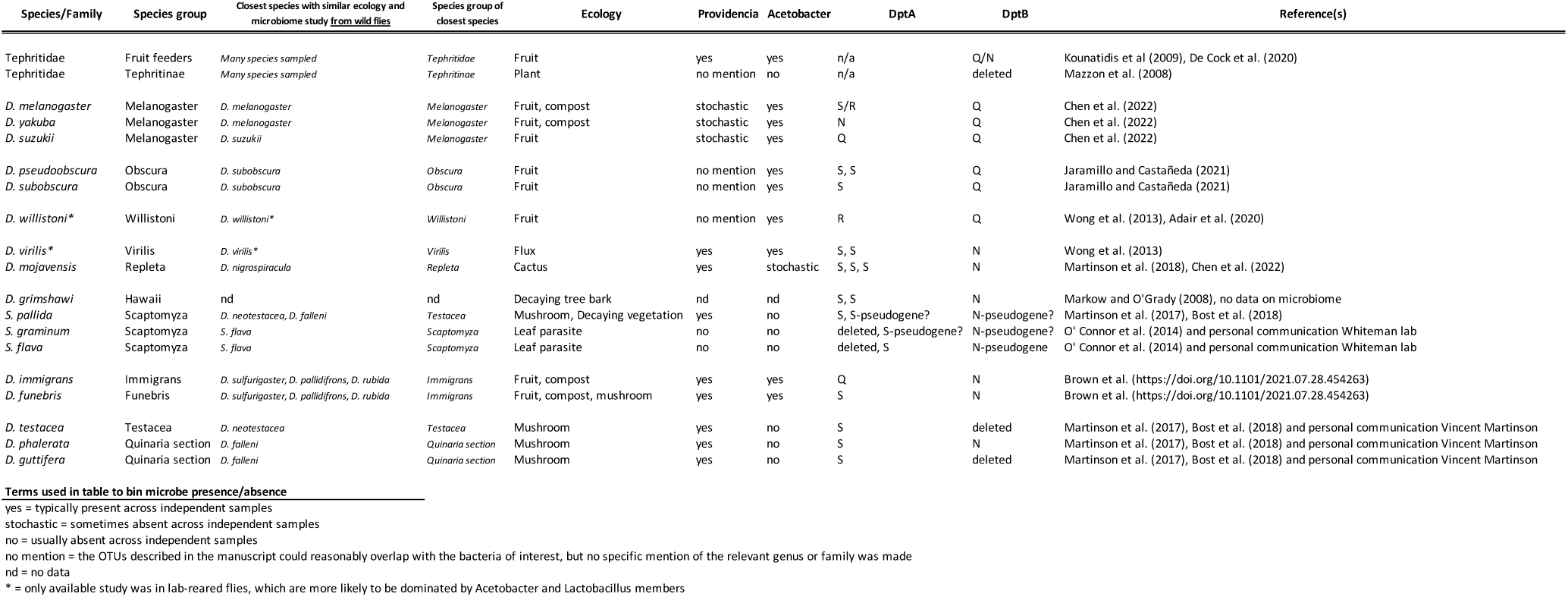
summary of microbiome studies informing interpretation of Figure 4. Species are listed by phylogeny. Microbe presence/absence is annotated according to terms outlined at bottom. Both the species itself, and the closest-related species with a microbiome study are given as some inferences are made by using microbiome studies in close relatives, whereas *DptA/DptB* evolution is screened specifically in species with sequenced genomes. Copy number variation indicated by commas in Diptericin residue columns.

**Figure 4supp5:**
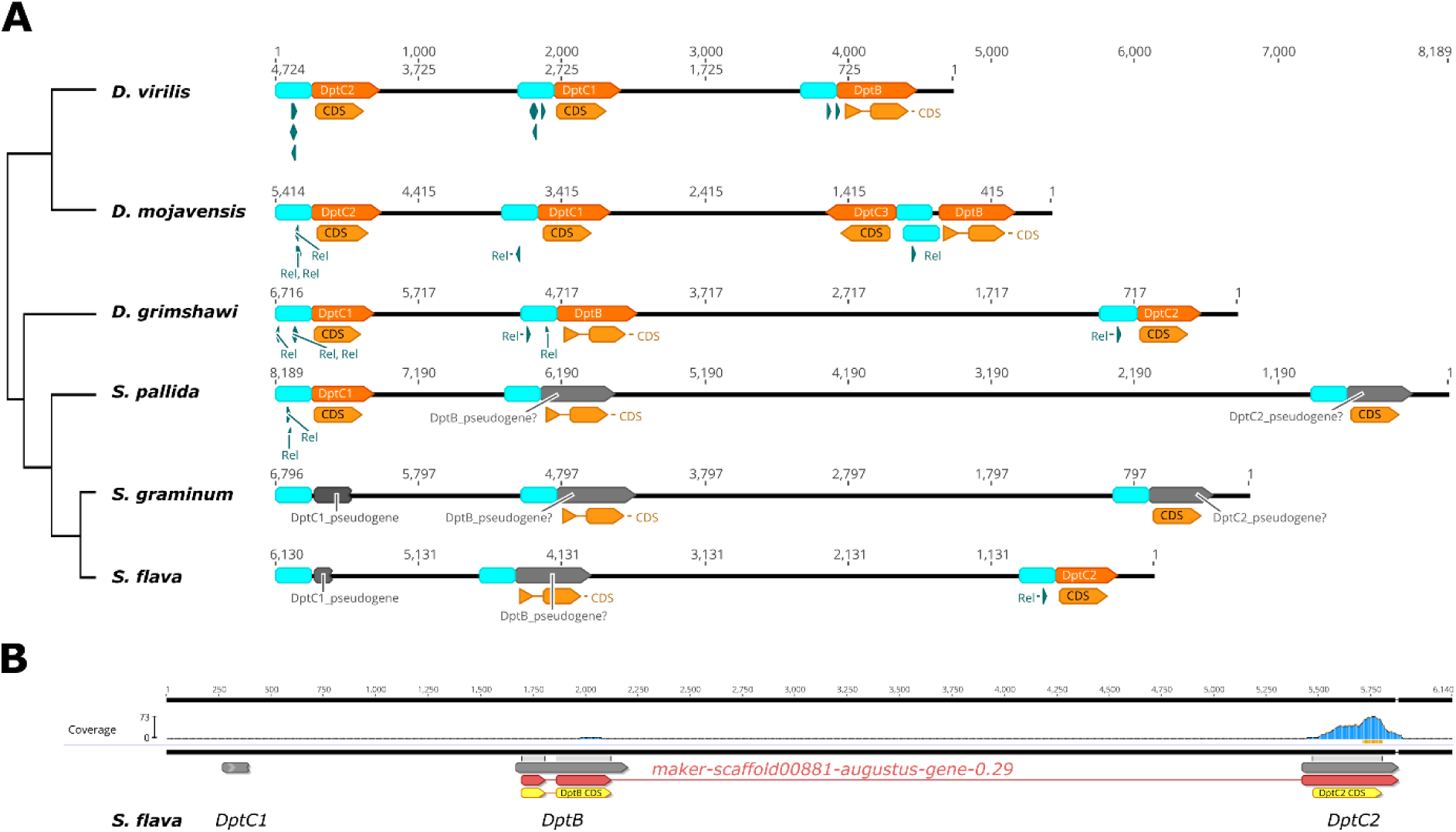
Analysis of *Diptericin* loci suggests pseudogenization of *Diptericin* genes in *Scaptomyza* compared to related flies. A) Light blue annotations indicate 250bp of sequence upstream of the transcription start site of each *Diptericin* gene, where *Relish* binding motifs are expected. These subgenus Drosophila species have had additional *DptA* locus duplication events (*DptA* named *“DptC”* here, according to cladistics of Hanson et al. (*48*), and see Fig. 4supp2): related to the main text, “*DptA1” = “DptC1”*. In both leaf-mining *Scaptomyza* species, *DptC1* has been pseudogenized by independent premature stop codons: Q43✱ in *S. flava* and G85✱ in *S. graminum* (numbering from start of CDS). *Scaptomyza pallida* feeds on decaying vegetation and mushroom (*54*), likely exposing it to *Providencia*, which should maintain selective pressure to keep *DptA/DptC*. Accordingly, its *DptC1* gene retains an intact coding sequence and Relish binding sites (“Rel” annotations). However, in all three *Scaptomyza* species, *DptB* appears to have lost Relish binding sites, which are needed for immune inducibility; Relish binding motif GGRDNNHHBS generated by Copley et al. (*55*). This same motif is abundant in the promoter of the *AttC* AMP gene across *Scaptomyza* species (Fig. 4supp6), indicating Scaptomyza still uses the same general motif as other species for Rel-mediated immune regulation. Two of the three *Scaptomyza* species have also lost all *Relish* binding sites in the promoter of *DptC2*, which could affect immune induction. B) Read mapping in the *Diptericin* gene region from RNA-seq of the *S. flava* midgut (SRR7649195). The *DptB* and *DptC2* genes are annotated as being part of the same mRNA transcript in data from (*56*), and so read data from this study attributes expression from either locus to the gene annotation “*maker-scaffold00881-augustus-gene-0.29*.” However, using BowTie2 in Geneious R10 with default settings, the vast majority of reads from this study mapping to *maker-sacffold00881-augustus-gene-0.29* stem from the *DptC2* gene region, which retains one Relish binding site (A), and is expected to be expressed. Only four reads were found that map to *Sfla\DptB*, while the same approach recovered 157 reads mapping back to *Sfla\DptC2* (a difference in expression of ∼39x). We confirmed our read mapping agreed with Gloss et al. (*56*) (full read mapping data from Gloss et al. (*56*) is unpublished). These data support the idea that *S. flava* has largely pseudogenized its *DptB* gene, confirm a lack of expression of *DptC1* that is pseudogenized by presence of a premature stop, and suggest that *DptC2* in *S. flava* is not pseudogenized in its coding sequence, promoter region, or expression.

**Figure 4supp6:**
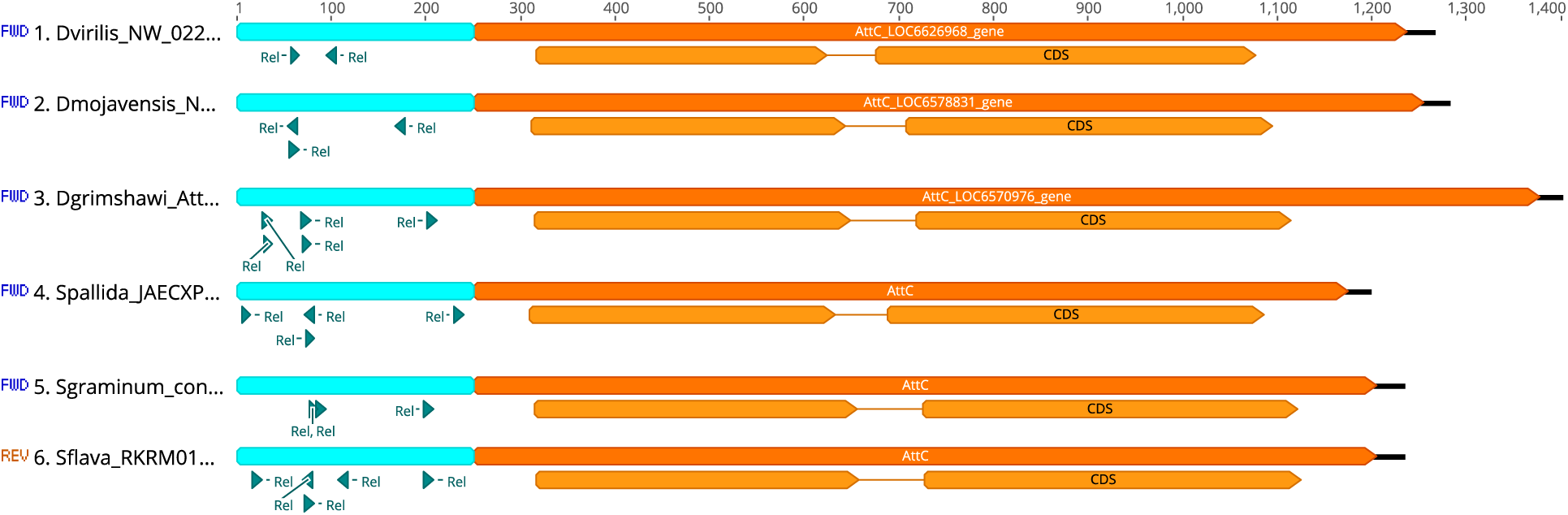
The Relish binding motif “GGRDNNHHBS” is observed upstream of the *AttC* AMP genes of *Scaptomyza* species. Scanning the *AttC* gene promoter confirms the same Relish binding motif used by other species for AMP induction is abundant in other Rel-regulated AMP genes of *Scaptomyza*. Thus, the absence of Relish binding sites upstream of Scaptomyza *Diptericins* described in Fig. 4supp5 likely affects their expression, consistent with pseudogenization.

**Figure 5supp1:**
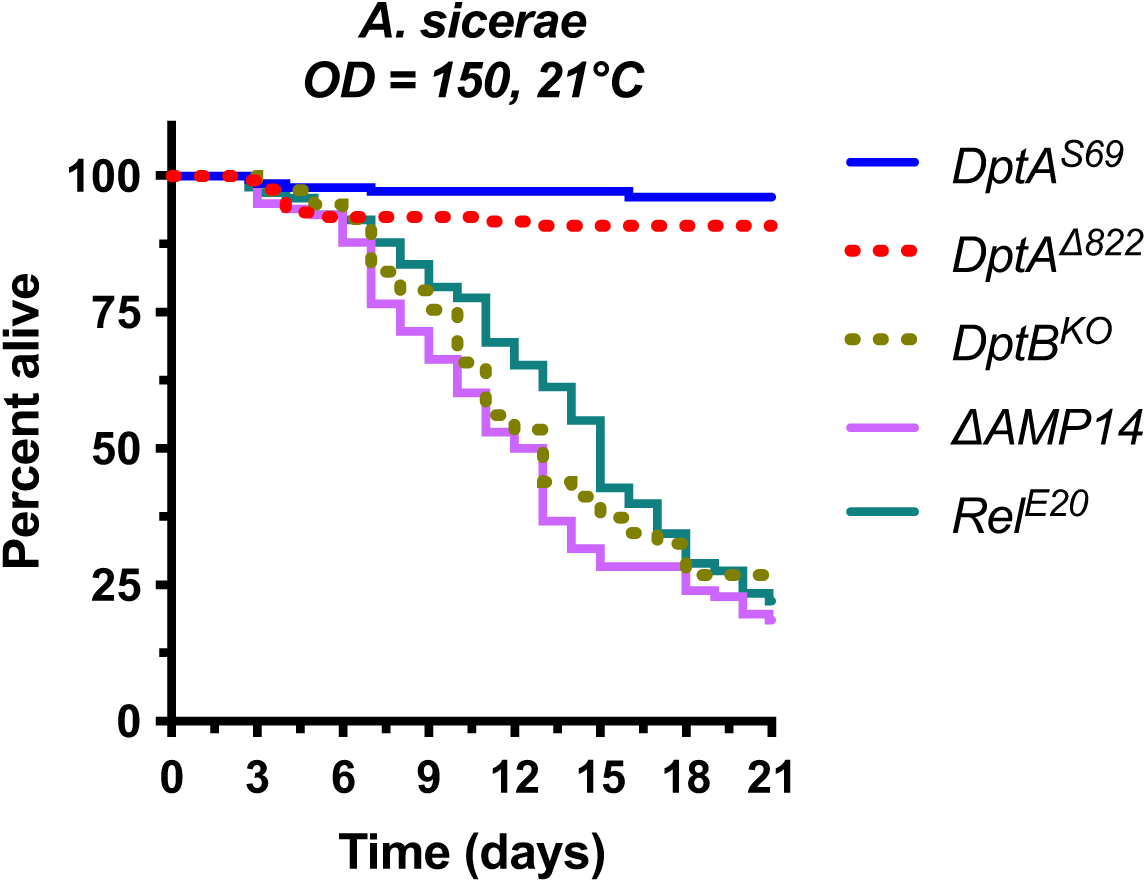
Use of 21°C reduces *A. sicerae* virulence compared to 25°C. Even at this temperature with reduced pathogen virulence, *DptB^KO^* susceptibility mirrors that of *ΔAMP14* and *Rel^E20^* flies. This further emphasizes the high importance of *DptB* as the main effector explaining Imd-mediated defence against this microbe.

**Figure 5supp2:**
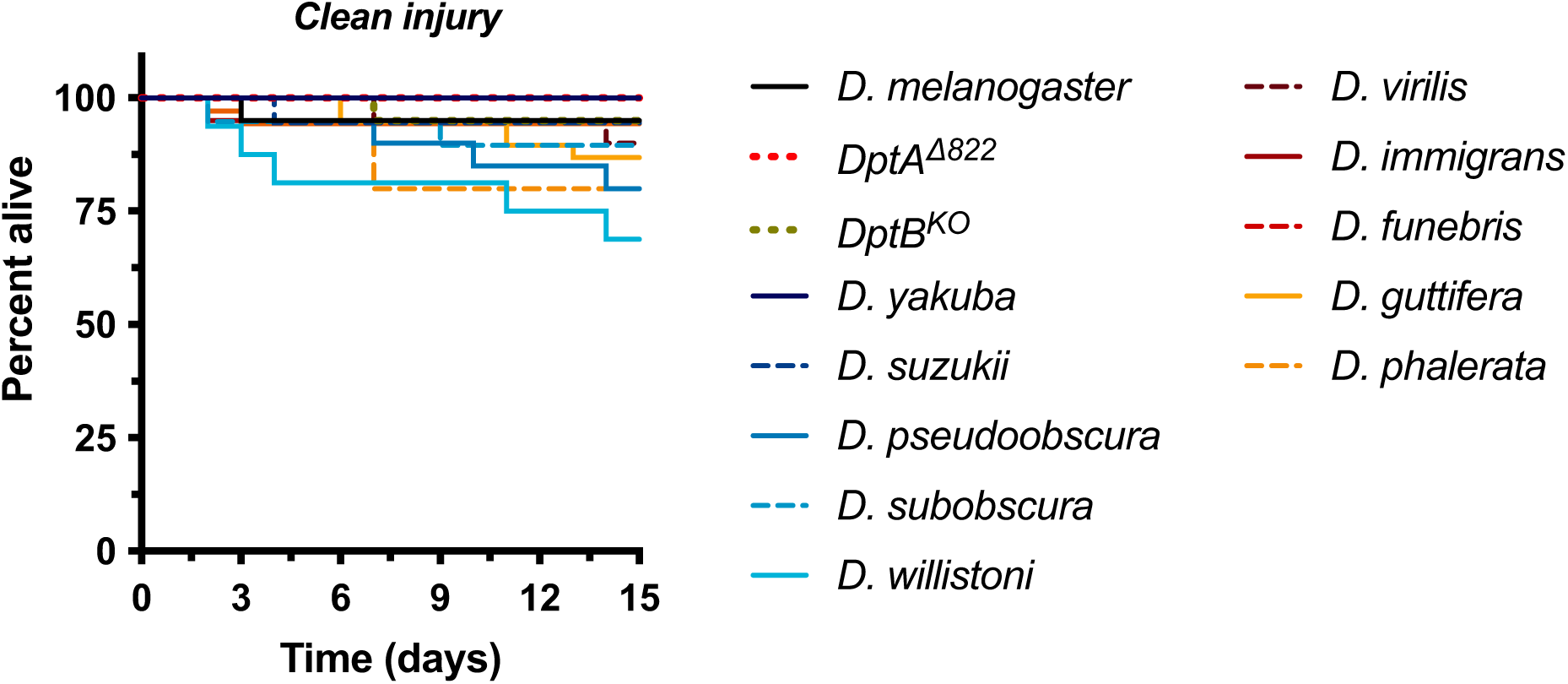
Clean injury controls of different fly species. No fly species succumbs to injury alone in a marked way, confirming that susceptibility to infections in Fig. 5 reflect bacteria-specific immune competence. Differential survival upon infection by *P. rettgeri* or *A. sicerae* per species in Fig. 5 also suggests no specific susceptibility to injury alone in any species tested.

**Figure 5supp3:**
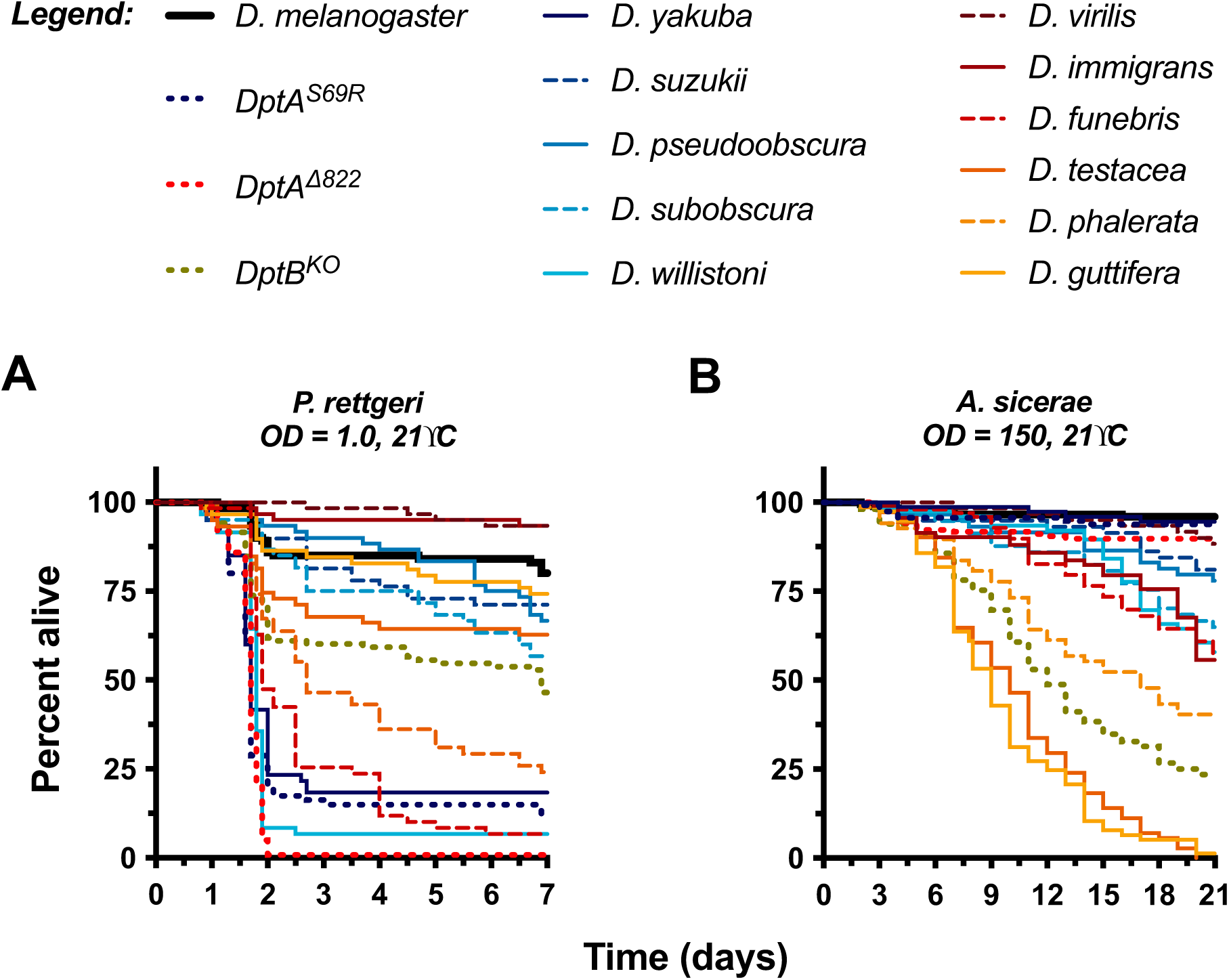
Survival curves for experiments summarized in Figure 5. *Acetobacter sicerae* experiments were monitored until day 21 to allow for complete mortality of the most susceptible fly strains/species given reduced virulence at this temperature (Fig. 5supp1).

