## Supplementary Data Files for "Ecology-relevant bacteria drive the evolution of host antimicrobial peptides in Drosophila": Aaceti_24-02-2022.pdf (from another version of this preprint: version 2, 2022-12-29)

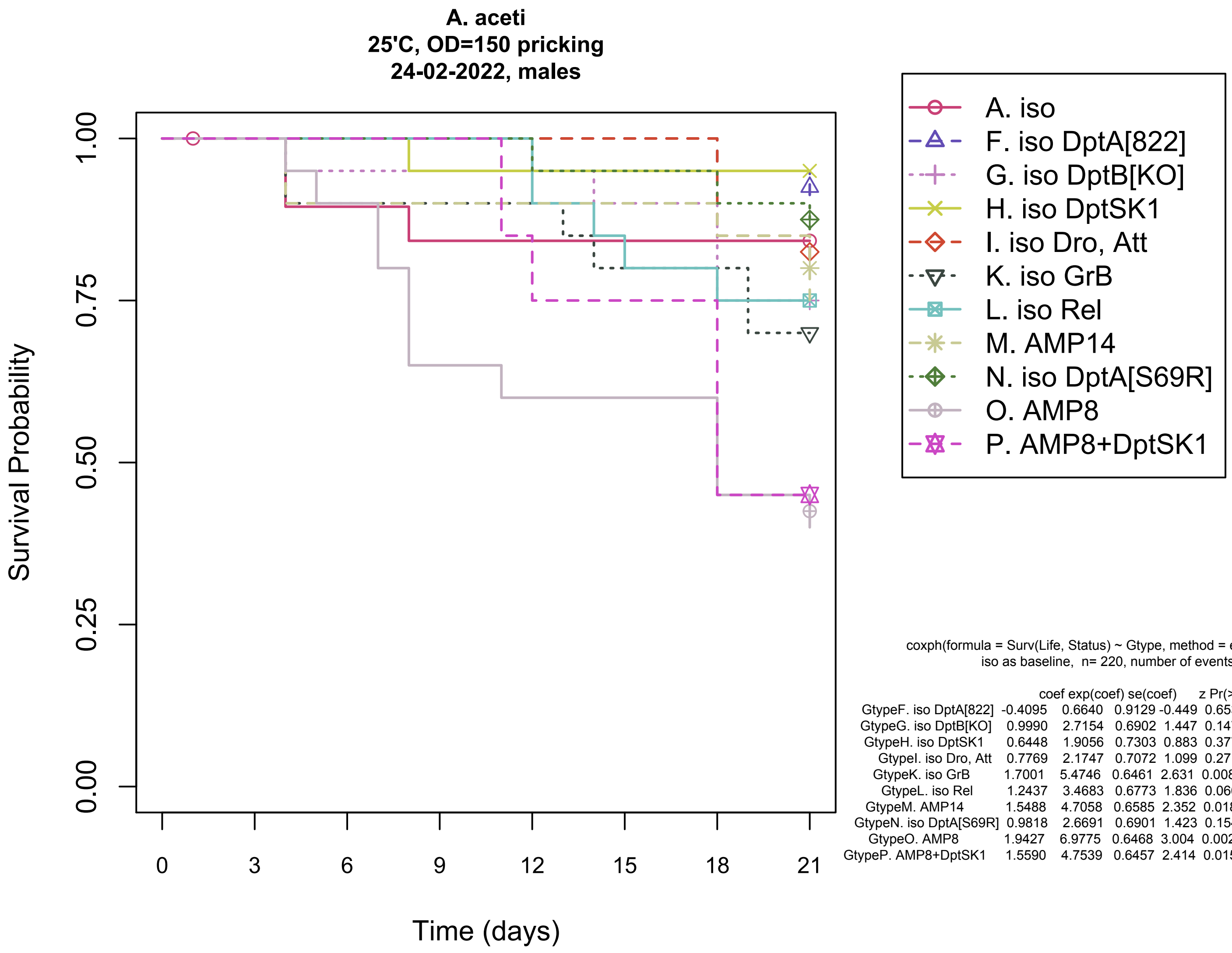


coxph(formula = Surv(Life, Status) ~ Gtype, method = efron)
iso as baseline, n= 220, number of events= 84

| | coef | exp(coef) | se(coef) | z | Pr(>\|z\|) | |
|---|---|---|---|---|---|---|
| GtypeF. iso DptA[822] | -0.4095 | 0.6640 | 0.9129 | -0.449 | 0.65375 | |
| GtypeG. iso DptB[KO] | 0.9990 | 2.7154 | 0.6902 | 1.447 | 0.14778 | |
| GtypeH. iso DptSK1 | 0.6448 | 1.9056 | 0.7303 | 0.883 | 0.37728 | |
| GtypeI. iso Dro, Att | 0.7769 | 2.1747 | 0.7072 | 1.099 | 0.27194 | |
| GtypeK. iso GrB | 1.7001 | 5.4746 | 0.6461 | 2.631 | 0.00851 | ** |
| GtypeL. iso Rel | 1.2437 | 3.4683 | 0.6773 | 1.836 | 0.06632 | . |
| GtypeM. AMP14 | 1.5488 | 4.7058 | 0.6585 | 2.352 | 0.01867 | * |
| GtypeN. iso DptA[S69R] | 0.9818 | 2.6691 | 0.6901 | 1.423 | 0.15486 | |
| GtypeO. AMP8 | 1.9427 | 6.9775 | 0.6468 | 3.004 | 0.00267 | ** |
| GtypeP. AMP8+DptSK1 | 1.5590 | 4.7539 | 0.6457 | 2.414 | 0.01577 | * |
