## Supplementary Data Files for "Ecology-relevant bacteria drive the evolution of host antimicrobial peptides in Drosophila": Aaceti_CoxPH.pdf (from another version of this preprint: version 2, 2022-12-29)

**A. aceti**
**25'C, OD=150 pricking, deaths within first 2 days censored, exp cens at 14dpi**
**15-12-2021, 24-02-2022, 05-04-2022 males**

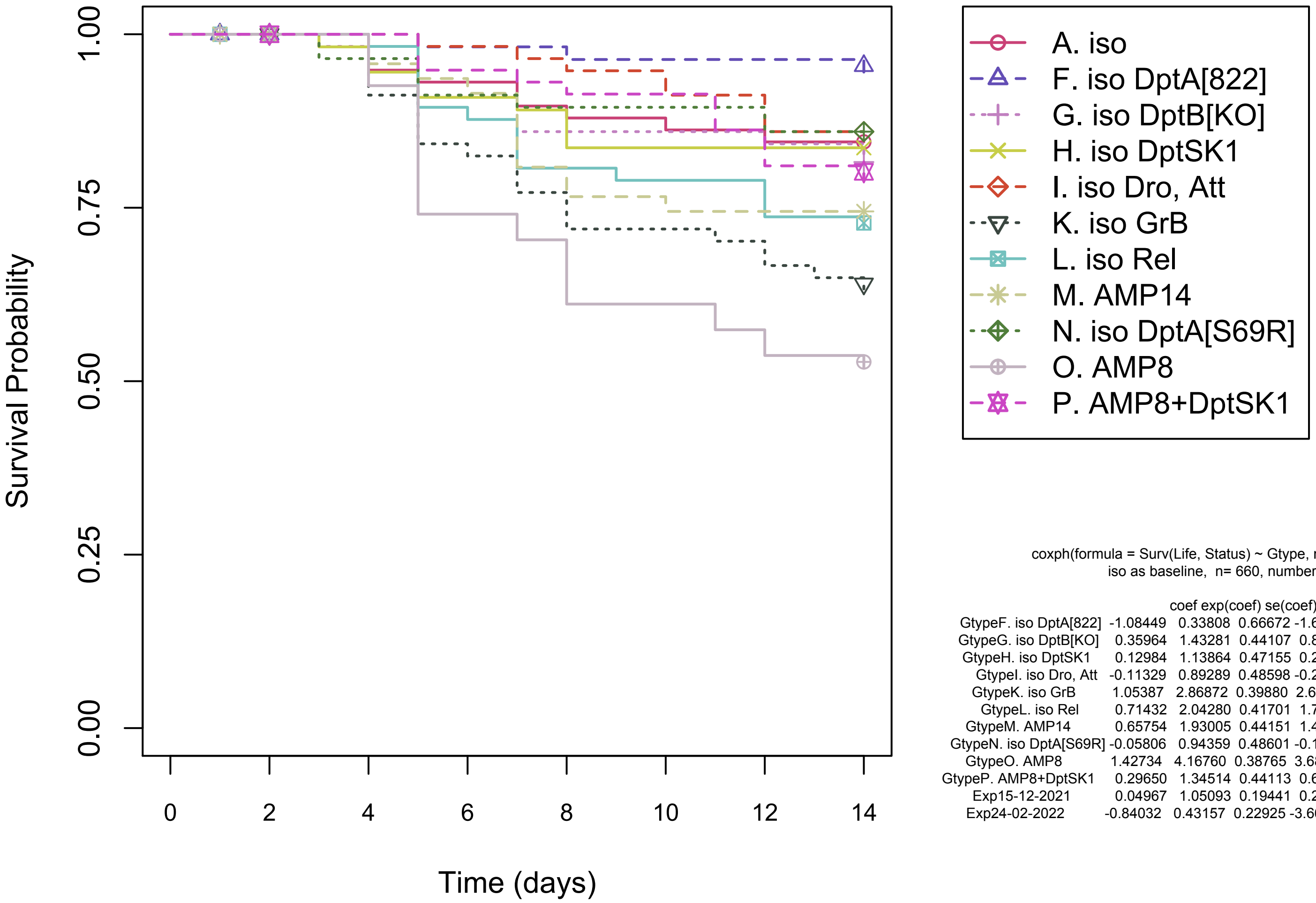


coxph(formula = Surv(Life, Status) ~ Gtype, method = efron)
iso as baseline, n= 660, number of events= 136

| | coef | exp(coef) | se(coef) | z | Pr(>\|z\|) | |
|---|---|---|---|---|---|---|
| GtypeF. iso DptA[822] | -1.08449 | 0.33808 | 0.66672 | -1.627 | 0.103825 | |
| GtypeG. iso DptB[KO] | 0.35964 | 1.43281 | 0.44107 | 0.815 | 0.414861 | |
| GtypeH. iso DptSK1 | 0.12984 | 1.13864 | 0.47155 | 0.275 | 0.783056 | |
| GtypeI. iso Dro, Att | -0.11329 | 0.89289 | 0.48598 | -0.233 | 0.815664 | |
| GtypeK. iso GrB | 1.05387 | 2.86872 | 0.39880 | 2.643 | 0.008227 | ** |
| GtypeL. iso Rel | 0.71432 | 2.04280 | 0.41701 | 1.713 | 0.086722 | . |
| GtypeM. AMP14 | 0.65754 | 1.93005 | 0.44151 | 1.489 | 0.136405 | |
| GtypeN. iso DptA[S69R] | -0.05806 | 0.94359 | 0.48601 | -0.119 | 0.904911 | |
| GtypeO. AMP8 | 1.42734 | 4.16760 | 0.38765 | 3.682 | 0.000231 | *** |
| GtypeP. AMP8+DptSK1 | 0.29650 | 1.34514 | 0.44113 | 0.672 | 0.501500 | |
| Exp15-12-2021 | 0.04967 | 1.05093 | 0.19441 | 0.256 | 0.798336 | |
| Exp24-02-2022 | -0.84032 | 0.43157 | 0.22925 | -3.666 | 0.000247 | *** |
