## Supplementary Data Files for "Ecology-relevant bacteria drive the evolution of host antimicrobial peptides in Drosophila": Aindon_03-12-2021.pdf (from another version of this preprint: version 2, 2022-12-29)

### A. indonesiensis
### 25'C, OD=150 pricking
### 03-12-2021, males

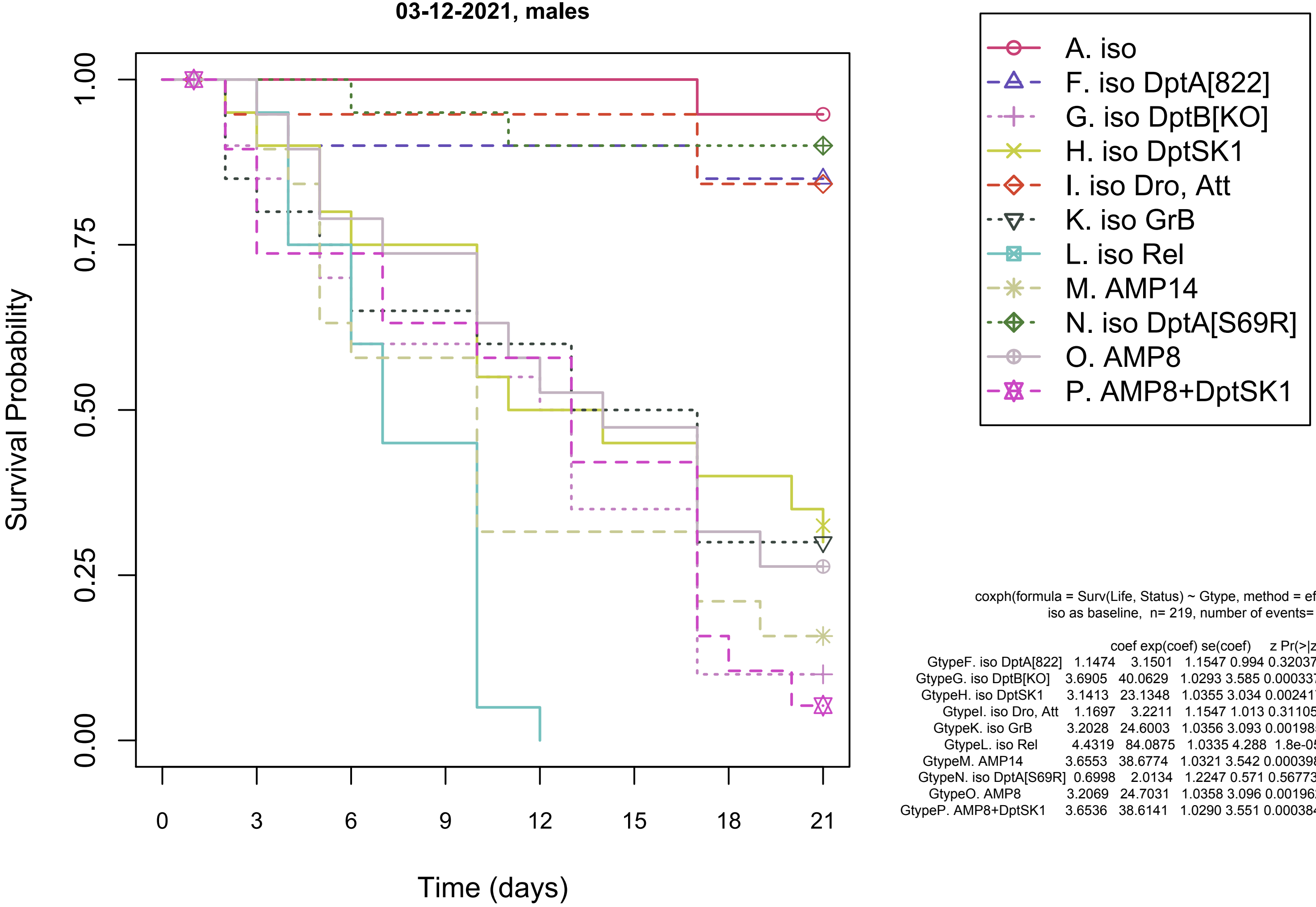


coxph(formula = Surv(Life, Status) ~ Gtype, method = efron)
iso as baseline, n= 219, number of events= 123

| | coef | exp(coef) | se(coef) | z | Pr(>\|z\|) | |
|---|---|---|---|---|---|---|
| GtypeF. iso DptA[822] | 1.1474 | 3.1501 | 1.1547 | 0.994 | 0.320371 | |
| GtypeG. iso DptB[KO] | 3.6905 | 40.0629 | 1.0293 | 3.585 | 0.000337 | *** |
| GtypeH. iso DptSK1 | 3.1413 | 23.1348 | 1.0355 | 3.034 | 0.002417 | ** |
| GtypeI. iso Dro, Att | 1.1697 | 3.2211 | 1.1547 | 1.013 | 0.311053 | |
| GtypeK. iso GrB | 3.2028 | 24.6003 | 1.0356 | 3.093 | 0.001985 | ** |
| GtypeL. iso Rel | 4.4319 | 84.0875 | 1.0335 | 4.288 | 1.8e-05 | *** |
| GtypeM. AMP14 | 3.6553 | 38.6774 | 1.0321 | 3.542 | 0.000398 | *** |
| GtypeN. iso DptA[S69R] | 0.6998 | 2.0134 | 1.2247 | 0.571 | 0.567737 | |
| GtypeO. AMP8 | 3.2069 | 24.7031 | 1.0358 | 3.096 | 0.001962 | ** |
| GtypeP. AMP8+DptSK1 | 3.6536 | 38.6141 | 1.0290 | 3.551 | 0.000384 | *** |
