## Supplementary Data Files for "Ecology-relevant bacteria drive the evolution of host antimicrobial peptides in Drosophila": Aindon_20-01-2022.pdf (from another version of this preprint: version 2, 2022-12-29)

**A. indonesiensis**
**25'C, OD=150 pricking**
**20-01-2022, males**

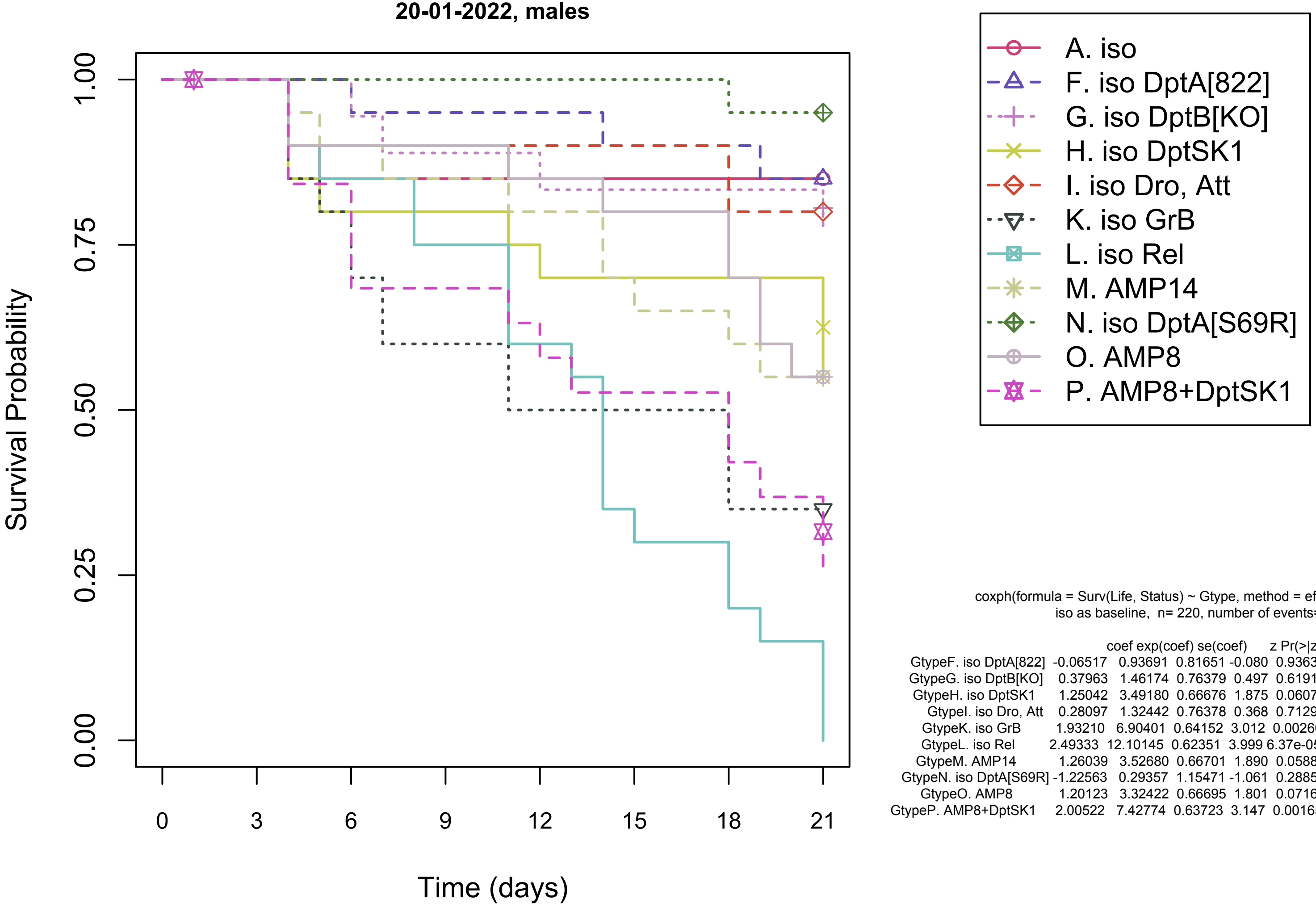


coxph(formula = Surv(Life, Status) ~ Gtype, method = efron)
iso as baseline,  n= 220, number of events= 89

| | coef | exp(coef) | se(coef) | z | Pr(>\|z\|) | |
|---|---|---|---|---|---|---|
| GtypeF. iso DptA[822] | -0.06517 | 0.93691 | 0.81651 | -0.080 | 0.93639 | |
| GtypeG. iso DptB[KO] | 0.37963 | 1.46174 | 0.76379 | 0.497 | 0.61917 | |
| GtypeH. iso DptSK1 | 1.25042 | 3.49180 | 0.66676 | 1.875 | 0.06074 | . |
| GtypeI. iso Dro, Att | 0.28097 | 1.32442 | 0.76378 | 0.368 | 0.71297 | |
| GtypeK. iso GrB | 1.93210 | 6.90401 | 0.64152 | 3.012 | 0.00260 | ** |
| GtypeL. iso Rel | 2.49333 | 12.10145 | 0.62351 | 3.999 | 6.37e-05 | *** |
| GtypeM. AMP14 | 1.26039 | 3.52680 | 0.66701 | 1.890 | 0.05881 | . |
| GtypeN. iso DptA[S69R] | -1.22563 | 0.29357 | 1.15471 | -1.061 | 0.28850 | |
| GtypeO. AMP8 | 1.20123 | 3.32422 | 0.66695 | 1.801 | 0.07169 | . |
| GtypeP. AMP8+DptSK1 | 2.00522 | 7.42774 | 0.63723 | 3.147 | 0.00165 | ** |
