## Supplementary Data Files for "Ecology-relevant bacteria drive the evolution of host antimicrobial peptides in Drosophila": Aindo_CoxPH.pdf (from another version of this preprint: version 2, 2022-12-29)

**Acetobacter indonesiensis DSM15552**
**25'C, OD=150 pricking**
**CoxPH 03-12-2021, 20-01-2022, 03-05-2022 males**

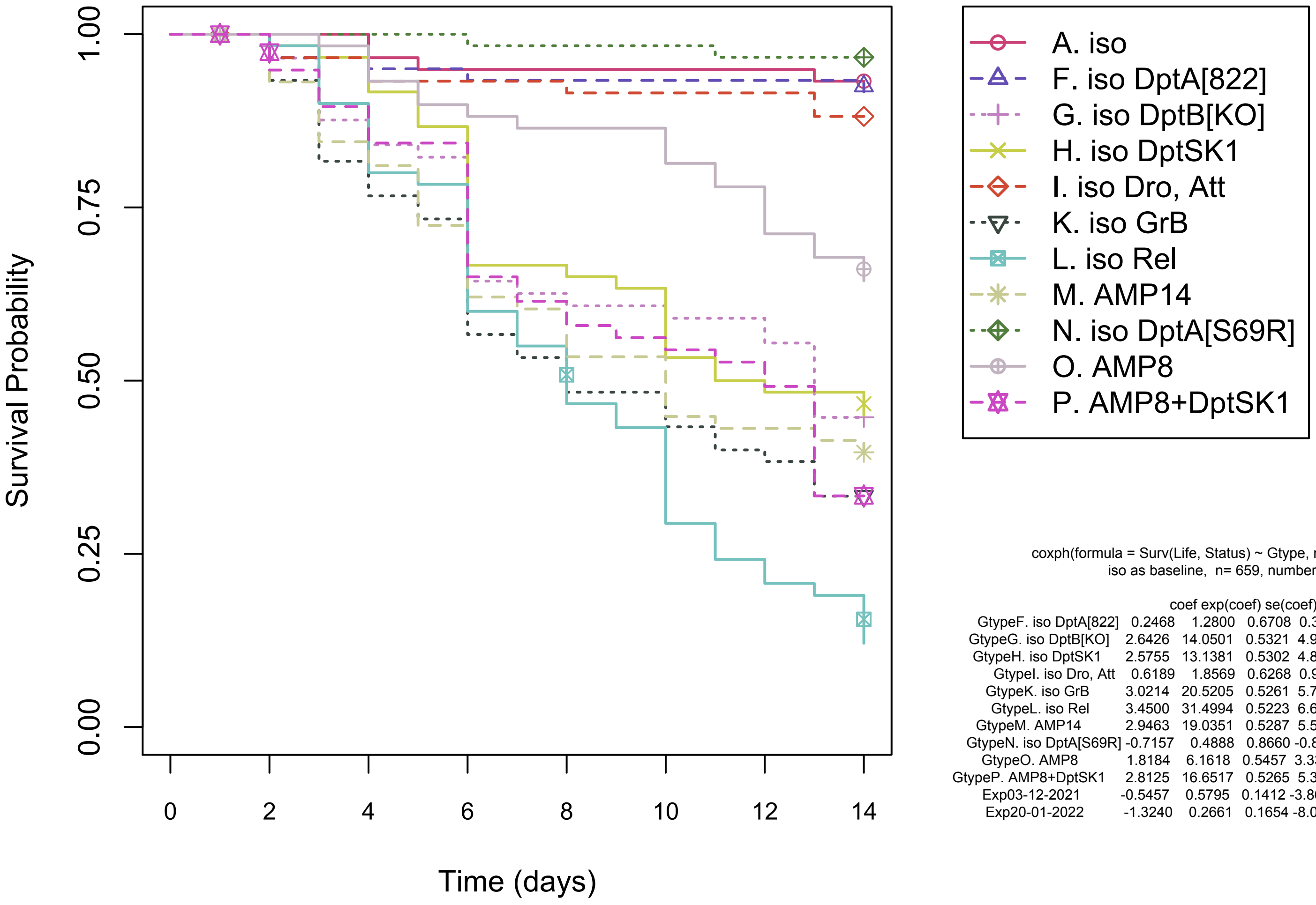


coxph(formula = Surv(Life, Status) ~ Gtype, method = efron)
iso as baseline,  n= 659, number of events= 269

| | coef | exp(coef) | se(coef) | z | Pr(>\|z\|) | |
|---|---|---|---|---|---|---|
| GtypeF. iso DptA[822] | 0.2468 | 1.2800 | 0.6708 | 0.368 | 0.712921 | |
| GtypeG. iso DptB[KO] | 2.6426 | 14.0501 | 0.5321 | 4.966 | 6.84e-07 | *** |
| GtypeH. iso DptSK1 | 2.5755 | 13.1381 | 0.5302 | 4.857 | 1.19e-06 | *** |
| GtypeI. iso Dro, Att | 0.6189 | 1.8569 | 0.6268 | 0.987 | 0.323437 | |
| GtypeK. iso GrB | 3.0214 | 20.5205 | 0.5261 | 5.743 | 9.28e-09 | *** |
| GtypeL. iso Rel | 3.4500 | 31.4994 | 0.5223 | 6.605 | 3.98e-11 | *** |
| GtypeM. AMP14 | 2.9463 | 19.0351 | 0.5287 | 5.573 | 2.50e-08 | *** |
| GtypeN. iso DptA[S69R] | -0.7157 | 0.4888 | 0.8660 | -0.826 | 0.408536 | |
| GtypeO. AMP8 | 1.8184 | 6.1618 | 0.5457 | 3.332 | 0.000861 | *** |
| GtypeP. AMP8+DptSK1 | 2.8125 | 16.6517 | 0.5265 | 5.342 | 9.20e-08 | *** |
| Exp03-12-2021 | -0.5457 | 0.5795 | 0.1412 | -3.865 | 0.000111 | *** |
| Exp20-01-2022 | -1.3240 | 0.2661 | 0.1654 | -8.003 | 1.22e-15 | *** |
