## Supplementary Data Files for "Ecology-relevant bacteria drive the evolution of host antimicrobial peptides in Drosophila": Aorient_01-03-2022.pdf (from another version of this preprint: version 2, 2022-12-29)

**A. orientalis**
**25'C, OD=150 pricking**
**01-03-2022, males**

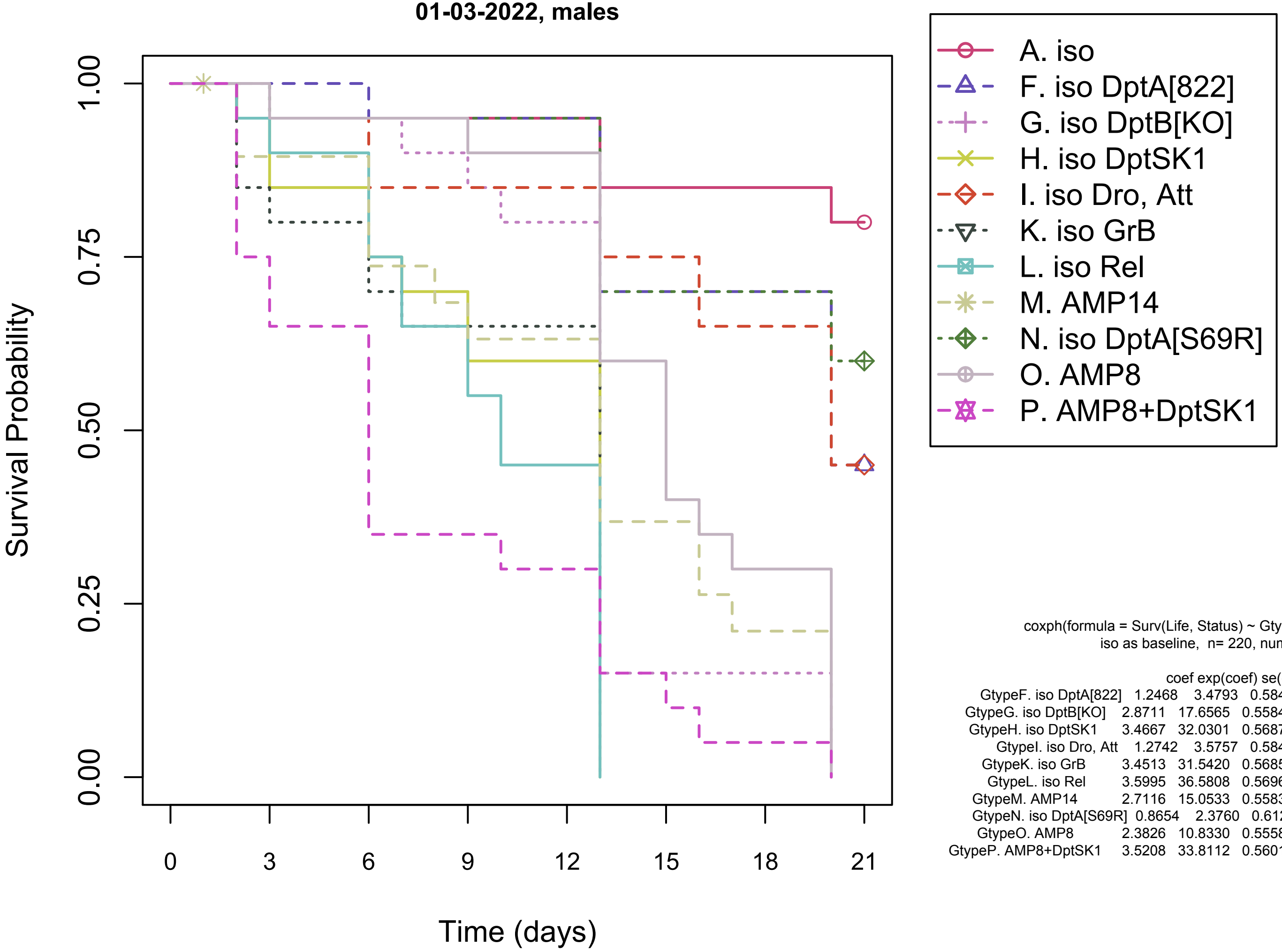


coxph(formula = Surv(Life, Status) ~ Gtype, method = efron)
iso as baseline, n= 220, number of events= 173

| | coef | exp(coef) | se(coef) | z | Pr(>\|z\|) | |
|---|---|---|---|---|---|---|
| GtypeF. iso DptA[822] | 1.2468 | 3.4793 | 0.5843 | 2.134 | 0.0329 | * |
| GtypeG. iso DptB[KO] | 2.8711 | 17.6565 | 0.5584 | 5.142 | 2.72e-07 | *** |
| GtypeH. iso DptSK1 | 3.4667 | 32.0301 | 0.5687 | 6.095 | 1.09e-09 | *** |
| GtypeI. iso Dro, Att | 1.2742 | 3.5757 | 0.5843 | 2.181 | 0.0292 | * |
| GtypeK. iso GrB | 3.4513 | 31.5420 | 0.5685 | 6.071 | 1.27e-09 | *** |
| GtypeL. iso Rel | 3.5995 | 36.5808 | 0.5696 | 6.320 | 2.62e-10 | *** |
| GtypeM. AMP14 | 2.7116 | 15.0533 | 0.5583 | 4.857 | 1.19e-06 | *** |
| GtypeN. iso DptA[S69R] | 0.8654 | 2.3760 | 0.6125 | 1.413 | 0.1577 | |
| GtypeO. AMP8 | 2.3826 | 10.8330 | 0.5558 | 4.287 | 1.81e-05 | *** |
| GtypeP. AMP8+DptSK1 | 3.5208 | 33.8112 | 0.5601 | 6.286 | 3.25e-10 | *** |
