## Supplementary Data Files for "Ecology-relevant bacteria drive the evolution of host antimicrobial peptides in Drosophila": Aorient_07-12-2021.pdf (from another version of this preprint: version 2, 2022-12-29)

**A. orientalis**
**25'C, OD=150 pricking**
**07-12-2021, males, CAUTION: medium contam? Flies contam?**

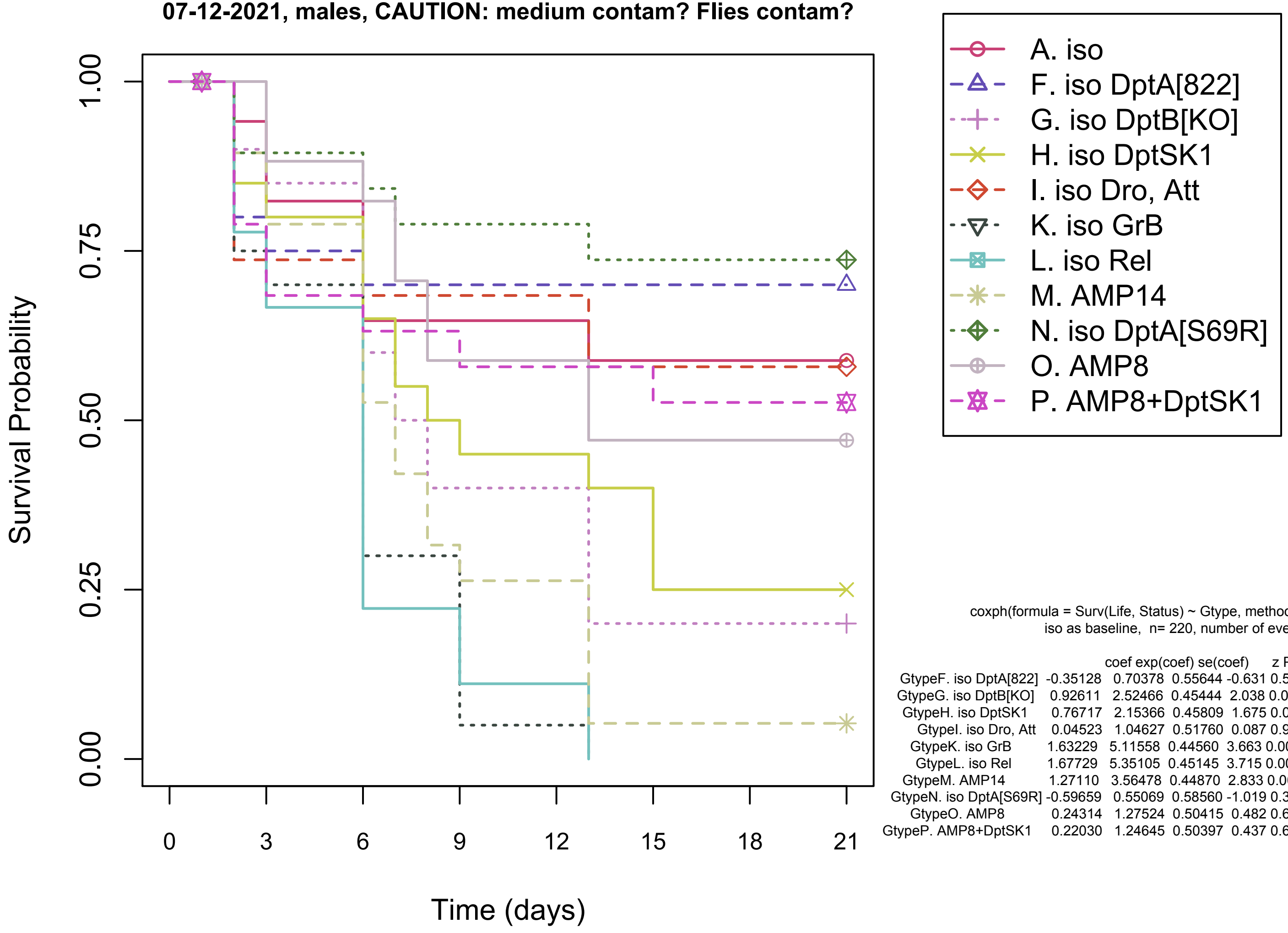


coxph(formula = Surv(Life, Status) ~ Gtype, method = efron)
iso as baseline,  n= 220, number of events= 131

| | coef | exp(coef) | se(coef) | z | Pr(>\|z\|) | |
|---|---|---|---|---|---|---|
| GtypeF. iso DptA[822] | -0.35128 | 0.70378 | 0.55644 | -0.631 | 0.527837 | |
| GtypeG. iso DptB[KO] | 0.92611 | 2.52466 | 0.45444 | 2.038 | 0.041558 | * |
| GtypeH. iso DptSK1 | 0.76717 | 2.15366 | 0.45809 | 1.675 | 0.093992 | . |
| GtypeI. iso Dro, Att | 0.04523 | 1.04627 | 0.51760 | 0.087 | 0.930360 | |
| GtypeK. iso GrB | 1.63229 | 5.11558 | 0.44560 | 3.663 | 0.000249 | *** |
| GtypeL. iso Rel | 1.67729 | 5.35105 | 0.45145 | 3.715 | 0.000203 | *** |
| GtypeM. AMP14 | 1.27110 | 3.56478 | 0.44870 | 2.833 | 0.004614 | ** |
| GtypeN. iso DptA[S69R] | -0.59659 | 0.55069 | 0.58560 | -1.019 | 0.308316 | |
| GtypeO. AMP8 | 0.24314 | 1.27524 | 0.50415 | 0.482 | 0.629616 | |
| GtypeP. AMP8+DptSK1 | 0.22030 | 1.24645 | 0.50397 | 0.437 | 0.662015 | |
