## Supplementary Data Files for "Ecology-relevant bacteria drive the evolution of host antimicrobial peptides in Drosophila": Aorient_17-12-2021.pdf (from another version of this preprint: version 2, 2022-12-29)

**A. orientalis**
**25'C, OD=150 pricking**
**17-12-2021, males**

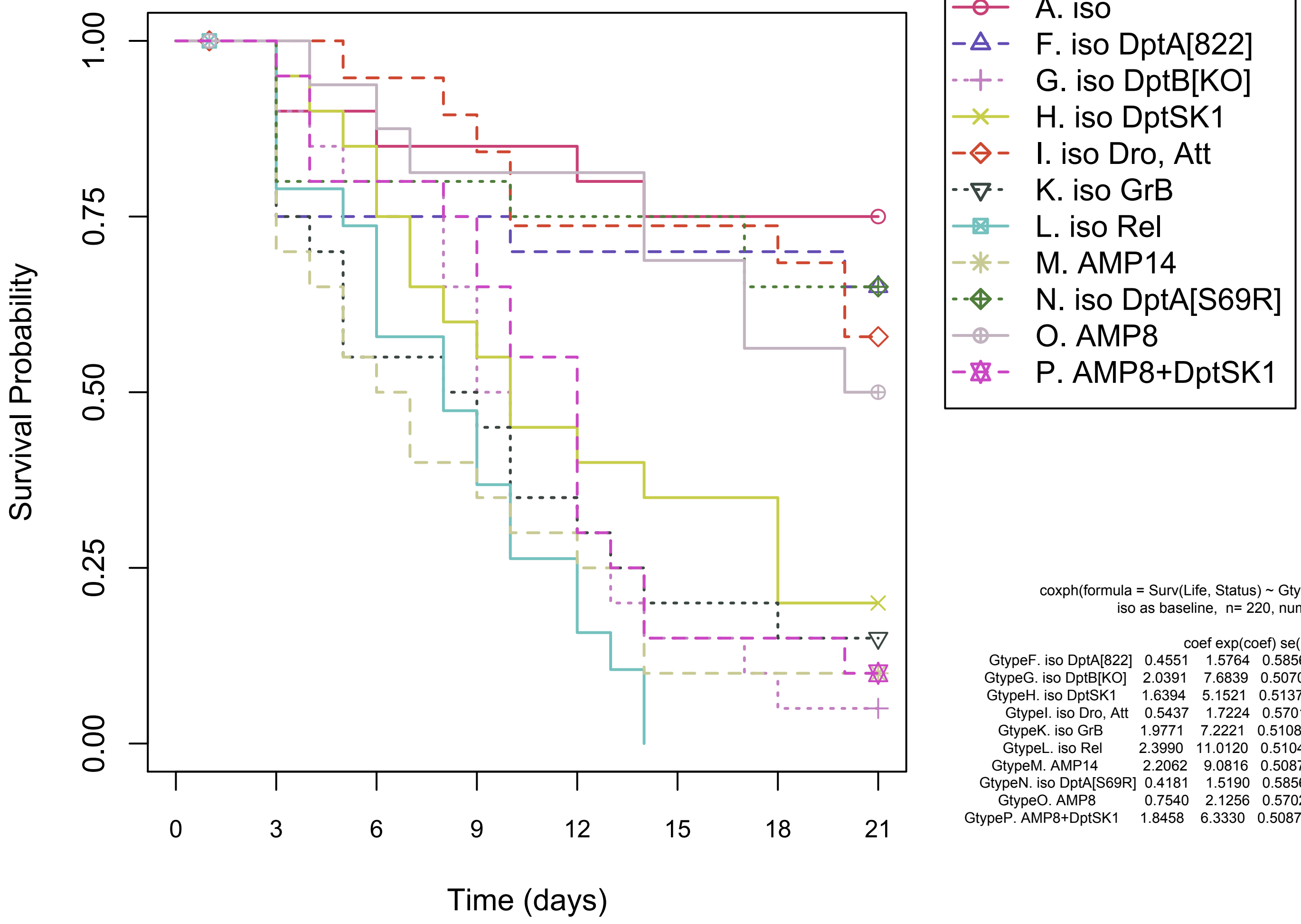


coxph(formula = Surv(Life, Status) ~ Gtype, method = efron)
iso as baseline,  n= 220, number of events= 142

| | coef | exp(coef) | se(coef) | z | Pr(>\|z\|) | |
|---|---|---|---|---|---|---|
| GtypeF. iso DptA[822] | 0.4551 | 1.5764 | 0.5856 | 0.777 | 0.437016 | |
| GtypeG. iso DptB[KO] | 2.0391 | 7.6839 | 0.5070 | 4.022 | 5.77e-05 | *** |
| GtypeH. iso DptSK1 | 1.6394 | 5.1521 | 0.5137 | 3.192 | 0.001415 | ** |
| GtypeI. iso Dro, Att | 0.5437 | 1.7224 | 0.5701 | 0.954 | 0.340231 | |
| GtypeK. iso GrB | 1.9771 | 7.2221 | 0.5108 | 3.871 | 0.000108 | *** |
| GtypeL. iso Rel | 2.3990 | 11.0120 | 0.5104 | 4.700 | 2.60e-06 | *** |
| GtypeM. AMP14 | 2.2062 | 9.0816 | 0.5087 | 4.337 | 1.45e-05 | *** |
| GtypeN. iso DptA[S69R] | 0.4181 | 1.5190 | 0.5856 | 0.714 | 0.475256 | |
| GtypeO. AMP8 | 0.7540 | 2.1256 | 0.5702 | 1.322 | 0.186063 | |
| GtypeP. AMP8+DptSK1 | 1.8458 | 6.3330 | 0.5087 | 3.628 | 0.000285 | *** |
