## Supplementary Data Files for "Ecology-relevant bacteria drive the evolution of host antimicrobial peptides in Drosophila": Aorient_CoxPH.pdf (from another version of this preprint: version 2, 2022-12-29)

**A. orientalis**
**25'C, OD=150 pricking: 17-12-2021, 01-03-2022, males**
**07-12-2021 medium possibly contam, and AMP8/AMP10 mixup possible. Censored.**

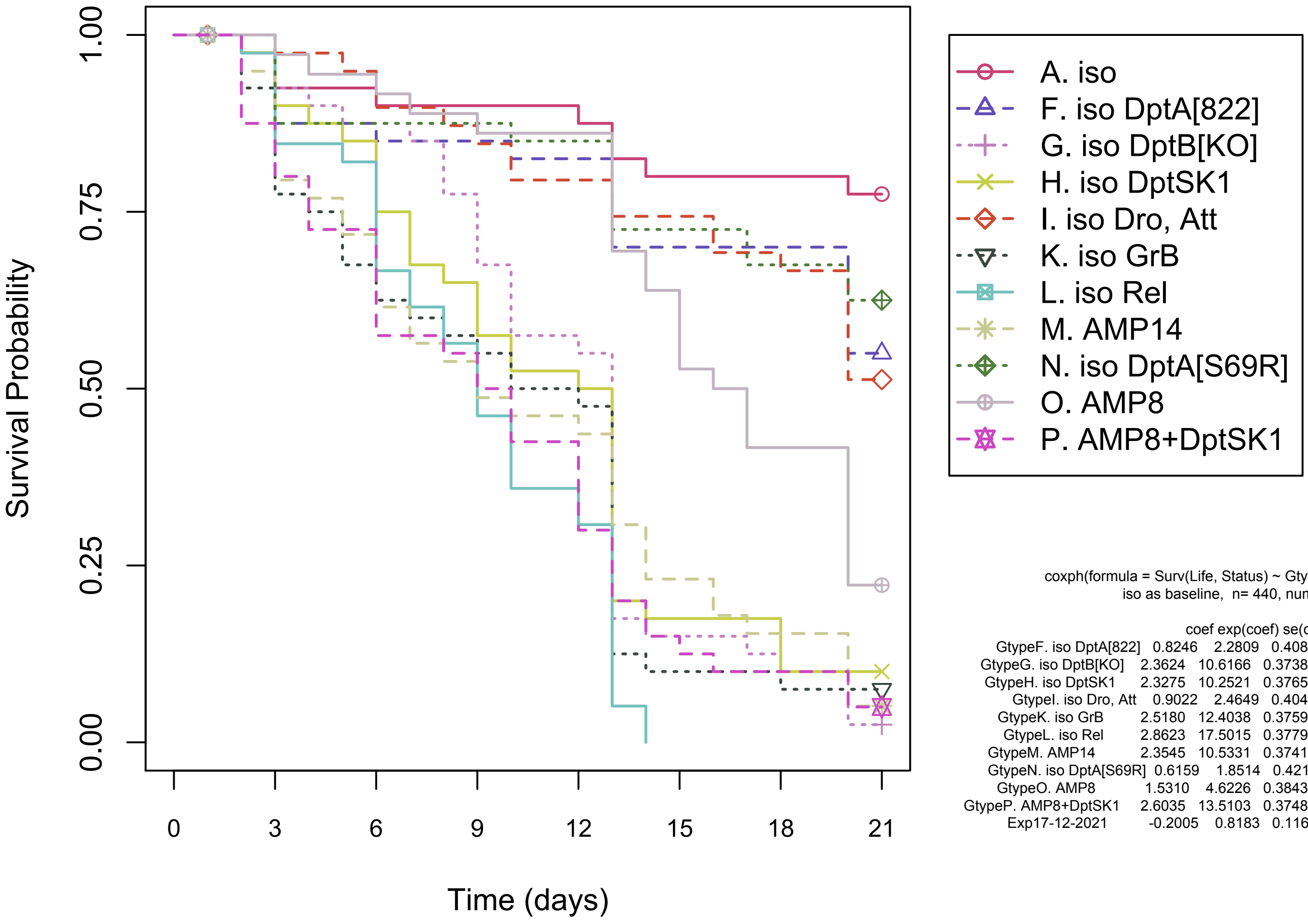


coxph(formula = Surv(Life, Status) ~ Gtype, method = efron)
iso as baseline, n= 440, number of events= 315

| | coef | exp(coef) | se(coef) | z | Pr(>\|z\|) | |
|---|---|---|---|---|---|---|
| GtypeF. iso DptA[822] | 0.8246 | 2.2809 | 0.4083 | 2.019 | 0.0434 | * |
| GtypeG. iso DptB[KO] | 2.3624 | 10.6166 | 0.3738 | 6.320 | 2.61e-10 | *** |
| GtypeH. iso DptSK1 | 2.3275 | 10.2521 | 0.3765 | 6.182 | 6.34e-10 | *** |
| GtypeI. iso Dro, Att | 0.9022 | 2.4649 | 0.4048 | 2.229 | 0.0258 | * |
| GtypeK. iso GrB | 2.5180 | 12.4038 | 0.3759 | 6.699 | 2.10e-11 | *** |
| GtypeL. iso Rel | 2.8623 | 17.5015 | 0.3779 | 7.574 | 3.63e-14 | *** |
| GtypeM. AMP14 | 2.3545 | 10.5331 | 0.3741 | 6.294 | 3.09e-10 | *** |
| GtypeN. iso DptA[S69R] | 0.6159 | 1.8514 | 0.4217 | 1.461 | 0.1441 | |
| GtypeO. AMP8 | 1.5310 | 4.6226 | 0.3843 | 3.984 | 6.78e-05 | *** |
| GtypeP. AMP8+DptSK1 | 2.6035 | 13.5103 | 0.3748 | 6.947 | 3.74e-12 | *** |
| Exp17-12-2021 | -0.2005 | 0.8183 | 0.1163 | -1.723 | 0.0849 | . |
