## Supplementary Data Files for "Ecology-relevant bacteria drive the evolution of host antimicrobial peptides in Drosophila": Apomorum_05-10-2021.pdf (from another version of this preprint: version 2, 2022-12-29)

**Apomorum**
**25'C, OD=150 pricking**
**05-10-2021, males**

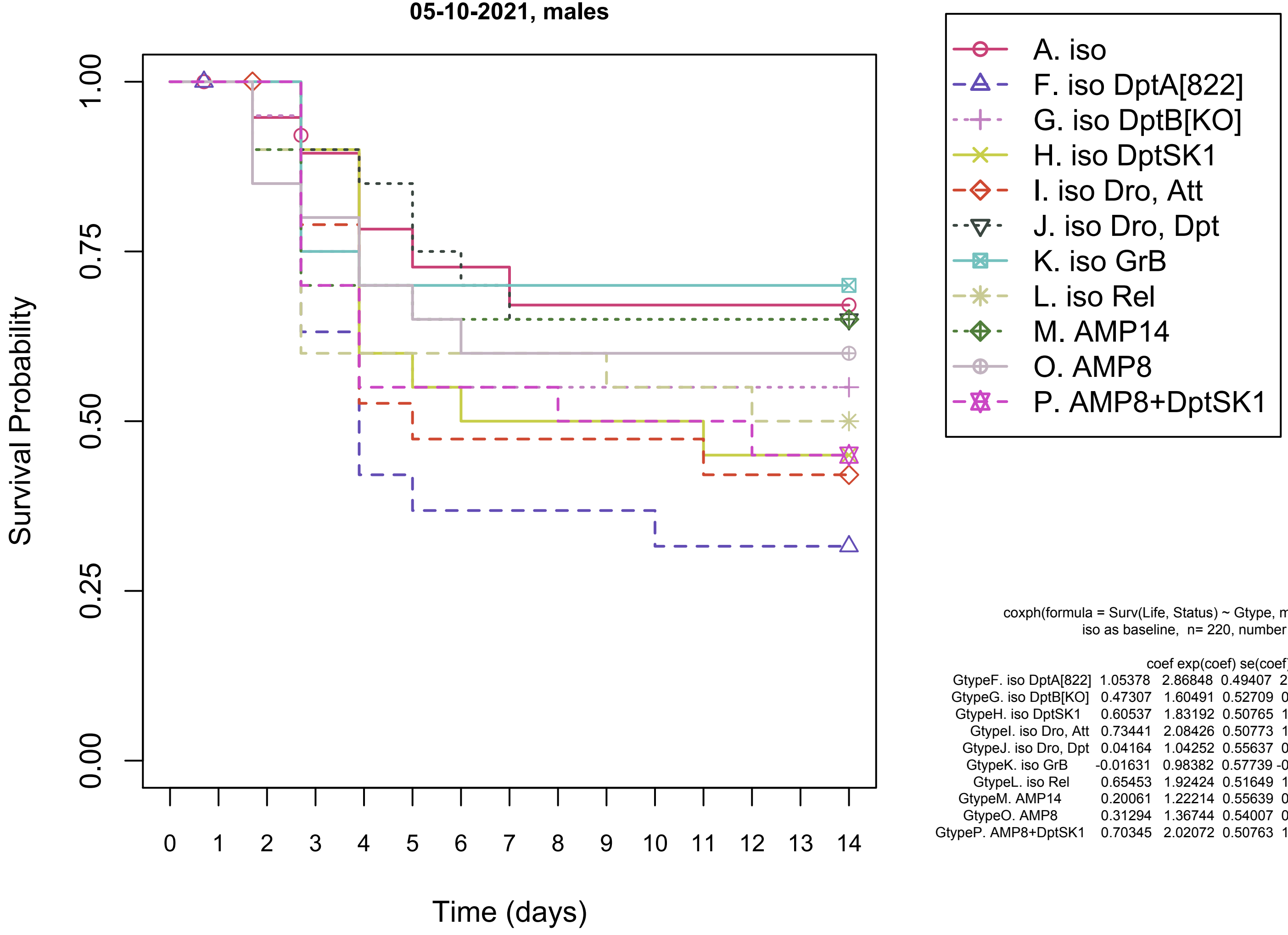


coxph(formula = Surv(Life, Status) ~ Gtype, method = efron)
iso as baseline, n= 220, number of events= 99

| | coef | exp(coef) | se(coef) | z | Pr(>\|z\|) | |
|---|---|---|---|---|---|---|
| GtypeF. iso DptA[822] | 1.05378 | 2.86848 | 0.49407 | 2.133 | 0.0329 | * |
| GtypeG. iso DptB[KO] | 0.47307 | 1.60491 | 0.52709 | 0.898 | 0.3694 | |
| GtypeH. iso DptSK1 | 0.60537 | 1.83192 | 0.50765 | 1.192 | 0.2331 | |
| GtypeI. iso Dro, Att | 0.73441 | 2.08426 | 0.50773 | 1.446 | 0.1480 | |
| GtypeJ. iso Dro, Dpt | 0.04164 | 1.04252 | 0.55637 | 0.075 | 0.9403 | |
| GtypeK. iso GrB | -0.01631 | 0.98382 | 0.57739 | -0.028 | 0.9775 | |
| GtypeL. iso Rel | 0.65453 | 1.92424 | 0.51649 | 1.267 | 0.2051 | |
| GtypeM. AMP14 | 0.20061 | 1.22214 | 0.55639 | 0.361 | 0.7184 | |
| GtypeO. AMP8 | 0.31294 | 1.36744 | 0.54007 | 0.579 | 0.5623 | |
| GtypeP. AMP8+DptSK1 | 0.70345 | 2.02072 | 0.50763 | 1.386 | 0.1658 | |
