## Supplementary Data Files for "Ecology-relevant bacteria drive the evolution of host antimicrobial peptides in Drosophila": Apomorum_10-12-2021.pdf (from another version of this preprint: version 2, 2022-12-29)

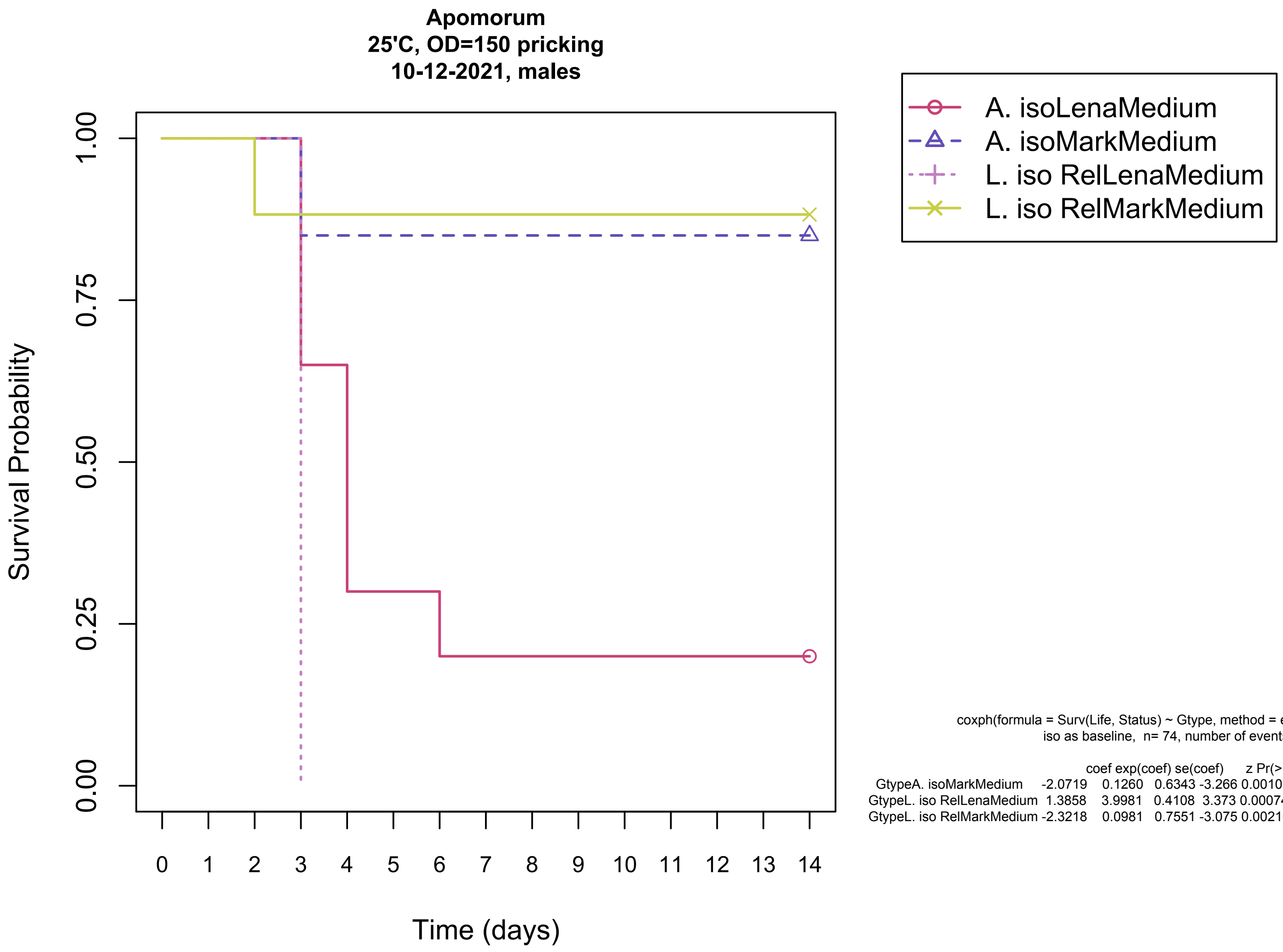


coxph(formula = Surv(Life, Status) ~ Gtype, method = efron)
iso as baseline,  n= 74, number of events= 38

| | coef | exp(coef) | se(coef) | z | Pr(>\|z\|) | |
|---|---|---|---|---|---|---|
| GtypeA. isoMarkMedium | -2.0719 | 0.1260 | 0.6343 | -3.266 | 0.001089 | ** |
| GtypeL. iso RelLenaMedium | 1.3858 | 3.9981 | 0.4108 | 3.373 | 0.000742 | *** |
| GtypeL. iso RelMarkMedium | -2.3218 | 0.0981 | 0.7551 | -3.075 | 0.002107 | ** |
