## Supplementary Data Files for "Ecology-relevant bacteria drive the evolution of host antimicrobial peptides in Drosophila": Apomorum_12-11-2021.pdf (from another version of this preprint: version 2, 2022-12-29)

**Apomorum**
**25'C, OD=150 pricking**
**12-11-2021, males**

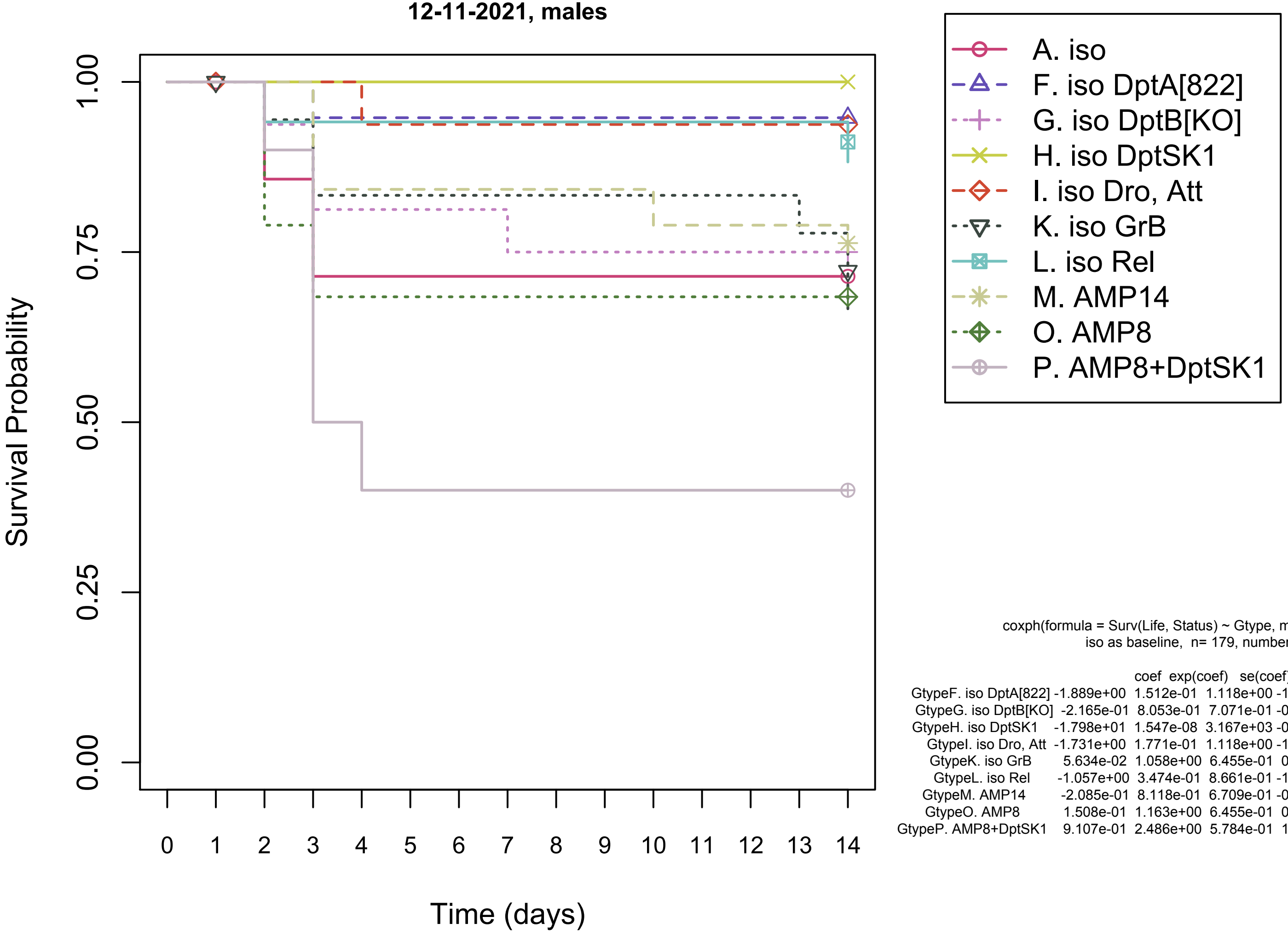


coxph(formula = Surv(Life, Status) ~ Gtype, method = efron)
iso as baseline, n= 179, number of events= 41

| | coef | exp(coef) | se(coef) | z | Pr(>\|z\|) | |
|---|---|---|---|---|---|---|
| GtypeF. iso DptA[822] | -1.889e+00 | 1.512e-01 | 1.118e+00 | -1.690 | 0.0911 | . |
| GtypeG. iso DptB[KO] | -2.165e-01 | 8.053e-01 | 7.071e-01 | -0.306 | 0.7595 | |
| GtypeH. iso DptSK1 | -1.798e+01 | 1.547e-08 | 3.167e+03 | -0.006 | 0.9955 | |
| GtypeI. iso Dro, Att | -1.731e+00 | 1.771e-01 | 1.118e+00 | -1.548 | 0.1216 | |
| GtypeK. iso GrB | 5.634e-02 | 1.058e+00 | 6.455e-01 | 0.087 | 0.9305 | |
| GtypeL. iso Rel | -1.057e+00 | 3.474e-01 | 8.661e-01 | -1.221 | 0.2222 | |
| GtypeM. AMP14 | -2.085e-01 | 8.118e-01 | 6.709e-01 | -0.311 | 0.7560 | |
| GtypeO. AMP8 | 1.508e-01 | 1.163e+00 | 6.455e-01 | 0.234 | 0.8152 | |
| GtypeP. AMP8+DptSK1 | 9.107e-01 | 2.486e+00 | 5.784e-01 | 1.575 | 0.1153 | |
