## Supplementary Data Files for "Ecology-relevant bacteria drive the evolution of host antimicrobial peptides in Drosophila": Aceto_05-10-2021.pdf (from another version of this preprint: version 2, 2022-12-29)

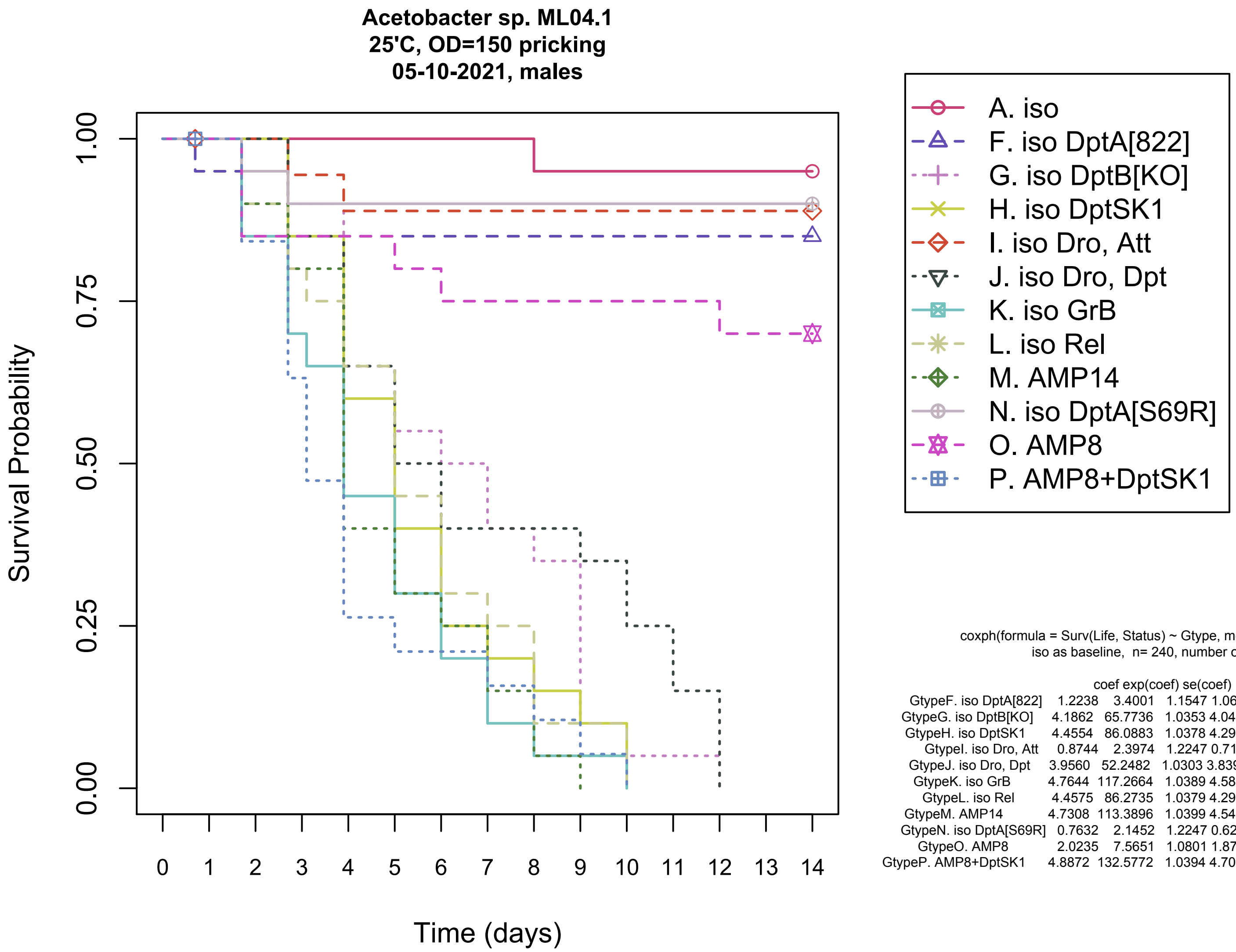


coxph(formula = Surv(Life, Status) ~ Gtype, method = efron)
iso as baseline, n= 240, number of events= 153

| | coef | exp(coef) | se(coef) | z | Pr(>\|z\|) | |
|---|---|---|---|---|---|---|
| GtypeF. iso DptA[822] | 1.2238 | 3.4001 | 1.1547 | 1.060 | 0.289213 | |
| GtypeG. iso DptB[KO] | 4.1862 | 65.7736 | 1.0353 | 4.043 | 5.27e-05 | *** |
| GtypeH. iso DptSK1 | 4.4554 | 86.0883 | 1.0378 | 4.293 | 1.76e-05 | *** |
| GtypeI. iso Dro, Att | 0.8744 | 2.3974 | 1.2247 | 0.714 | 0.475277 | |
| GtypeJ. iso Dro, Dpt | 3.9560 | 52.2482 | 1.0303 | 3.839 | 0.000123 | *** |
| GtypeK. iso GrB | 4.7644 | 117.2664 | 1.0389 | 4.586 | 4.52e-06 | *** |
| GtypeL. iso Rel | 4.4575 | 86.2735 | 1.0379 | 4.295 | 1.75e-05 | *** |
| GtypeM. AMP14 | 4.7308 | 113.3896 | 1.0399 | 4.549 | 5.38e-06 | *** |
| GtypeN. iso DptA[S69R] | 0.7632 | 2.1452 | 1.2247 | 0.623 | 0.533171 | |
| GtypeO. AMP8 | 2.0235 | 7.5651 | 1.0801 | 1.873 | 0.061011 | . |
| GtypeP. AMP8+DptSK1 | 4.8872 | 132.5772 | 1.0394 | 4.702 | 2.58e-06 | *** |
