## Supplementary Data Files for "Ecology-relevant bacteria drive the evolution of host antimicrobial peptides in Drosophila": Aceto_07-09-2021.pdf (from another version of this preprint: version 2, 2022-12-29)

**Acetobacter sp. ML04.1**
**25'C, OD=150 pricking**
**07-09-2021, males**

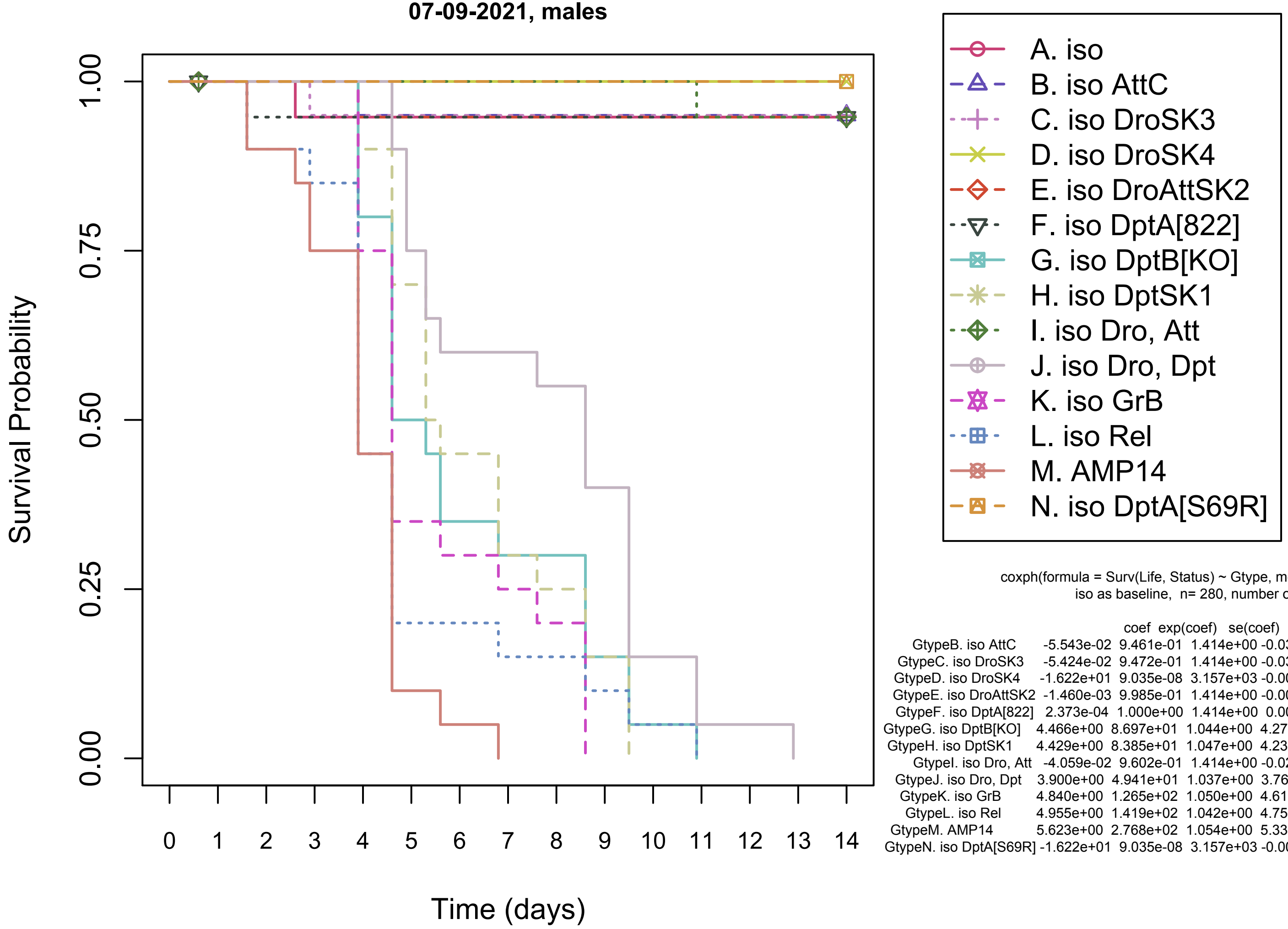


coxph(formula = Surv(Life, Status) ~ Gtype, method = efron)
iso as baseline, n= 280, number of events= 126

| | coef | exp(coef) | se(coef) | z | Pr(>\|z\|) | |
|---|---|---|---|---|---|---|
| GtypeB. iso AttC | -5.543e-02 | 9.461e-01 | 1.414e+00 | -0.039 | 0.96874 | |
| GtypeC. iso DroSK3 | -5.424e-02 | 9.472e-01 | 1.414e+00 | -0.038 | 0.96940 | |
| GtypeD. iso DroSK4 | -1.622e+01 | 9.035e-08 | 3.157e+03 | -0.005 | 0.99590 | |
| GtypeE. iso DroAttSK2 | -1.460e-03 | 9.985e-01 | 1.414e+00 | -0.001 | 0.99918 | |
| GtypeF. iso DptA[822] | 2.373e-04 | 1.000e+00 | 1.414e+00 | 0.000 | 0.99987 | |
| GtypeG. iso DptB[KO] | 4.466e+00 | 8.697e+01 | 1.044e+00 | 4.279 | 1.88e-05 | *** |
| GtypeH. iso DptSK1 | 4.429e+00 | 8.385e+01 | 1.047e+00 | 4.230 | 2.34e-05 | *** |
| GtypeI. iso Dro, Att | -4.059e-02 | 9.602e-01 | 1.414e+00 | -0.029 | 0.97710 | |
| GtypeJ. iso Dro, Dpt | 3.900e+00 | 4.941e+01 | 1.037e+00 | 3.760 | 0.00017 | *** |
| GtypeK. iso GrB | 4.840e+00 | 1.265e+02 | 1.050e+00 | 4.610 | 4.03e-06 | *** |
| GtypeL. iso Rel | 4.955e+00 | 1.419e+02 | 1.042e+00 | 4.755 | 1.99e-06 | *** |
| GtypeM. AMP14 | 5.623e+00 | 2.768e+02 | 1.054e+00 | 5.336 | 9.52e-08 | *** |
| GtypeN. iso DptA[S69R] | -1.622e+01 | 9.035e-08 | 3.157e+03 | -0.005 | 0.99590 | |
