## Supplementary Data Files for "Ecology-relevant bacteria drive the evolution of host antimicrobial peptides in Drosophila": Aceto_17-09-2021.pdf (from another version of this preprint: version 2, 2022-12-29)

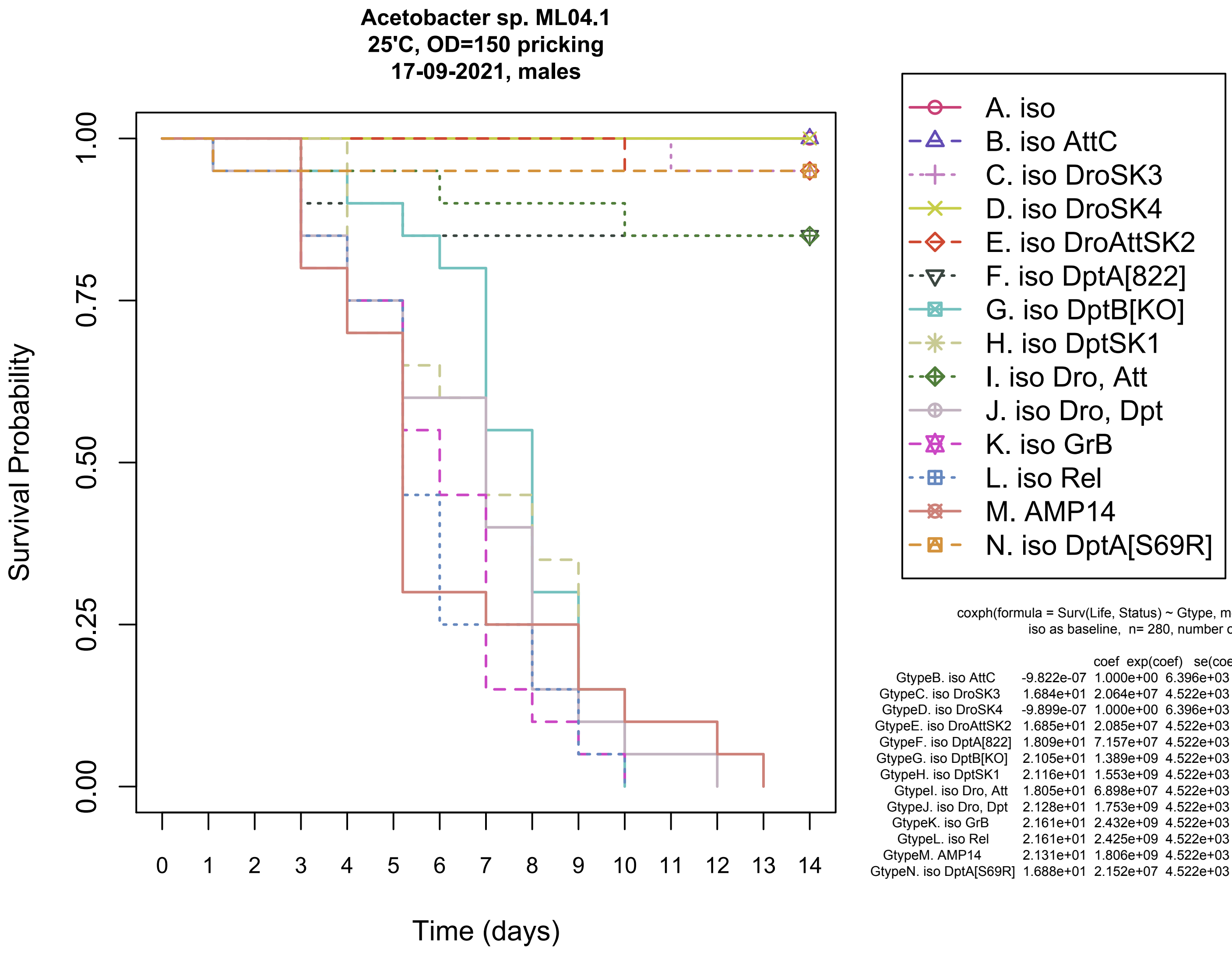


coxph(formula = Surv(Life, Status) ~ Gtype, method = efron)
iso as baseline, n= 280, number of events= 129

| | coef | exp(coef) | se(coef) | z | Pr(>\|z\|) |
|---|---|---|---|---|---|
| GtypeB. iso AttC | -9.822e-07 | 1.000e+00 | 6.396e+03 | 0.000 | 1.000 |
| GtypeC. iso DroSK3 | 1.684e+01 | 2.064e+07 | 4.522e+03 | 0.004 | 0.997 |
| GtypeD. iso DroSK4 | -9.899e-07 | 1.000e+00 | 6.396e+03 | 0.000 | 1.000 |
| GtypeE. iso DroAttSK2 | 1.685e+01 | 2.085e+07 | 4.522e+03 | 0.004 | 0.997 |
| GtypeF. iso DptA[822] | 1.809e+01 | 7.157e+07 | 4.522e+03 | 0.004 | 0.997 |
| GtypeG. iso DptB[KO] | 2.105e+01 | 1.389e+09 | 4.522e+03 | 0.005 | 0.996 |
| GtypeH. iso DptSK1 | 2.116e+01 | 1.553e+09 | 4.522e+03 | 0.005 | 0.996 |
| GtypeI. iso Dro, Att | 1.805e+01 | 6.898e+07 | 4.522e+03 | 0.004 | 0.997 |
| GtypeJ. iso Dro, Dpt | 2.128e+01 | 1.753e+09 | 4.522e+03 | 0.005 | 0.996 |
| GtypeK. iso GrB | 2.161e+01 | 2.432e+09 | 4.522e+03 | 0.005 | 0.996 |
| GtypeL. iso Rel | 2.161e+01 | 2.425e+09 | 4.522e+03 | 0.005 | 0.996 |
| GtypeM. AMP14 | 2.131e+01 | 1.806e+09 | 4.522e+03 | 0.005 | 0.996 |
| GtypeN. iso DptA[S69R] | 1.688e+01 | 2.152e+07 | 4.522e+03 | 0.004 | 0.997 |
