## Supplementary Data Files for "Ecology-relevant bacteria drive the evolution of host antimicrobial peptides in Drosophila": Aceto_23-11-2021.pdf (from another version of this preprint: version 2, 2022-12-29)

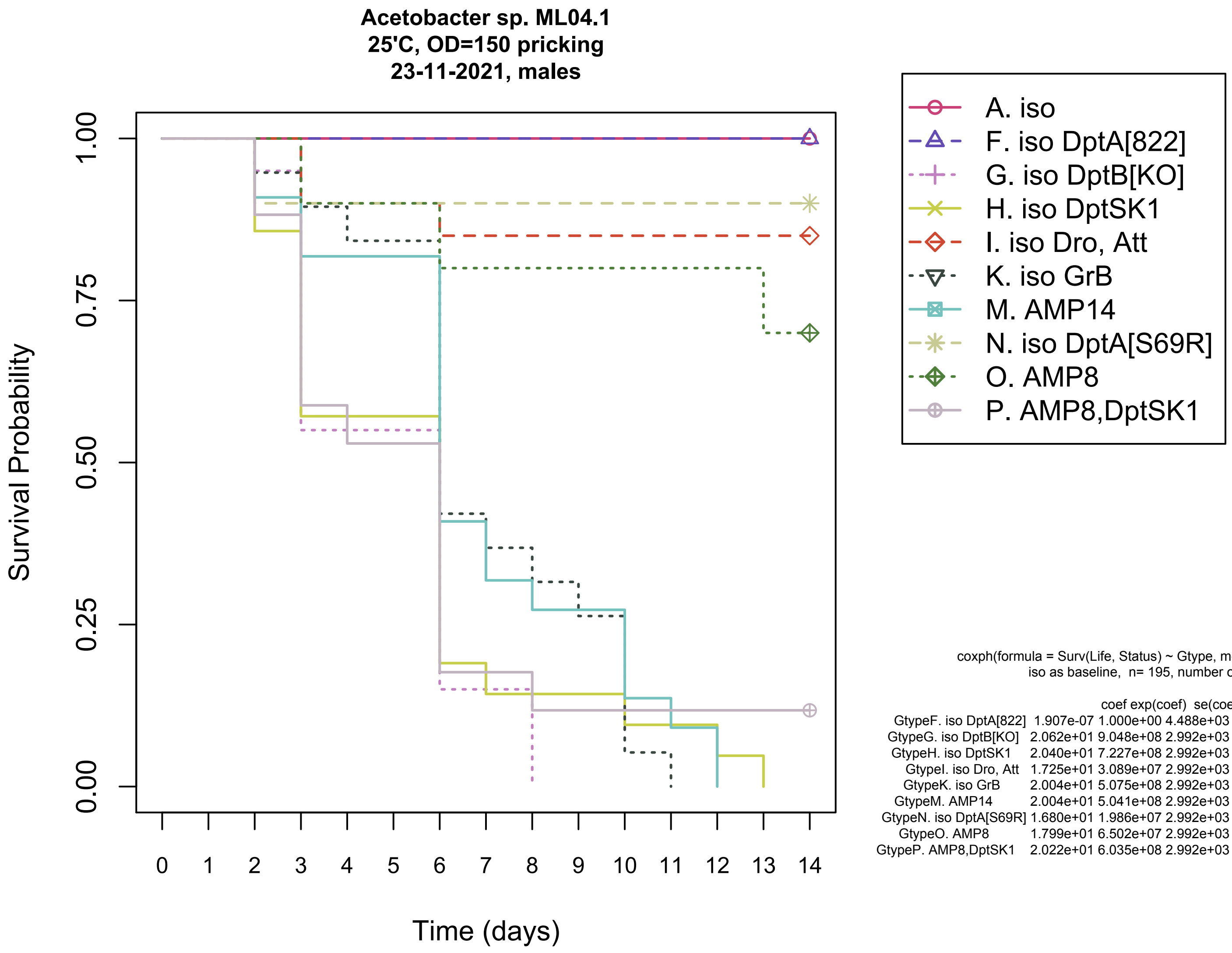


coxph(formula = Surv(Life, Status) ~ Gtype, method = efron)
iso as baseline,  n= 195, number of events= 108

| | coef | exp(coef) | se(coef) | z | Pr(>\|z\|) |
|---|---|---|---|---|---|
| GtypeF. iso DptA[822] | 1.907e-07 | 1.000e+00 | 4.488e+03 | 0.000 | 1.000 |
| GtypeG. iso DptB[KO] | 2.062e+01 | 9.048e+08 | 2.992e+03 | 0.007 | 0.995 |
| GtypeH. iso DptSK1 | 2.040e+01 | 7.227e+08 | 2.992e+03 | 0.007 | 0.995 |
| GtypeI. iso Dro, Att | 1.725e+01 | 3.089e+07 | 2.992e+03 | 0.006 | 0.995 |
| GtypeK. iso GrB | 2.004e+01 | 5.075e+08 | 2.992e+03 | 0.007 | 0.995 |
| GtypeM. AMP14 | 2.004e+01 | 5.041e+08 | 2.992e+03 | 0.007 | 0.995 |
| GtypeN. iso DptA[S69R] | 1.680e+01 | 1.986e+07 | 2.992e+03 | 0.006 | 0.996 |
| GtypeO. AMP8 | 1.799e+01 | 6.502e+07 | 2.992e+03 | 0.006 | 0.995 |
| GtypeP. AMP8,DptSK1 | 2.022e+01 | 6.035e+08 | 2.992e+03 | 0.007 | 0.995 |
