## Supplementary Data Files for "Ecology-relevant bacteria drive the evolution of host antimicrobial peptides in Drosophila": Aceto_CoxPH.pdf (from another version of this preprint: version 2, 2022-12-29)

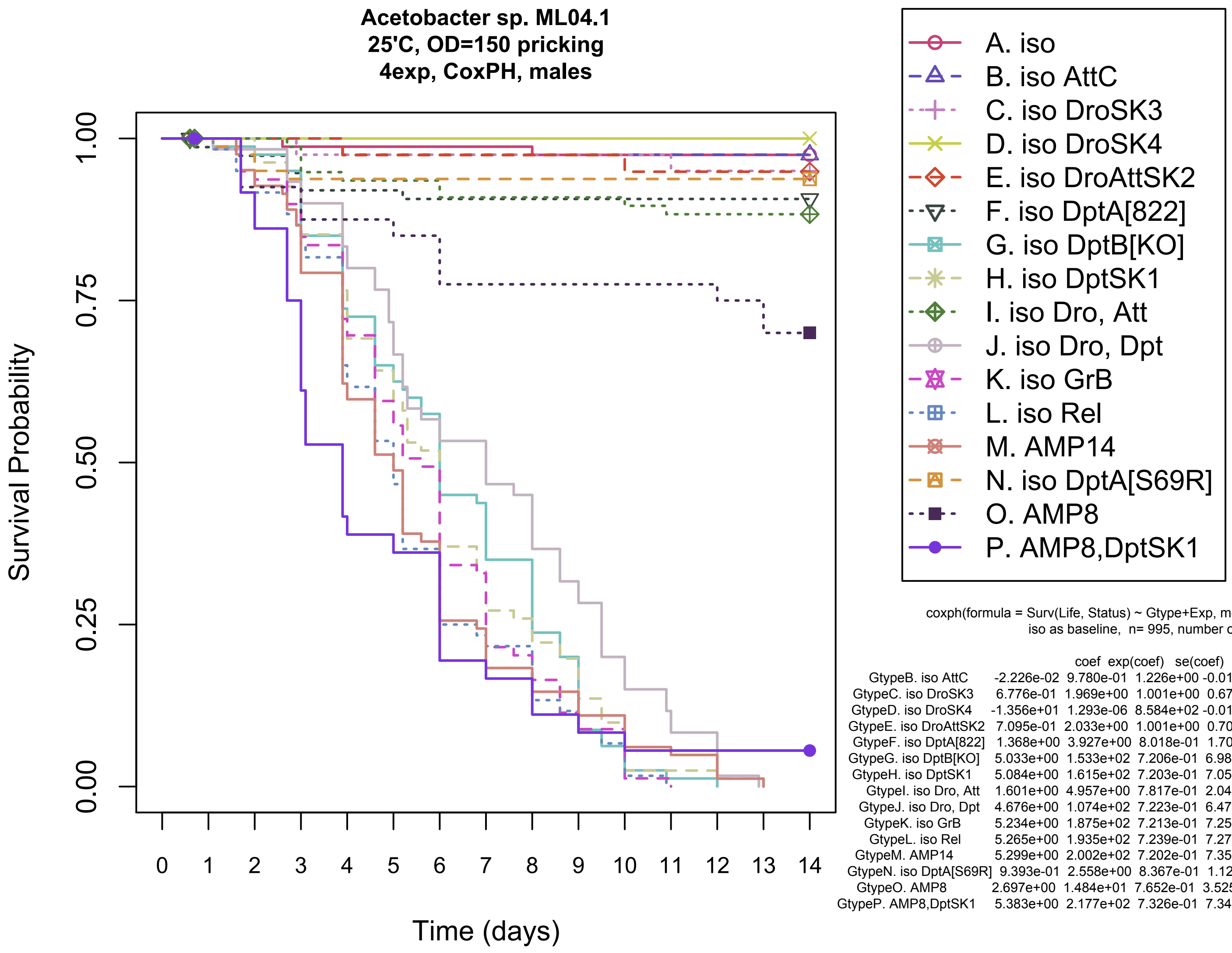


coxph(formula = Surv(Life, Status) ~ Gtype+Exp, method = efron)
iso as baseline, n= 995, number of events= 516

| | coef | exp(coef) | se(coef) | z | Pr(>\|z\|) | |
|---|---|---|---|---|---|---|
| GtypeB. iso AttC | -2.226e-02 | 9.780e-01 | 1.226e+00 | -0.018 | 0.985508 | |
| GtypeC. iso DroSK3 | 6.776e-01 | 1.969e+00 | 1.001e+00 | 0.677 | 0.498523 | |
| GtypeD. iso DroSK4 | -1.356e+01 | 1.293e-06 | 8.584e+02 | -0.016 | 0.987397 | |
| GtypeE. iso DroAttSK2 | 7.095e-01 | 2.033e+00 | 1.001e+00 | 0.709 | 0.478497 | |
| GtypeF. iso DptA[822] | 1.368e+00 | 3.927e+00 | 8.018e-01 | 1.706 | 0.088001 | . |
| GtypeG. iso DptB[KO] | 5.033e+00 | 1.533e+02 | 7.206e-01 | 6.984 | 2.87e-12 | *** |
| GtypeH. iso DptSK1 | 5.084e+00 | 1.615e+02 | 7.203e-01 | 7.059 | 1.68e-12 | *** |
| GtypeI. iso Dro, Att | 1.601e+00 | 4.957e+00 | 7.817e-01 | 2.048 | 0.040599 | * |
| GtypeJ. iso Dro, Dpt | 4.676e+00 | 1.074e+02 | 7.223e-01 | 6.474 | 9.51e-11 | *** |
| GtypeK. iso GrB | 5.234e+00 | 1.875e+02 | 7.213e-01 | 7.256 | 3.98e-13 | *** |
| GtypeL. iso Rel | 5.265e+00 | 1.935e+02 | 7.239e-01 | 7.274 | 3.49e-13 | *** |
| GtypeM. AMP14 | 5.299e+00 | 2.002e+02 | 7.202e-01 | 7.358 | 1.87e-13 | *** |
| GtypeN. iso DptA[S69R] | 9.393e-01 | 2.558e+00 | 8.367e-01 | 1.123 | 0.261561 | |
| GtypeO. AMP8 | 2.697e+00 | 1.484e+01 | 7.652e-01 | 3.525 | 0.000424 | *** |
| GtypeP. AMP8,DptSK1 | 5.383e+00 | 2.177e+02 | 7.326e-01 | 7.347 | 2.02e-13 | *** |
| exp | | | | | | ** |
