## Supplementary Data Files for "Ecology-relevant bacteria drive the evolution of host antimicrobial peptides in Drosophila": ML04-1_Monoassoc.pdf (from another version of this preprint: version 2, 2022-12-29)

**Longevity at 25'C**
**Antibiotic-reared flies exposed to Asicerae ML04.1 OD100 twice (Day 0, Day 14)**
**i.e. 'monoassociation'**

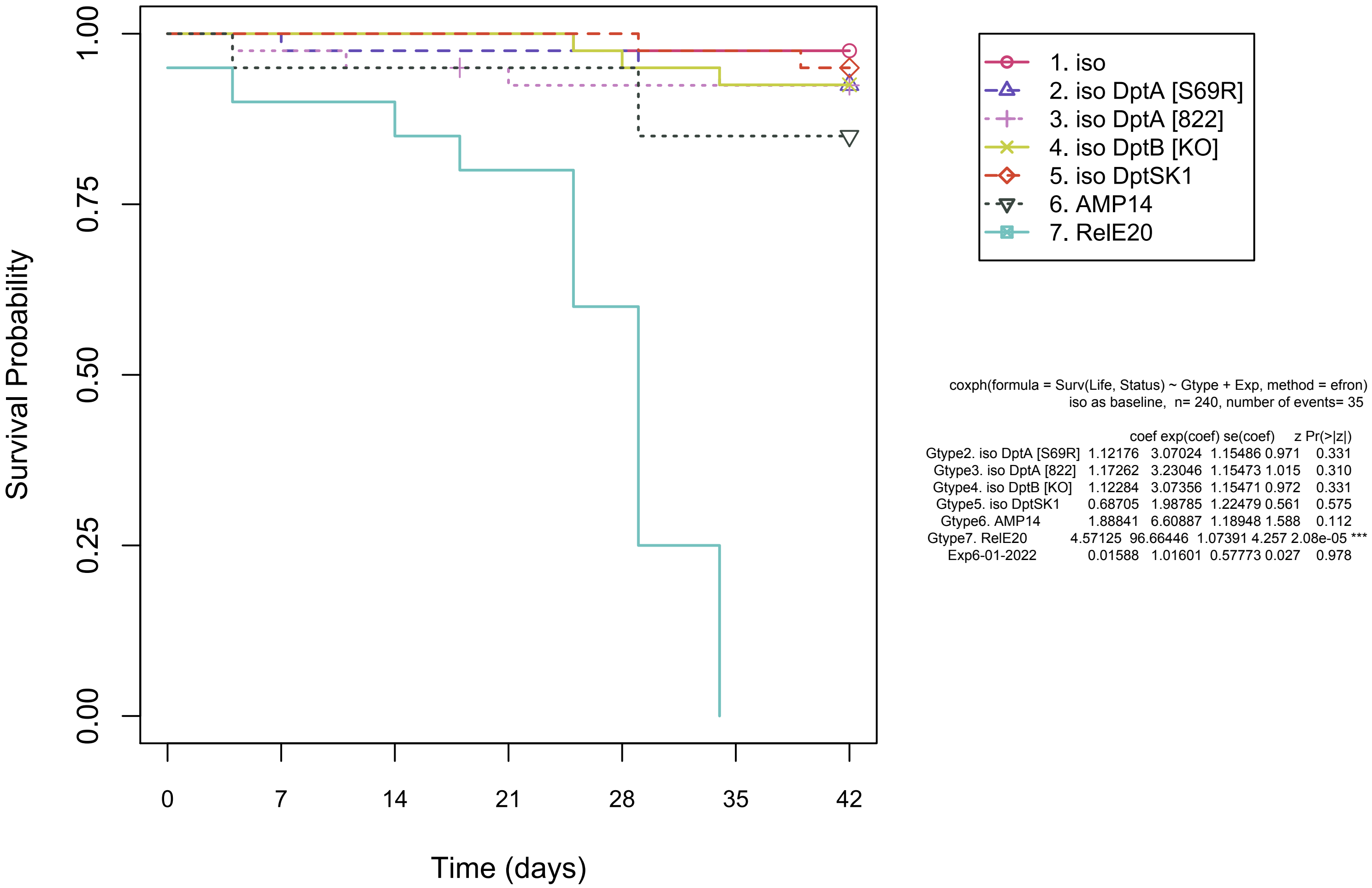
