## Supplementary Data Files for "Ecology-relevant bacteria drive the evolution of host antimicrobial peptides in Drosophila": Atrop_26-01-2022.pdf (from another version of this preprint: version 2, 2022-12-29)

**Acetobacter sp. ML04.1**
**25'C, OD=150 pricking**
**26-01-2022, males**

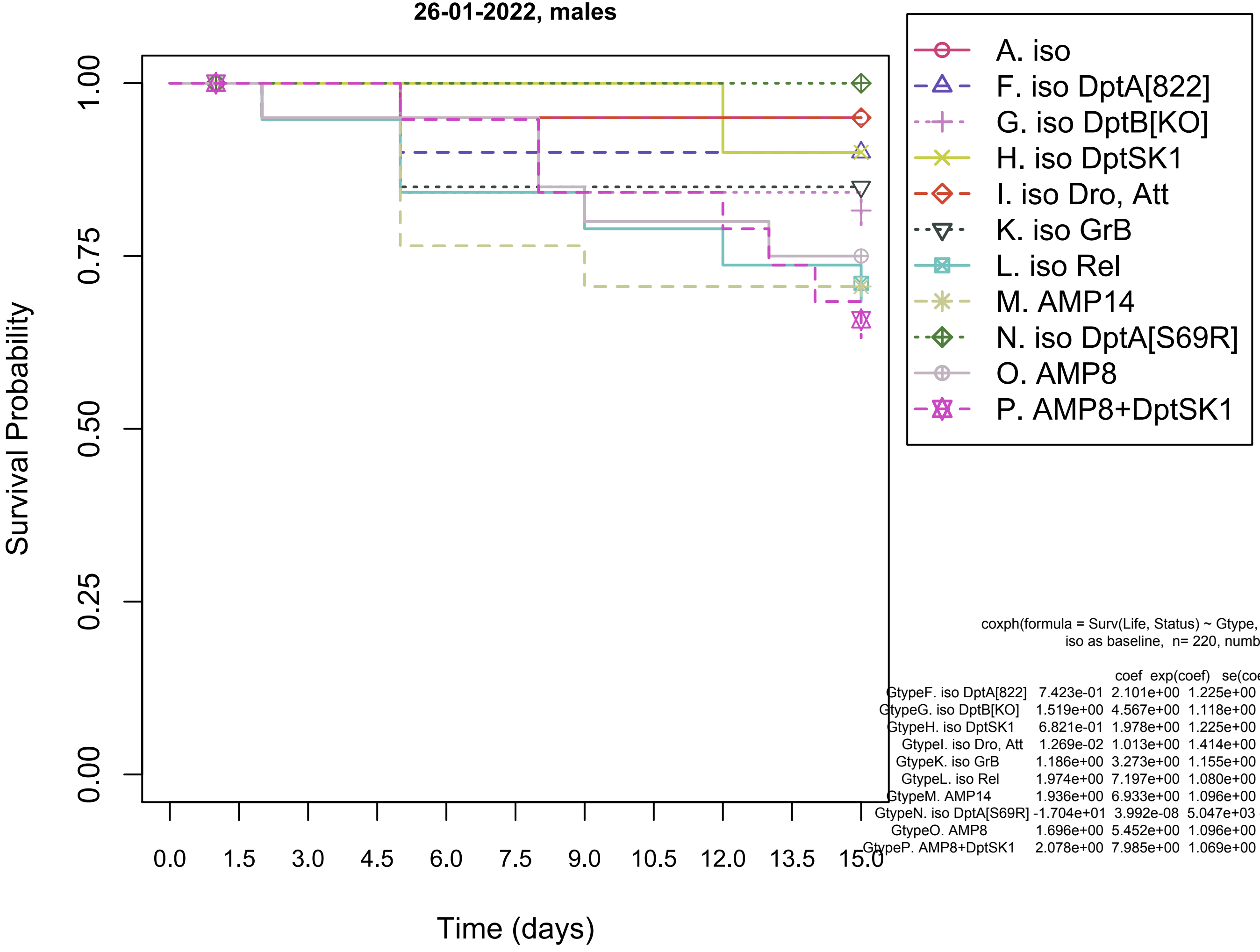


coxph(formula = Surv(Life, Status) ~ Gtype, method = efron)
iso as baseline, n= 220, number of events= 36

| | coef | exp(coef) | se(coef) | z | Pr(>\|z\|) | |
|---|---|---|---|---|---|---|
| GtypeF. iso DptA[822] | 7.423e-01 | 2.101e+00 | 1.225e+00 | 0.606 | 0.5445 | |
| GtypeG. iso DptB[KO] | 1.519e+00 | 4.567e+00 | 1.118e+00 | 1.358 | 0.1743 | |
| GtypeH. iso DptSK1 | 6.821e-01 | 1.978e+00 | 1.225e+00 | 0.557 | 0.5776 | |
| GtypeI. iso Dro, Att | 1.269e-02 | 1.013e+00 | 1.414e+00 | 0.009 | 0.9928 | |
| GtypeK. iso GrB | 1.186e+00 | 3.273e+00 | 1.155e+00 | 1.027 | 0.3046 | |
| GtypeL. iso Rel | 1.974e+00 | 7.197e+00 | 1.080e+00 | 1.827 | 0.0677 | . |
| GtypeM. AMP14 | 1.936e+00 | 6.933e+00 | 1.096e+00 | 1.767 | 0.0772 | . |
| GtypeN. iso DptA[S69R] | -1.704e+01 | 3.992e-08 | 5.047e+03 | -0.003 | 0.9973 | |
| GtypeO. AMP8 | 1.696e+00 | 5.452e+00 | 1.096e+00 | 1.548 | 0.1216 | |
| GtypeP. AMP8+DptSK1 | 2.078e+00 | 7.985e+00 | 1.069e+00 | 1.943 | 0.0520 | . |
