## Supplementary Data Files for "Ecology-relevant bacteria drive the evolution of host antimicrobial peptides in Drosophila": Atrop_26-11-2021.pdf (from another version of this preprint: version 2, 2022-12-29)

**Acetobacter sp. ML04.1**
**25'C, OD=150 pricking**
**26-11-2021, males**

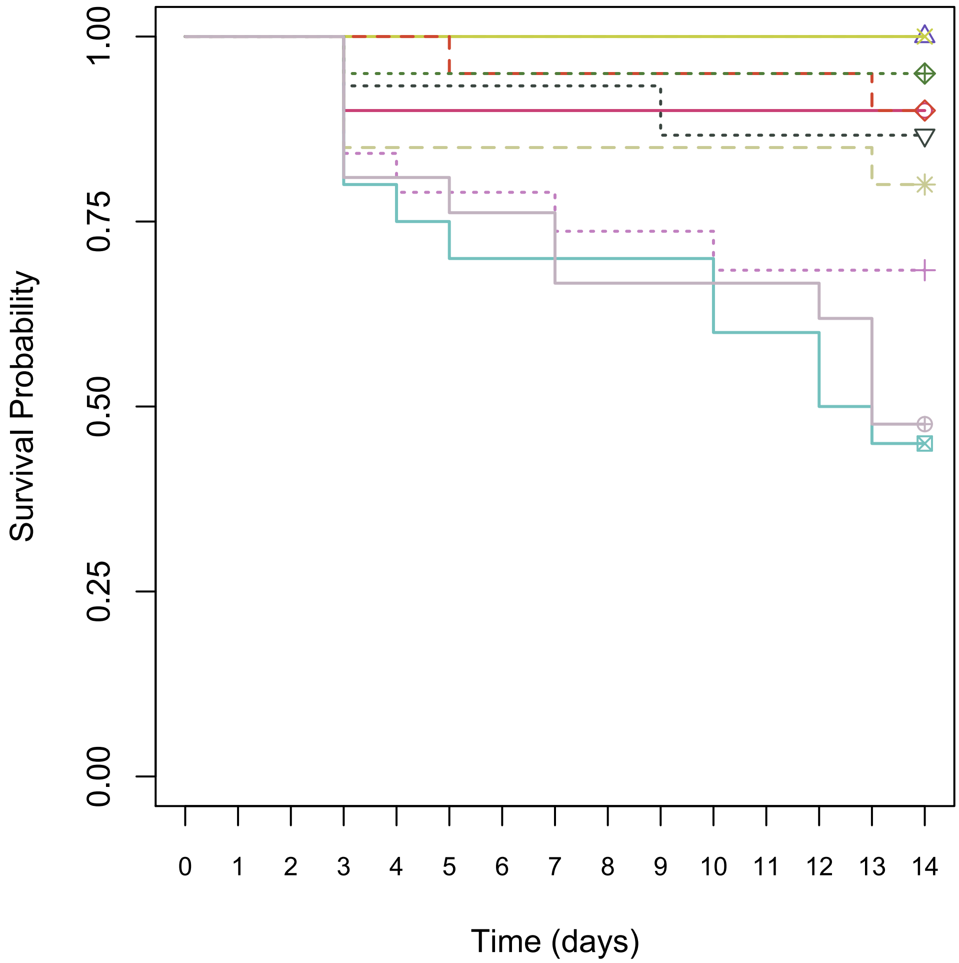


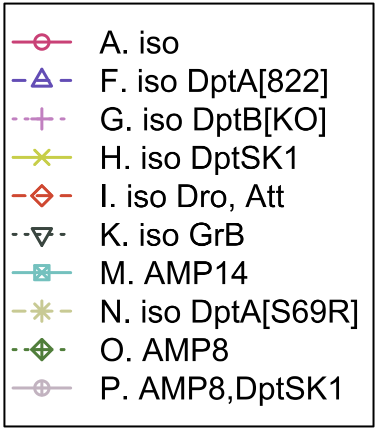


A BIT WEIRD THAT DptSK1 SURVIVED BUT
AMP14, AMP8,DptSK1, DptBKO, ALL DIED
WITH SIMILAR KINETICS OVER FIRST ~10 DAYS.

DOUBLE CHECK DptSK1 IN FUTURE EXPERIMENTS.

coxph(formula = Surv(Life, Status) ~ Gtype, method = efron)
iso as baseline,  n= 195, number of events= 39

| | coef | exp(coef) | se(coef) | z | Pr(>\|z\|) | |
|---|---|---|---|---|---|---|
| GtypeF. iso DptA[822] | -1.823e+01 | 1.216e-08 | 6.152e+03 | -0.003 | 0.9976 | |
| GtypeG. iso DptB[KO] | 1.282e+00 | 3.606e+00 | 8.167e-01 | 1.570 | 0.1163 | |
| GtypeH. iso DptSK1 | -1.823e+01 | 1.216e-08 | 6.152e+03 | -0.003 | 0.9976 | |
| GtypeI. iso Dro, Att | -5.326e-02 | 9.481e-01 | 1.000e+00 | -0.053 | 0.9575 | |
| GtypeK. iso GrB | 2.855e-01 | 1.330e+00 | 1.000e+00 | 0.286 | 0.7753 | |
| GtypeM. AMP14 | 1.937e+00 | 6.935e+00 | 7.695e-01 | 2.517 | 0.0118 | * |
| GtypeN. iso DptA[S69R] | 7.420e-01 | 2.100e+00 | 8.660e-01 | 0.857 | 0.3916 | |
| GtypeO. AMP8 | -7.354e-01 | 4.793e-01 | 1.225e+00 | -0.600 | 0.5482 | |
| GtypeP. AMP8,DptSK1 | 1.845e+00 | 6.327e+00 | 7.692e-01 | 2.398 | 0.0165 | * |
