## Supplementary Data Files for "Ecology-relevant bacteria drive the evolution of host antimicrobial peptides in Drosophila": Atrop_CoxPH.pdf (from another version of this preprint: version 2, 2022-12-29)

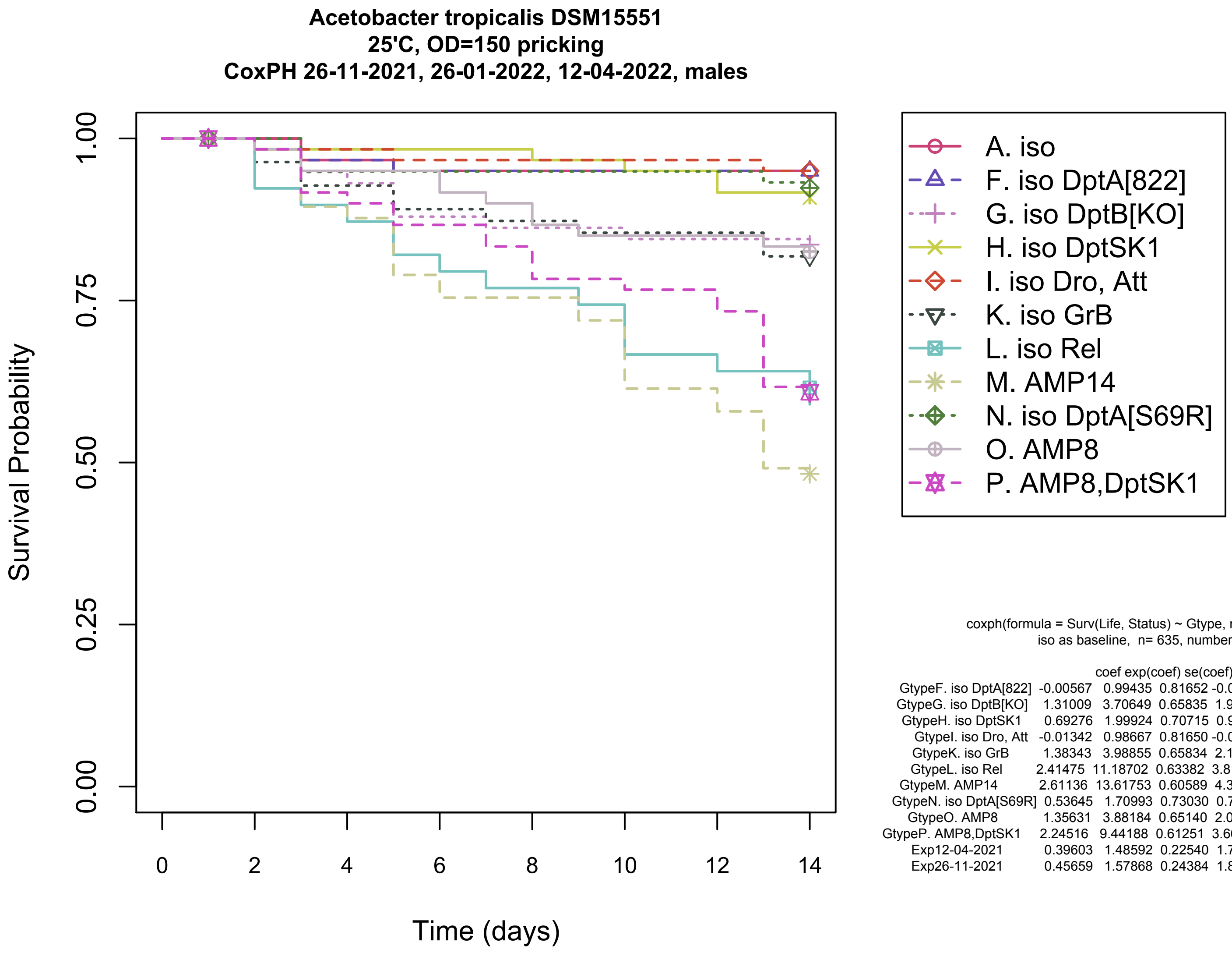


coxph(formula = Surv(Life, Status) ~ Gtype, method = efron)
iso as baseline, n= 635, number of events= 121

| | coef | exp(coef) | se(coef) | z | Pr(>\|z\|) | |
|---|---|---|---|---|---|---|
| GtypeF. iso DptA[822] | -0.00567 | 0.99435 | 0.81652 | -0.007 | 0.994460 | |
| GtypeG. iso DptB[KO] | 1.31009 | 3.70649 | 0.65835 | 1.990 | 0.046596 | * |
| GtypeH. iso DptSK1 | 0.69276 | 1.99924 | 0.70715 | 0.980 | 0.327256 | |
| GtypeI. iso Dro, Att | -0.01342 | 0.98667 | 0.81650 | -0.016 | 0.986887 | |
| GtypeK. iso GrB | 1.38343 | 3.98855 | 0.65834 | 2.101 | 0.035607 | * |
| GtypeL. iso Rel | 2.41475 | 11.18702 | 0.63382 | 3.810 | 0.000139 | *** |
| GtypeM. AMP14 | 2.61136 | 13.61753 | 0.60589 | 4.310 | 1.63e-05 | *** |
| GtypeN. iso DptA[S69R] | 0.53645 | 1.70993 | 0.73030 | 0.735 | 0.462606 | |
| GtypeO. AMP8 | 1.35631 | 3.88184 | 0.65140 | 2.082 | 0.037330 | * |
| GtypeP. AMP8,DptSK1 | 2.24516 | 9.44188 | 0.61251 | 3.665 | 0.000247 | *** |
| Exp12-04-2021 | 0.39603 | 1.48592 | 0.22540 | 1.757 | 0.078917 | . |
| Exp26-11-2021 | 0.45659 | 1.57868 | 0.24384 | 1.872 | 0.061143 | . |
