## Supplementary Data Files for "Ecology-relevant bacteria drive the evolution of host antimicrobial peptides in Drosophila": Prett_DptB-IR-CoxPH-2021.pdf (from another version of this preprint: version 2, 2022-12-29)

**P. rettgeri 25'C, OD=1.0 pricking, DptB-IR**
**CoxPH-2021, 6exp (3Mark + 3Lena with alt AMP ctrls), MALES**
**Note: Mark exp generally killed more than Lena exp,**
**perhaps why Lena's triplicate finds no effect of DptB-IR and Mark found small effect.**
**Prefer to merge both datasets.**

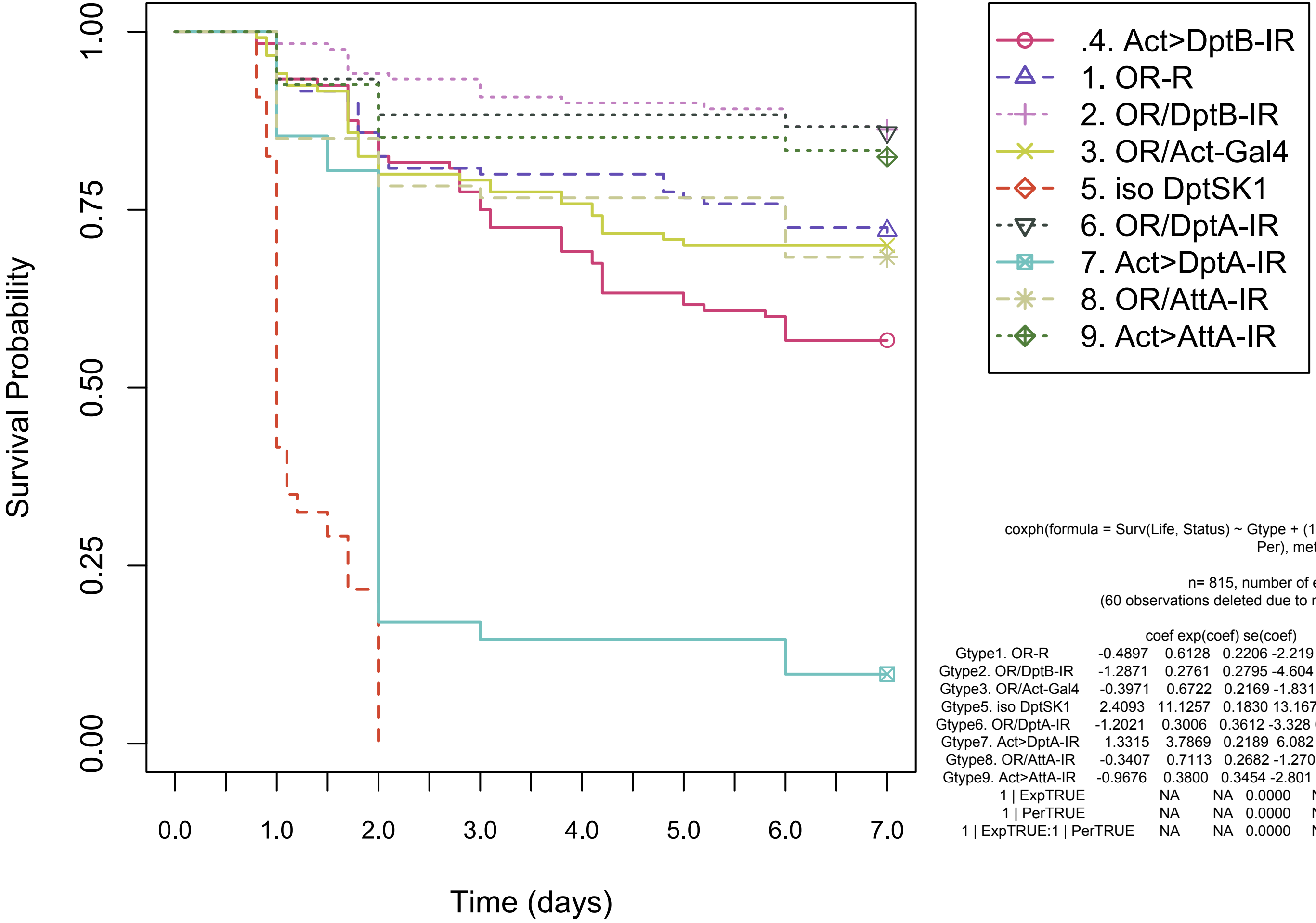


coxph(formula = Surv(Life, Status) ~ Gtype + (1 | Exp) * (1 | Per), method = efron)

n= 815, number of events= 334
(60 observations deleted due to missingness)

| | coef | exp(coef) | se(coef) | z | Pr(>\|z\|) | |
|---|---|---|---|---|---|---|
| Gtype1. OR-R | -0.4897 | 0.6128 | 0.2206 | -2.219 | 0.026470 | * |
| Gtype2. OR/DptB-IR | -1.2871 | 0.2761 | 0.2795 | -4.604 | 4.14e-06 | *** |
| Gtype3. OR/Act-Gal4 | -0.3971 | 0.6722 | 0.2169 | -1.831 | 0.067080 | . |
| Gtype5. iso DptSK1 | 2.4093 | 11.1257 | 0.1830 | 13.167 | < 2e-16 | *** |
| Gtype6. OR/DptA-IR | -1.2021 | 0.3006 | 0.3612 | -3.328 | 0.000873 | *** |
| Gtype7. Act>DptA-IR | 1.3315 | 3.7869 | 0.2189 | 6.082 | 1.19e-09 | *** |
| Gtype8. OR/AttA-IR | -0.3407 | 0.7113 | 0.2682 | -1.270 | 0.203940 | |
| Gtype9. Act>AttA-IR | -0.9676 | 0.3800 | 0.3454 | -2.801 | 0.005093 | ** |
| 1 \| ExpTRUE | NA | NA | 0.0000 | NA | NA | |
| 1 \| PerTRUE | NA | NA | 0.0000 | NA | NA | |
| 1 \| ExpTRUE:1 \| PerTRUE | NA | NA | 0.0000 | NA | NA | |
