## Supplementary Data Files for "Ecology-relevant bacteria drive the evolution of host antimicrobial peptides in Drosophila": Prett_02-03-2021.pdf (from another version of this preprint: version 2, 2022-12-29)

**P. rettgeri**
**25'C, OD=0.1 pricking, 3 days only**
**02-03-2021, MALES - age not well controlled**

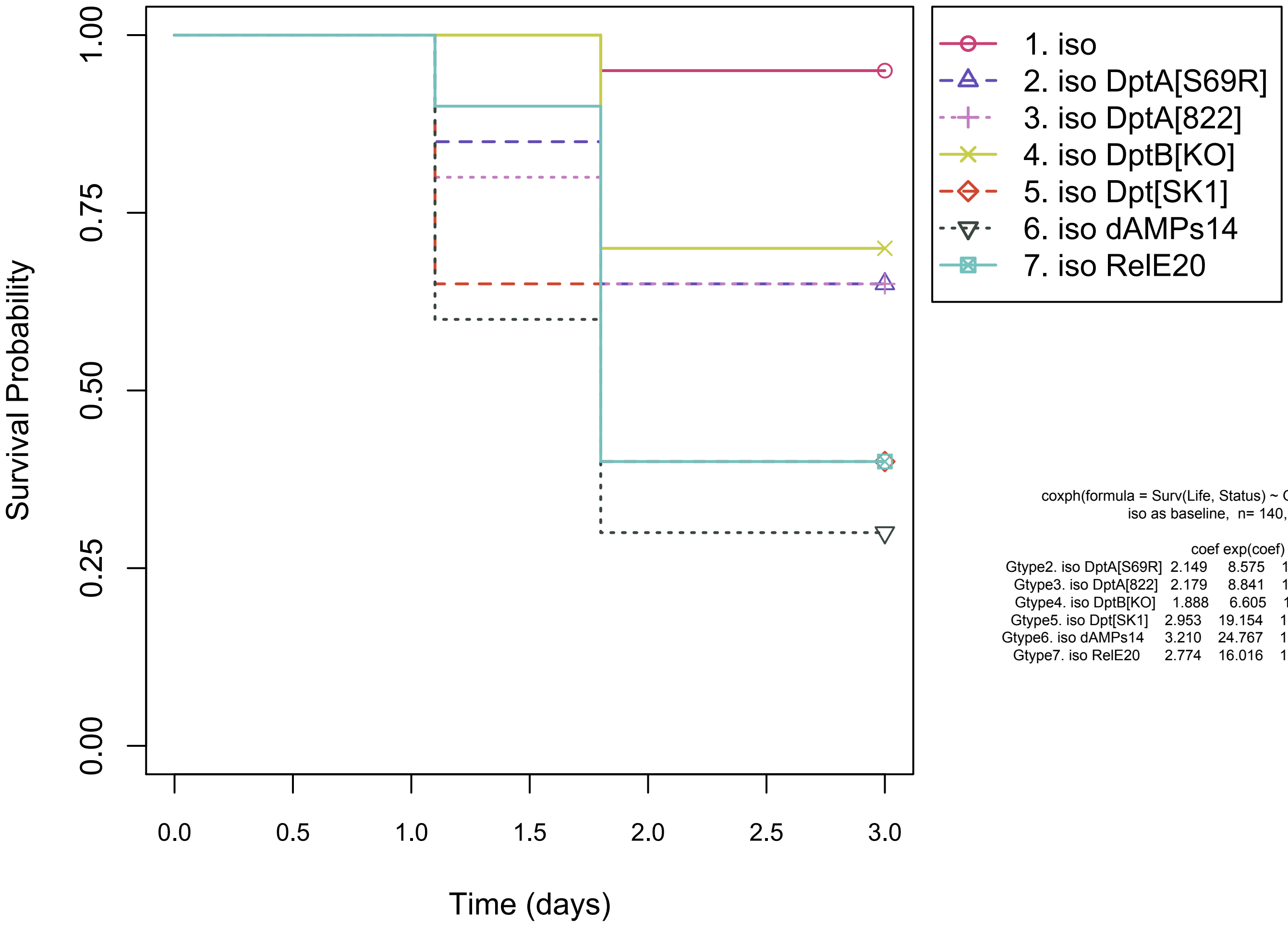


coxph(formula = Surv(Life, Status) ~ Gtype, method = efron)
iso as baseline,  n= 140, number of events= 59

| | coef | exp(coef) | se(coef) | z | Pr(>\|z\|) | |
|---|---|---|---|---|---|---|
| Gtype2. iso DptA[S69R] | 2.149 | 8.575 | 1.069 | 2.010 | 0.04445 | * |
| Gtype3. iso DptA[822] | 2.179 | 8.841 | 1.069 | 2.038 | 0.04151 | * |
| Gtype4. iso DptB[KO] | 1.888 | 6.605 | 1.080 | 1.748 | 0.08051 | . |
| Gtype5. iso Dpt[SK1] | 2.953 | 19.154 | 1.041 | 2.835 | 0.00458 | ** |
| Gtype6. iso dAMPs14 | 3.210 | 24.767 | 1.036 | 3.098 | 0.00195 | ** |
| Gtype7. iso RelE20 | 2.774 | 16.016 | 1.041 | 2.663 | 0.00773 | ** |
