## Supplementary Data Files for "Ecology-relevant bacteria drive the evolution of host antimicrobial peptides in Drosophila": Prett_05-03-2021.pdf (from another version of this preprint: version 2, 2022-12-29)

**P. rettgeri**
**25'C, OD=1.0 pricking**
**05-03-2021, MALES - Prett grown ~6h to OD=~0.9**

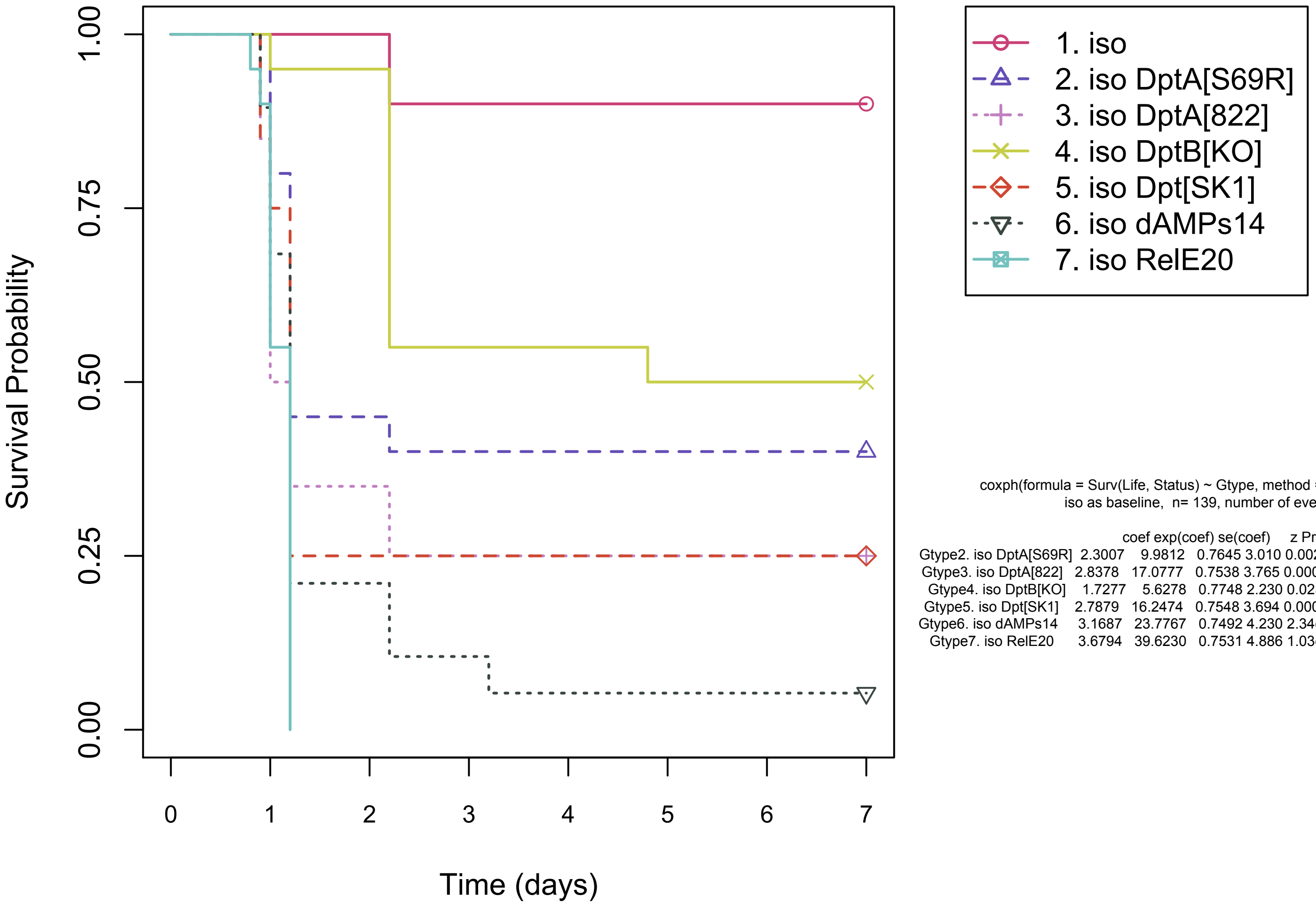


coxph(formula = Surv(Life, Status) ~ Gtype, method = efron)
iso as baseline, n= 139, number of events= 92

| | coef | exp(coef) | se(coef) | z | Pr(>\|z\|) | |
|---|---|---|---|---|---|---|
| Gtype2. iso DptA[S69R] | 2.3007 | 9.9812 | 0.7645 | 3.010 | 0.002616 | ** |
| Gtype3. iso DptA[822] | 2.8378 | 17.0777 | 0.7538 | 3.765 | 0.000167 | *** |
| Gtype4. iso DptB[KO] | 1.7277 | 5.6278 | 0.7748 | 2.230 | 0.025759 | * |
| Gtype5. iso Dpt[SK1] | 2.7879 | 16.2474 | 0.7548 | 3.694 | 0.000221 | *** |
| Gtype6. iso dAMPs14 | 3.1687 | 23.7767 | 0.7492 | 4.230 | 2.34e-05 | *** |
| Gtype7. iso RelE20 | 3.6794 | 39.6230 | 0.7531 | 4.886 | 1.03e-06 | *** |
