## Supplementary Data Files for "Ecology-relevant bacteria drive the evolution of host antimicrobial peptides in Drosophila": Prett_CoxPH-2021.pdf (from another version of this preprint: version 2, 2022-12-29)

**P. rettgeri**

**25'C, OD=1.0 pricking, CoxPH-2021, 3exp, MALES**

**Consistent in all 3 (though to *very* different extents) that dAMPs14 dies faster than DptSK1 or DptA822**

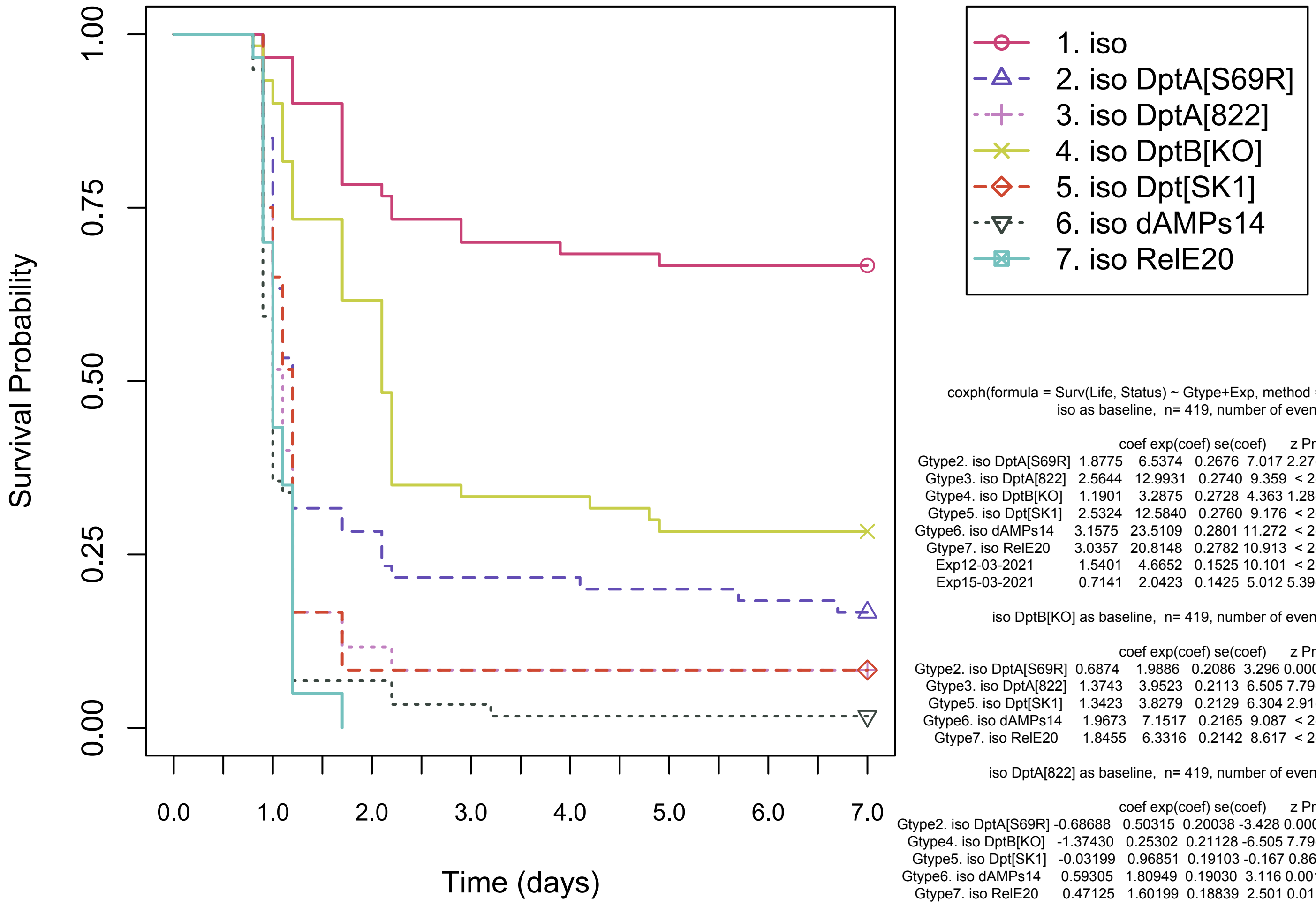


```
coxph(formula = Surv(Life, Status) ~ Gtype+Exp, method = efron)
iso as baseline,  n= 419, number of events= 341

                          coef exp(coef) se(coef)      z Pr(>|z|)
Gtype2. iso DptA[S69R]  1.8775    6.5374   0.2676  7.017 2.27e-12 ***
Gtype3. iso DptA[822]   2.5644   12.9931   0.2740  9.359  < 2e-16 ***
Gtype4. iso DptB[KO]    1.1901    3.2875   0.2728  4.363 1.28e-05 ***
Gtype5. iso Dpt[SK1]    2.5324   12.5840   0.2760  9.176  < 2e-16 ***
Gtype6. iso dAMPs14     3.1575   23.5109   0.2801 11.272  < 2e-16 ***
Gtype7. iso RelE20      3.0357   20.8148   0.2782 10.913  < 2e-16 ***
Exp12-03-2021           1.5401    4.6652   0.1525 10.101  < 2e-16 ***
Exp15-03-2021           0.7141    2.0423   0.1425  5.012 5.39e-07 ***

iso DptB[KO] as baseline,  n= 419, number of events= 341

                          coef exp(coef) se(coef)      z Pr(>|z|)
Gtype2. iso DptA[S69R]  0.6874    1.9886   0.2086  3.296 0.000982 ***
Gtype3. iso DptA[822]   1.3743    3.9523   0.2113  6.505 7.79e-11 ***
Gtype5. iso Dpt[SK1]    1.3423    3.8279   0.2129  6.304 2.91e-10 ***
Gtype6. iso dAMPs14     1.9673    7.1517   0.2165  9.087  < 2e-16 ***
Gtype7. iso RelE20      1.8455    6.3316   0.2142  8.617  < 2e-16 ***

iso DptA[822] as baseline,  n= 419, number of events= 341

                            coef exp(coef) se(coef)      z Pr(>|z|)
Gtype2. iso DptA[S69R] -0.68688   0.50315  0.20038 -3.428 0.000608 ***
Gtype4. iso DptB[KO]   -1.37430   0.25302  0.21128 -6.505 7.79e-11 ***
Gtype5. iso Dpt[SK1]   -0.03199   0.96851  0.19103 -0.167 0.867000
Gtype6. iso dAMPs14     0.59305   1.80949  0.19030  3.116 0.001831 **
Gtype7. iso RelE20      0.47125   1.60199  0.18839  2.501 0.012370 *
```
