## Supplementary figures and images for "Ecology-relevant bacteria drive the evolution of host antimicrobial peptides in Drosophila" (from another version of this preprint: version 2, 2022-12-29)

### Aceto_DptB-IR-CoxPH-2022_Apr8-2022.pdf

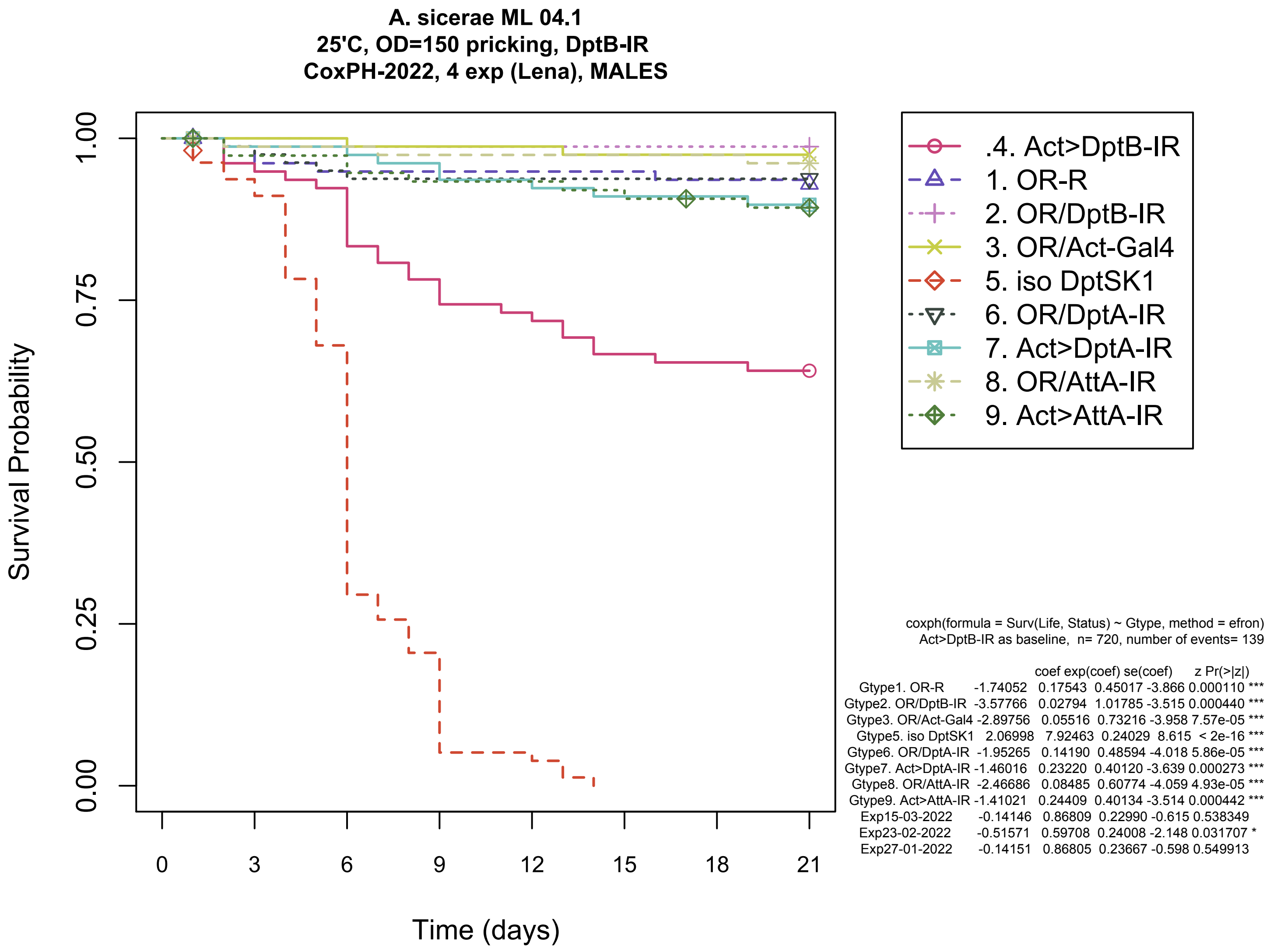

### Prett_12-03-2021.pdf

**P. rettgeri**
**25'C, OD=1.0 pricking**
**12-03-2021, MALES - ages better controlled**

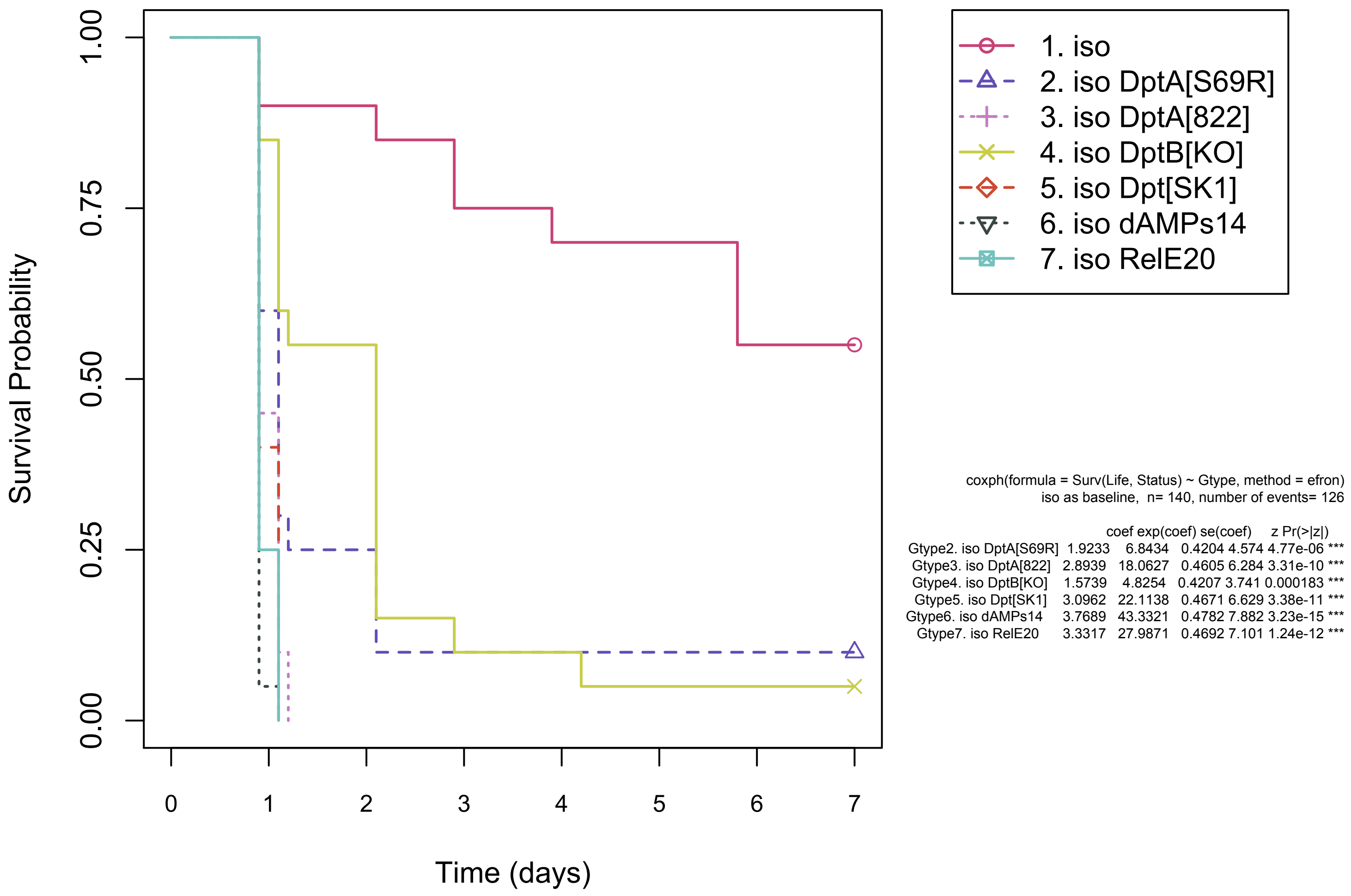

### Prett_15-03-2021.pdf

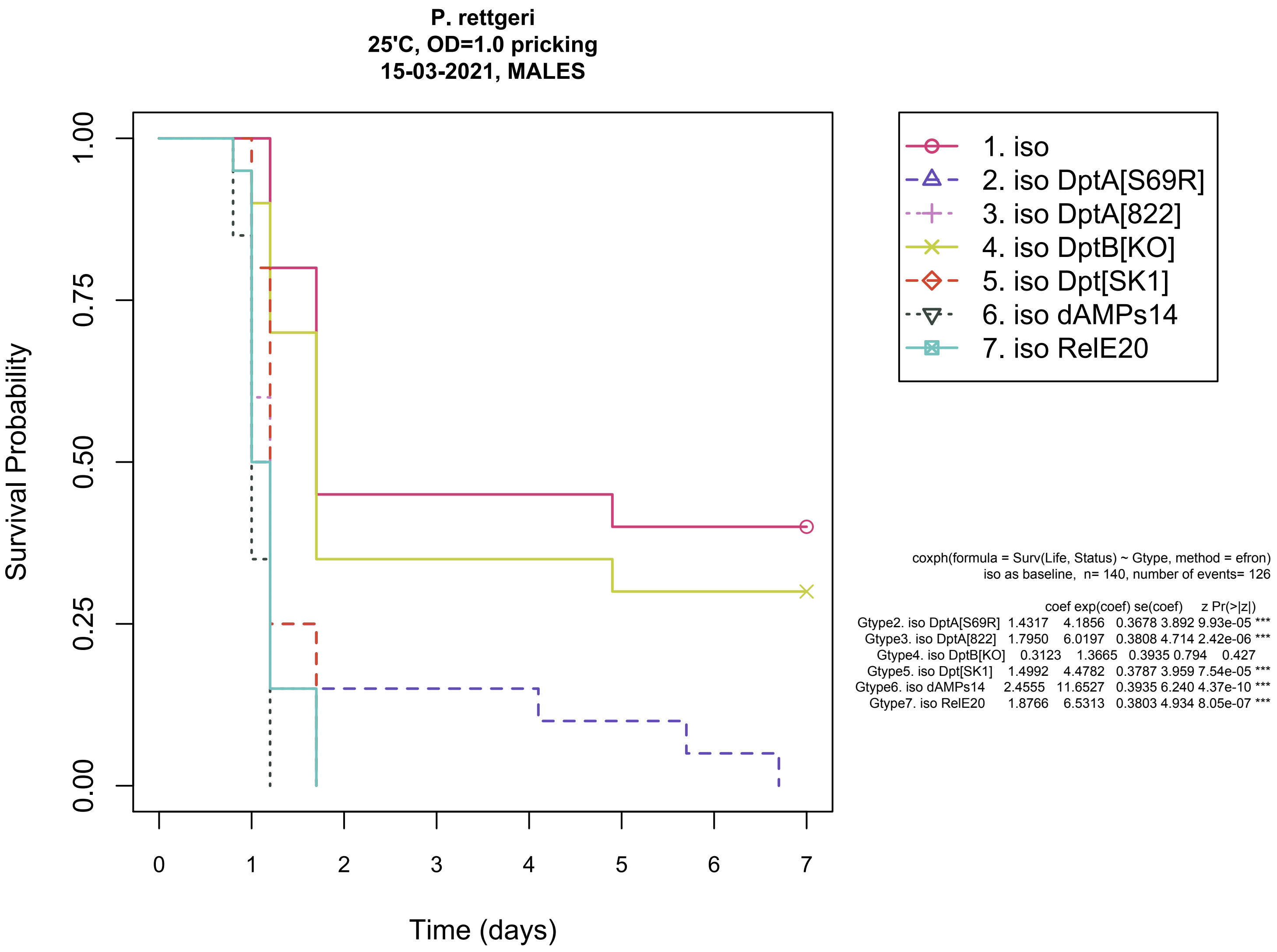

### Results_v2.pdf

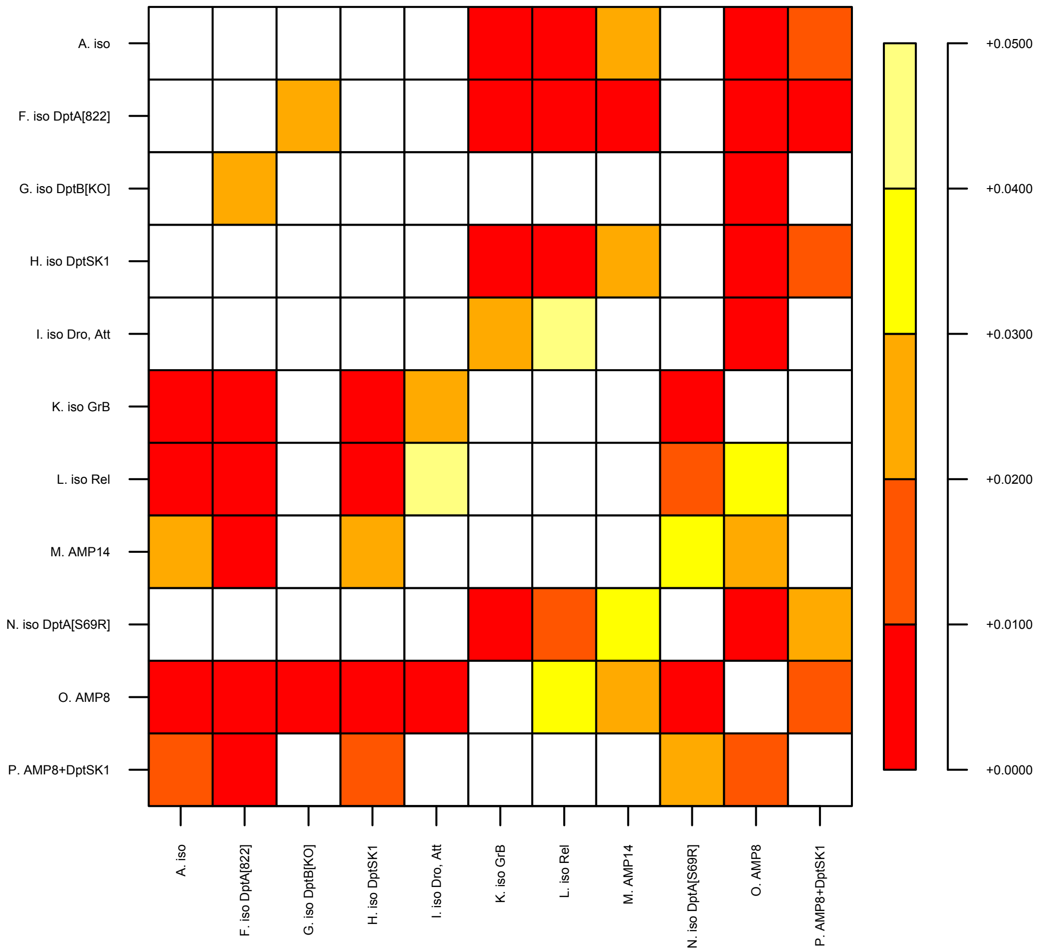
